# Transcriptional stochasticity reveals multiple mechanisms of long non-coding RNA regulation at the Xist – Tsix locus

**DOI:** 10.1101/2025.01.05.631290

**Authors:** Benjamin K. Kesler, John Adams, Gregor Neuert

## Abstract

Long noncoding RNAs (LncRNAs) are increasingly recognized as being involved in human physiology and diseases, but there is a lack of mechanistic understanding for the majority of lncRNAs. We comparatively tested proposed mechanisms of antisense lncRNA regulation at the X-chromosome Inactivation (XCI) locus. We find that due to stochasticity in transcription, different mechanisms based on the act of transcription regulate Xist and Tsix at different levels of nascent transcription. At medium levels, RNA polymerases transcribe Xist and Tsix on each strand at the same transcription site and deposit significant amounts of the histone mark H3K36me3, which inhibits Xist. At high levels of nascent transcription, many RNA polymerases transcribe Xist or Tsix resulting in transcriptional interference. Therefore, lncRNA expression variability is not just a quirk of transcription but an important aspect of regulation that allows multiple mechanisms to be employed by the same gene locus within the same cell population.

## INTRODUCTION

Protein-coding genes make up only a small portion of the human genome while transcribed noncoding genes encompass a much larger fraction of the human genome ^1,2^. In comparison to protein-coding genes, most of these noncoding genes have unknown functions and mechanisms ^2,3^, though many of them are mutated or misregulated in diseases ^3–5^. Previous studies have proposed various mechanisms for noncoding RNA functions. These mechanisms fit into two broad categories: (i) transcript-based mechanisms by which the noncoding RNA molecule regulates a target gene after transcription, and (ii) transcription-based mechanisms by which the noncoding RNA regulates the target gene through the act of transcription (Fig. 1).

**Fig. 1.**
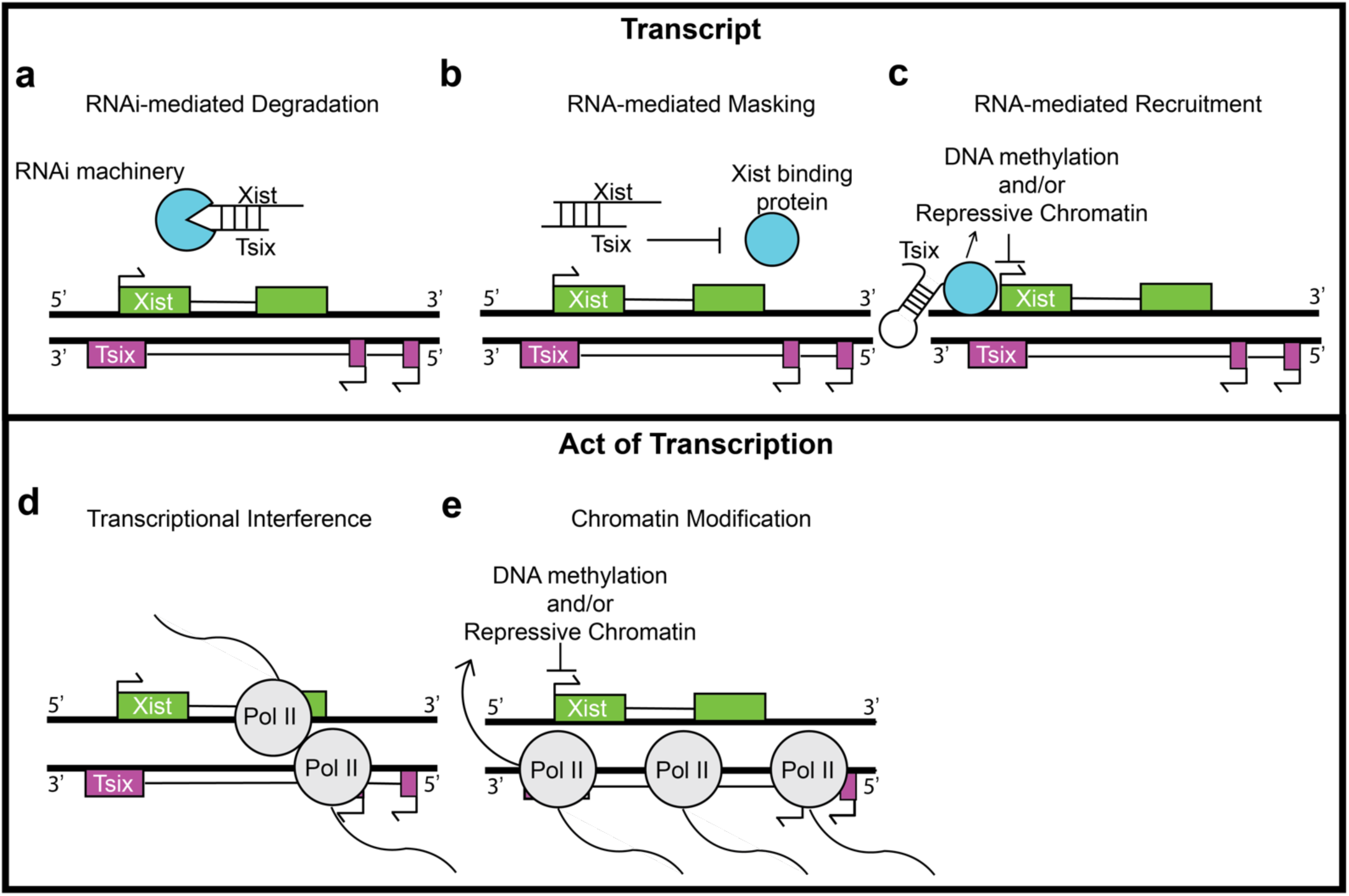
The proposed mechanisms for Tsix regulation of Xist. Transcript-based mechanisms (a-c) are on the top row, and mechanisms based on the act of transcription (d-e) are on the bottom row. **a** RNA interference (RNAi) machinery could cause degradation in response to double-stranded RNA due to Xist and Tsix binding in the same cell ^34^. **b** Binding of Tsix to Xist could prevent proteins important for XCI from binding to Xist ^16^. **c** Tsix could act as a scaffold to recruit proteins that would inhibit Xist ^35^. **d** Polymerases could collide and inhibit transcription through transcriptional interference (TI) ^36^. **e** Proteins can bind to the polymerase tails and cause chromatin changes during transcription ^14,37–40^.

Some transcript-based mechanisms involve the direct binding of a noncoding RNA to a target gene transcript through complementary base pairs. This binding can then recruit proteins that recognize double-stranded RNA, such as Dicer with noncoding microRNAs ^6^ (Fig. 1a), or the noncoding RNA can obstruct an RNA-binding protein from binding to its target, as proposed for conserved noncoding antisense transcript for β-secretase-1 (BACE1-AS), which is defined as an RNA transcribed from the opposite DNA strand of BACE1 ^7^ (Fig. 1b). Another transcript-based mechanism is for the noncoding RNA transcript to recruit and guide a regulatory protein to the specific genomic location, as proposed for the long non-coding antisense RNA transcript overlapping the INK4b/ARF/INK4a locus (ANRIL) ^8^ (Fig. 1c).

Several mechanisms instead involve the act of transcription, and one example is transcriptional interference (TI). TI occurs when polymerases physically collide upon simultaneous antisense transcription, resulting in the expression of only the sense or antisense transcript, as seen in galactose-inducible genes in yeast cells ^9^ (Fig. 1d). Another proposed mechanism is that the transcription of the noncoding RNA results in RNA Polymerase II-mediated recruitment of chromatin remodeling proteins such as Su(var)3-9, Enhancer-of-zeste and Trithorax (*SET2 & SET3*) in yeast cells ^10^ or SETD2 in mice ^11^. These histone lysine methyltransferase proteins change the chromatin environment and regulate the target gene ^12^.

Each of these mechanisms requires that the noncoding RNA and its associated proteins express and colocalize to regulate their target in the same cell. However, standard cell population assays such as RNA sequencing and chromatin or RNA immunoprecipitation cannot distinguish between cells and do not quantify the exact binding frequency ^13^. This can result in a misinterpretation of regulatory relationships and mechanisms because co-expression seen at the population level might reflect expression in different cells, binding events can occur at frequencies too low to be biologically relevant, or the proposed regulatory RNAs, proteins, or chromatin environment might not be associated with regulation of the target at the single-cell level. Considerable progress has contributed to achieving single-cell resolution for sequencing-based assays, but they still lack sensitivity required to reliably quantify noncoding RNA in a single cell ^2^.

Additionally, many previous studies utilize a variety of mutations to investigate lncRNAs of interest ^14–17^. Mutations are powerful tools for studying lncRNAs but also have potential problems. They can have off-target effects, such by inadvertently targeting an entirely different locus ^18^ or even disrupting an unknown transcript or gene regulatory element within the same locus ^19^. Investigating lncRNAs with their native sequences intact is beneficial to avoid these potential problems.

As we have shown previously, single-cell methods together with well-designed experiments can interrogate noncoding genes and their regulation in single cells simultaneously and without genetic intervention at high resolution, overcoming many of the limitations of cell population and single-cell genomic measurements mentioned above ^10,20^. However, nobody has used these quantitative methods to compare several proposed mechanisms against each other before.

As a proof of principle, we focus on X Inactive-Specific Transcript (Xist) ^21^ and its antisense transcript Tsix ^16,22^ in female mouse embryonic stem cells for which we agnostically tested several of the proposed mechanisms of Xist and Tsix mutual regulation. Xist and Tsix belong to a category of long noncoding RNAs (lncRNAs) called antisense lncRNAs ^2,23^. <u>There is also evidence that two other transcripts,</u> <u>RepA</u> ^24^ <u>and XistAR</u> ^19^<u> are present at the locus and might also be involved in antisense lncRNA regulation.</u> Several studies estimated that antisense RNAs account for 70% of all transcripts in human cells ^25^. They are ubiquitous in eukaryotes ^1^ and prokaryotes ^26^, and they are implicated in diseases ^27,28^.

This antisense lncRNA pair is of interest because Xist is necessary for X chromosome inactivation (XCI) ^29^. Abnormalities in XCI are associated with different cancers ^30,31^ and major affective disorders such as bipolar disorder and depression ^32^. The advantage of the Xist/Tsix system is that many studies have proposed different mechanisms of lncRNA gene regulation at this locus. A limitation of this system is that it is a sense/antisense pair that only covers a subclass of lncRNAs. However, it is a useful example to test many of the proposed mechanisms of noncoding RNA regulation through a rigorous quantitative framework of single-molecule and single-cell fluorescence microscopy and analysis. Although Xist and Tsix are among the most well-studied mammalian lncRNAs, and Tsix is an established regulator of Xist in mice ^33^, there are many proposed mechanisms for Tsix regulation of Xist, some of which have conflicting evidence or even a complete lack of evidence ^33^.

The proposed mechanisms for Tsix regulation of Xist fit in the same general classifications described above (Fig. 1). The mechanisms involved in this context are categorized into two types: those related to transcripts (Fig. 1a-c) and those concerning transcription (Fig. 1d-e). Transcript-based mechanisms encompass the RNA interference (RNAi) system, which can trigger the degradation of double-stranded RNA, a process that involves the binding of Xist and Tsix within the same cell (Fig. 1a) ^34^. Additionally, the binding of Tsix to Xist may block crucial proteins from associating with Xist, which is necessary for X-chromosome inactivation (XCI) (Fig. 1b) ^16^. Tsix may also serve as a structural framework to attract proteins that suppress Xist’s function (Fig. 1c) ^35^. On the other hand, transcription-based mechanisms include transcriptional interference (TI), where collisions between polymerases can impede the transcription process (Fig. 1d) ^36^. Furthermore, certain proteins are known to attach to polymerase tails, leading to alterations in chromatin during the transcriptional activity (Fig. 1e) ^14,37–40^. In addition, Xist contains a repeat A region that is reported to give rise to a separate transcript ^24^.

In this work, we agnostically tested all the proposed mechanisms of Xist and Tsix mutual regulation by combining single-molecule RNA fluorescent in-situ hybridization (smRNA-FISH) (Supplementary Fig. 1), immunofluorescence, and quantitative expression and colocalization analyses in wildtype female mouse embryonic stem cells during differentiation. The proposed method can be applied broadly to test any antisense transcript for these mechanisms in the future. Furthermore, the specific analysis methods utilized in this study are suitable for other noncoding and protein-coding pairs involving colocalization and fluorescence quantification.

## RESULTS

### Mechanisms based on transcripts: Tsix Binding to Xist

The possibility of complementary base pairing between antisense lncRNAs such as Xist and Tsix has been proposed to result in inhibition through at least two possible mechanisms: transcript degradation through RNA interference (RNAi) machinery (Fig. 1a) ^34^ and masking of a critical region (Fig. 1b) ^24^. RNAi recognizes and processes double-stranded RNA (dsRNA) with Dicer and degrades sequences matching the dsRNA (Fig. 1a) ^41^. Therefore, any antisense RNA could potentially act through this mechanism. A previous study suggested that Tsix and Xist form a duplex processed by Dicer to inhibit Xist ^34^. However, other studies showed conflicting evidence as to whether Tsix inhibits Xist by this mechanism ^24,42–44^. Sense-antisense binding could instead inhibit Xist through masking, because Tsix’s exonic sequences overlap with a section of Xist that is critical for XCI called the Repeat A region (Fig. 1b) ^24^. However, there is no experimental evidence for or against this mechanism. Furthermore, the Repeat A region encodes a separate transcript ^24^. Although earlier studies showed Xist and Tsix signals in the same cells ^22,45^, none of these previous studies demonstrated simultaneous colocalization of individual Tsix and Xist transcripts within the same cells and outside the transcription site, which would be necessary for either of these mechanisms.

To address this gap in knowledge, we utilized smRNA-FISH to investigate the single-cell expression and colocalization of Xist and Tsix. smRNA-FISH requires chemical cross linking, cell permeabilization, and hybridization of complementary fluorescently-labeled oligonucleotides to detect RNA molecules (Supplementary Fig. 1 & 2) ^46,47^. We utilized undirected differentiation of female mouse embryonic stem cells (mESCs) as an *ex vivo* system to recapitulate the early stages of Xist regulation and quantify thousands of cells. To achieve high expression of Xist, cells are grown in 2i media to stay in the pluripotent state (0.5 % of Xist cloud positive cells) (Supplementary Fig. 3) and then non directionally differentiated for two days by removing 2i media (52 % of Xist cloud positive cells) (Fig. 2a), which consistently resulted in high Xist expression in our experiments and previously published results using the same system and cell line (Supplementary Fig. 4 & 10) ^48,49^. We chose the day 2 time point because both Xist and Tsix show variable expression, which is necessary to analyze their co-expression within the same cell and allele. Time points before day 2 show less Xist expression because Xist is not upregulated yet, making it significantly harder to analyze both Xist and Tsix co-expression. Likewise, time points after day 2 show less expression variation in Tsix because Tsix gets repressed. To visualize whether spliced Tsix binds to spliced Xist, we designed three sets of RNA-FISH probes (Fig. 2b): (i) probes targeting the Repeat A region (A, RepA probes), (ii) probes targeting exons of Xist outside the Repeat A region (Xist Exon probes), and (iii) probes targeting exons of Tsix outside of the region that overlaps with Xist (Tsix Exon Probes). Because the Xist Exon probes and Tsix Exon probes target the regions of Xist and Tsix that are not complementary to each other, they can bind to mature Xist and Tsix even if the antisense lncRNA pair are hybridized together (Fig. 2b insert, left). As an example of a colocalized signal in RNA FISH is the hybridization of probes to Xist and RepA (Fig. 2b insert, right, Supplementary Fig. 11-14). Although we expected colocalization between Xist and RepA, previous studies showed that colocalization is not perfect ^46^.

**Fig. 2.**
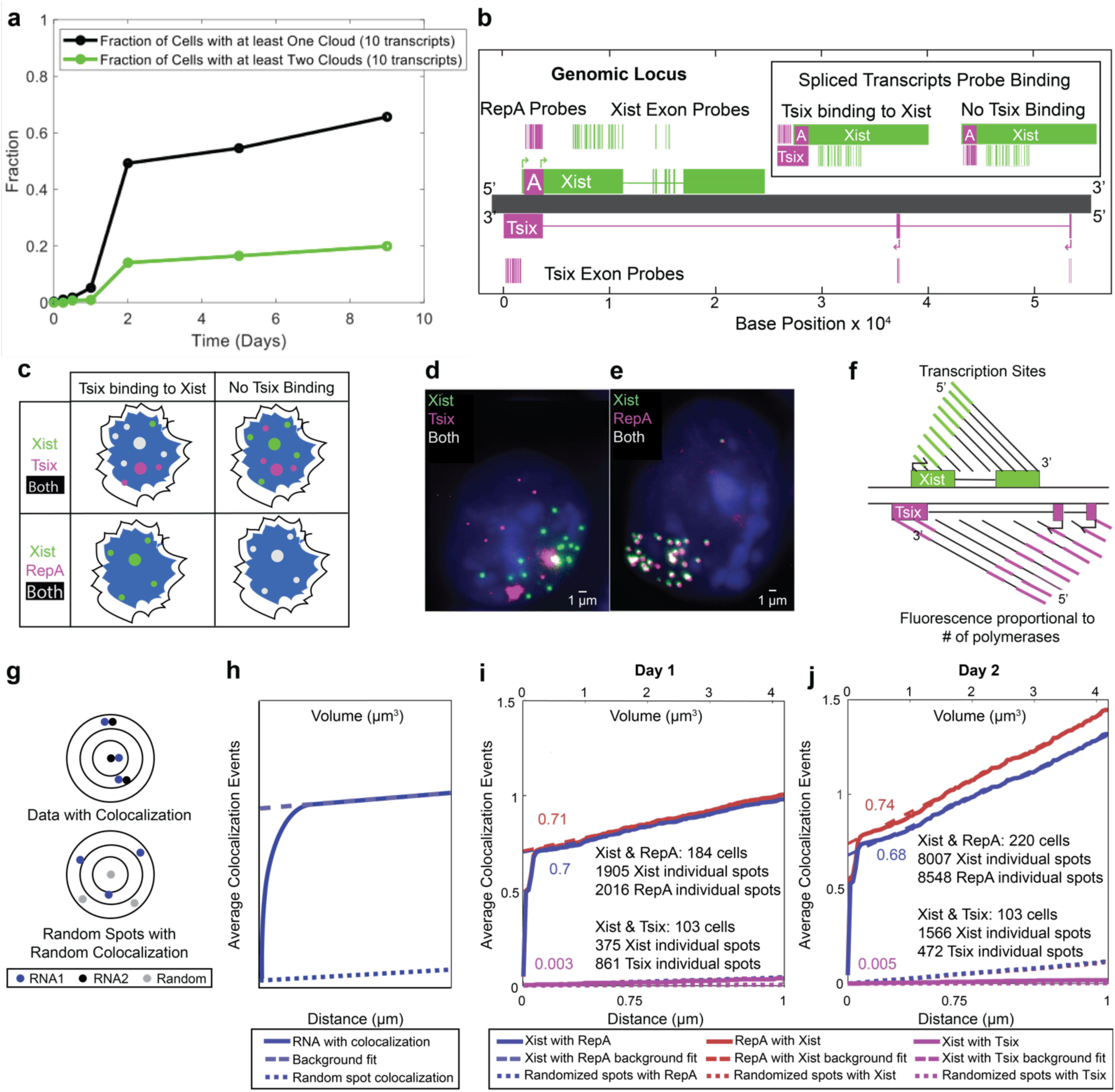
Lack of Xist and Tsix colocalization disproves mechanisms involving their binding. **a** Faction of cells with one (black) or two (green) Xist clouds during a 9 day differentiation experiment. **b** Diagram illustrating the Xist/Tsix locus (to scale) with the Xist (green) exonic and intronic region, the Repeat A (RepA) region (A, magenta), Tsix (magenta) exonic and intronic region (horizontal magenta line). Probe positions are shown as vertical lines. The insert shows where probes are expected to bind in cases of mature Tsix binding to mature Xist (left) or no Tsix binding to Xist which will result in binding of RepA probes to Xist (right). **c** Diagram showing expected results: Left shows Tsix binding to Xist with colocalization at transcription sites (large circles, blue nucleus), and right shows Tsix not binding to Xist with RepA colocalization. **d-e** Example images: **d** Tsix (magenta) rarely colocalizes (white) with Xist (green), except at bright transcription spots. **e** RepA (magenta) almost perfectly colocalizes with Xist (green). **f** Diagram illustrating brighter RNA-FISH signals at transcription sites due to multiple polymerases. **g** Diagrams of colocalization calculation using Particle Image Cross-Correlation Spectroscopy (PICCS), showing colocalization between RNA 1 (blue) and RNA 2 (black), and between random spots (gray) and RNA 1. ^51^**h-j** Colocalization was quantified by plotting the average events per transcript. The bottom x-axis shows the search radius (µm) on a cubic scale, with volume increase on the top x-axis. Solid lines represent data, long dashed lines show the background fit, and dotted lines indicate random colocalization. The y-intercept of the background fit gives the true colocalization after background subtraction. **h** Graph showing average colocalized spots between RNA 1 and RNA 2 (blue line) or random spots and RNA 1 (dotted line) by distance. **i** Day 1 and **j** Day 2 differentiation: Xist and RepA show significant colocalization (Day 1: 0.70-0.71; Day 2: 0.68-0.74), while Xist and Tsix show minimal colocalization (Day 1: 0.003; Day 2: 0.005).

Therefore, if Tsix binds to Xist in mESCs, we expect Xist and Tsix spots to colocalize (Fig. 2c, left). If Tsix does not bind Xist, we expect Tsix and Xist spots to not colocalize. An example of RepA and Xist spots colocalize is shown in Fig. 2c, right.

Imaging Xist and Tsix expression showed minimal colocalization of Xist and Tsix transcripts (Fig. 2d, Supplementary Fig. 4, 6, 7) and high colocalization of signal from Xist Exon and RepA probes (Fig. 2e, S11, S13, S14). Bright spots of Xist and Tsix that colocalize in the nucleus are likely transcription sites, which have greater intensity due to multiple polymerases transcribing RNA (Supplementary Fig. 2f). We quantified our data by automatically segmenting cells and nuclei ^50^ ( Supplementary Fig. 1, 5, 7, 12, 14, 21, S23, 25, 27, 31, 33). After cell segmentation, we applied a semi-automated image analysis pipeline to identify and quantify spots, transcription sites and Xist clouds (See Materials and Methods, Supplementary Fig. 1). Xist clouds form due to clustering of transcripts on the X chromosome away from the transcription site. After RNA spot detection, we applied Particle Image Cross-Correlation Spectroscopy (PICCS) ^51^ to quantify average colocalization events across different radii (Fig. 2g). PICCS typically plots average colocalization versus distance squared or area for 2D data. To adapt PICCS to our 3D data, we instead plotted colocalization events as a function of the distance cubed or volume (Fig. 2h, solid line). To account for random colocalization events (Fig. 2g), PICCS fits a line to the data far from where real colocalization occurs (Fig. 2h, dashed line). The y-intercept of this line is the colocalization fraction after subtracting random colocalization events. As a control, we also utilized randomly-generated spots throughout each nucleus (Fig. 2g) and quantified colocalization (Fig. 2h, dotted line). The y-intercept of this control should be zero, indicating no colocalization above what would be expected by chance. With this quantification scheme, we determined colocalization of Xist with RepA and Xist with Tsix at both day 1 and day 2 of differentiation (Fig. 2i, j). At both timepoints, we found low colocalization (0.003 and 0.005 at days 1 and 2 respectively) between individual Xist and Tsix molecules, indicating that they do not bind each other very often. Conversely, we found high colocalization (approximately 0.7) between the Xist Exon and RepA spots at both timepoints. As expected, random spots did not show colocalization (Fig. 2i, j, dotted lines).

Together, these results provide strong evidence against Tsix and Xist binding for the majority of transcripts, which most likely precludes RNAi machinery (Fig. 1a) or transcript masking (Fig. 1b) being involved in Tsix regulation of Xist. The hypothesis that RNAi-induced rapid RNA degradation is responsible for non-detection is not supported by our prior findings. We have established that RNA-FISH can effectively identify RNA molecules that have lifetimes of less than three minutes ^52,53^, suggesting that RNAi degradation speed should not hinder detection.

### Mechanisms based on transcripts: Xist and Tsix acting in *trans*

lncRNAs that recruit regulatory factors (Fig. 1c) can diffuse throughout the cell and regulate other genomic loci (also known as working in *trans*) ^2^. Previous studies showed a lack of *trans* inhibition of Xist by Tsix with mutants artificially overexpressing Tsix from specific alleles ^15,54^. However, whether these results would change in the wildtype context is unclear. Our data showed that wildtype Tsix can diffuse throughout the cell, potentially allowing Tsix to regulate Xist in *trans* (Fig. 3a). If Tsix regulates Xist in *trans*, then high levels of Tsix would reduce whole-cell levels of Xist and cause negative correlation between the two transcripts (Fig. 3b, left). Conversely, there would be little to no whole-cell correlation if there is no *trans* regulation (Fig. 3b, right). Therefore, we tested whether there was negative correlation between mature and dispersed Xist and Tsix transcripts. We quantified Xist and Tsix at day 2 of differentiation in three new biological replicas to determine the co-expression of Xist and Tsix within the same cells (Supplementary Fig. 3, 4-10, 15). To determine the number of transcripts in Xist clouds, we divided the total intensity in the cloud by the intensity of a single transcript (Supplementary Fig. 16). Although the distribution had an L-shaped pattern (Fig. 3c), which often indicates mutual exclusivity, this shape can occur even with uncorrelated data (Supplementary Fig. 17). The Spearman test, a nonparametric test of correlation, showed no statistically significant correlation for two out of three replicas (Fig. 3c, white text). As a control, we also used the same dataset but randomly permuted the single-cell relationship between the two genes. Critically, none of the replicas significantly differed statistically from randomized data when evaluated by the two-dimensional Kolmogorov-Smirnov test ^55^ (Fig. 3d). Therefore, our data and analyses argue against a regulation mechanism in *trans*.

**Fig. 3.**
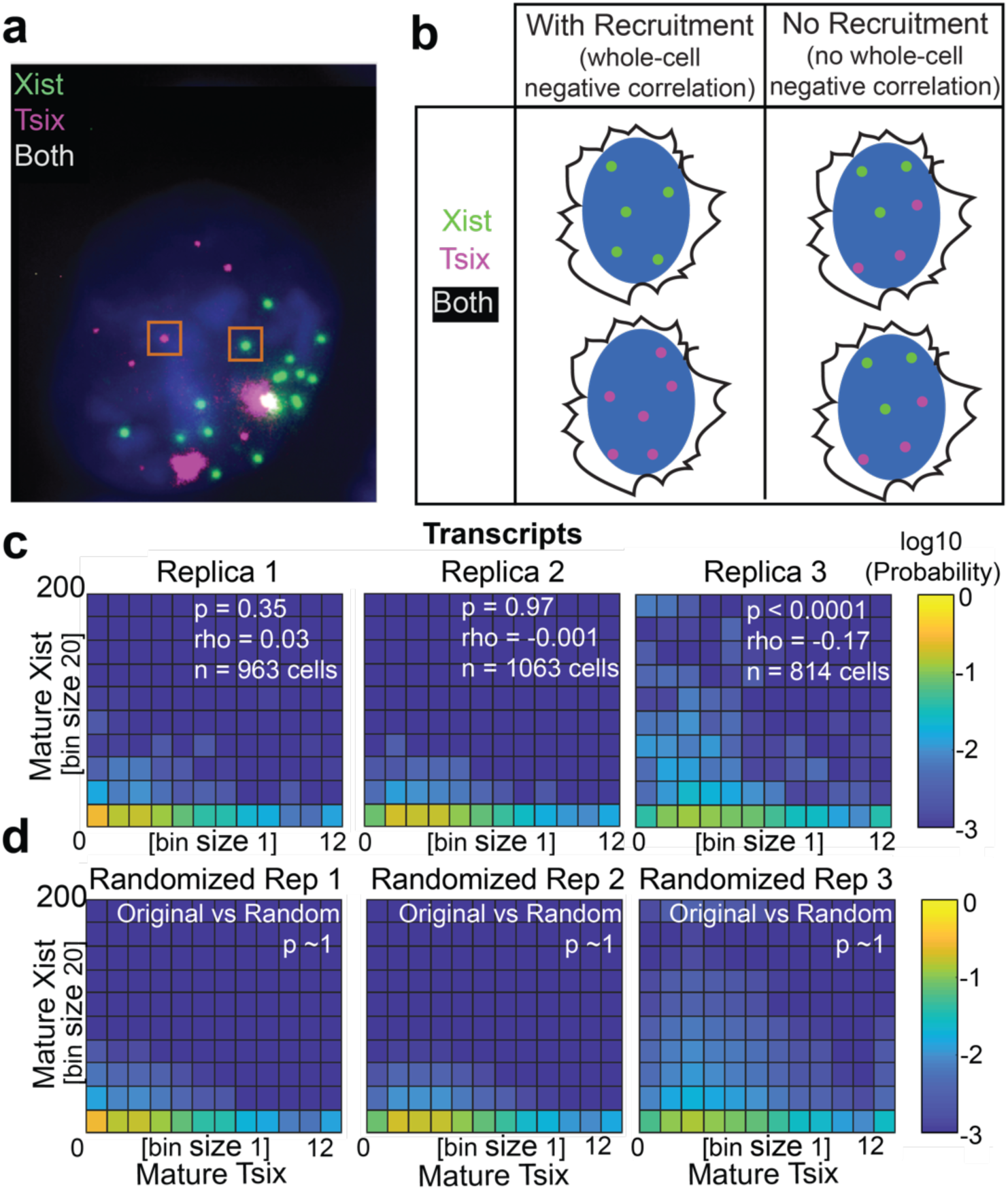
Lack of whole-cell negative correlation between Xist and Tsix argues against mechanisms involving transcript recruitment. **a** An example image of data showing individual molecules of mature Xist (green) and Tsix (magenta) surrounded by an orange box. This is the same cell as shown in Fig. 2D. **b** Conceptual diagram showing the expected data. Recruitment of transcription regulators on a whole-cell level would result in negative correlation when investigating whole-cell transcript levels (left). Otherwise, there would be no whole-cell correlation (right). **c** The joint probability distribution of whole-cell levels of Xist and Tsix mature transcripts in three different replica experiments. The correlation coefficient (rho) and the significance (p) of correlations are determined by the Spearman correlation test, and no significant correlation is found in two out of three replicas. **d** The joint probability distribution of the randomized data, which used the same data but randomly permuted the single-cell relationship between the two genes. A two-dimensional KS-test ^55^ (p-value shown) determined no significant difference between the original data and the randomized version in all replicas. Xist data is binned by 20 transcripts. Tsix data is binned by 1 transcript due to its low expression. Color code corresponds to the negative logarithm base 10 of the probability. Replica 1, n = 963 cells, 2289 Tsix transcripts, 5226 Xist transcripts; replica 2, n = 1063 cells, 3322 Tsix transcripts, 5690 Xist transcripts; replica 3, n=814 cells, 3116 Tsix transcripts, 19701 Xist transcripts.

### *Cis*-acting Mechanisms at the Site of Transcription

When lncRNAs regulate genes locally on the same allele, they are said to work in *cis* ^2,56^. Some examples of *cis* mechanisms such as transcriptional interference or chromatin modification regulate through the act of transcription, (Fig. 1d-e). Previous studies utilized multiple mutagenesis strategies including deleting Tsix’s promoter region, inserting a strong promoter to drive Tsix, or prematurely stopping Tsix transcription to demonstrate that Tsix regulates Xist in *cis* on the same allele ^15,17,22,54^. However, they did not investigate individual wildtype transcription sites quantitatively in single cells to show that Tsix is regulating Xist in *cis*. Because transcription sites are identifiable as spots brighter than individual transcripts, we were able to quantify the presence of Xist and Tsix specifically at the transcription sites and their correlation using the data generated in the previous section (Fig. 4A, Supplementary Fig. S4-7, 9, 10, 16, 18). If there is *cis*-acting regulation, there would be negative correlation between Tsix and Xist Exon signal at the site of transcription (Fig. 4b, left). If there is no *cis*-acting regulation, there would be no negative correlation and many instances of co-transcription (Fig. 4b, right). We investigated the transcription sites with the same quantitative framework as *trans*-acting mechanisms (Fig. 3). With this framework, we found strong negative correlation in all replicas (Fig. 4c). Every replica was also significantly different from the randomized dataset (Fig. 4d). This evidence strongly suggests that Tsix regulates Xist in *cis*.

**Fig. 4.**
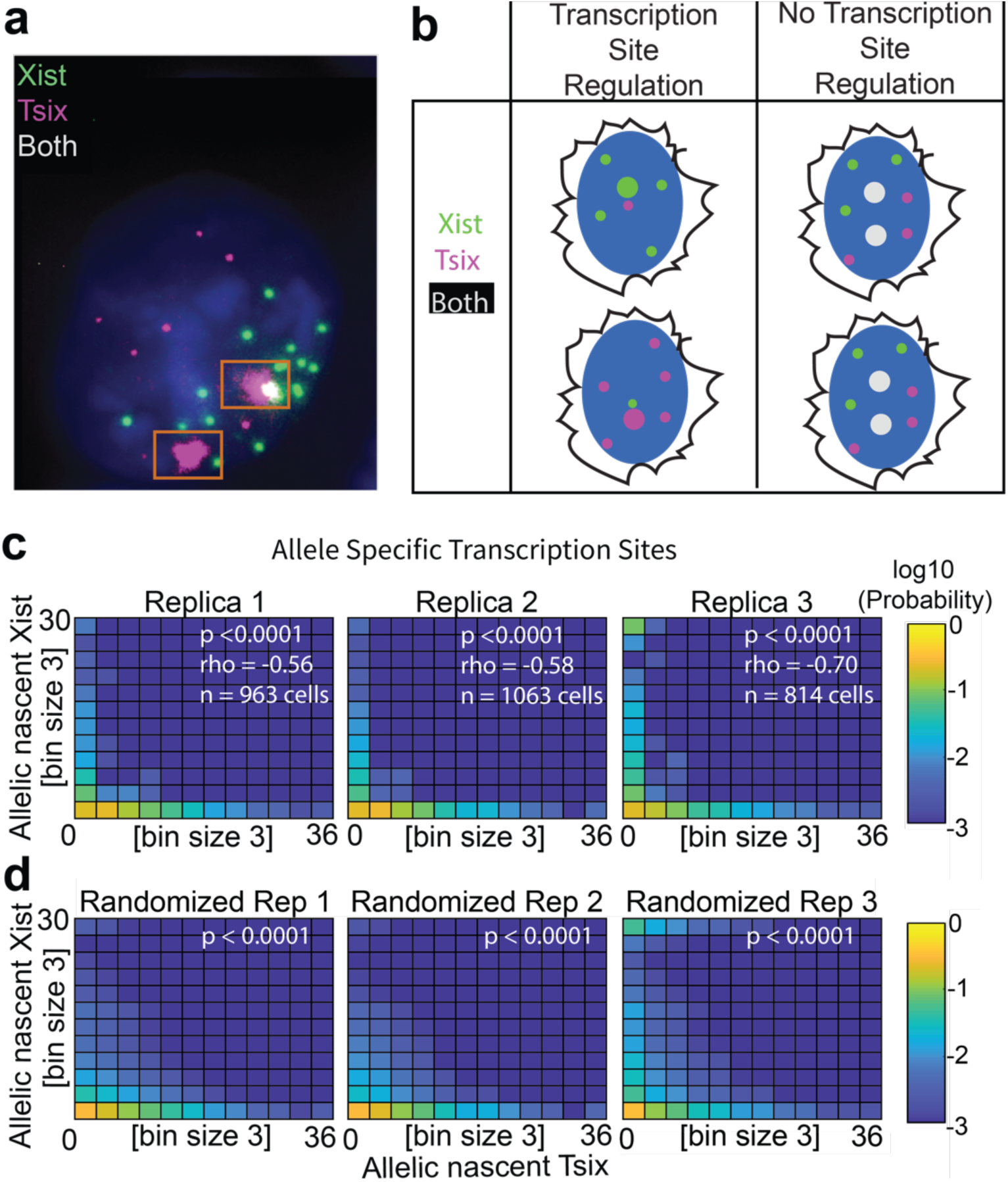
Allele specific quantification of transcripts at the site of transcription suggests a transcription-based mechanism. **a** An example image of data showing Xist (green) and Tsix (magenta), with brighter signal at allele specific transcription sites (surrounded by an orange box) compared to signal at individual transcripts. Colocalized signal is white. **b** Conceptual diagram showing the expected data. Regulation at the transcription site would result in negative correlation between Xist and Tsix levels at the site of transcription (left). No transcription-based regulation would result in no correlation at the transcription site and co-transcription would be likely (right). **c** The joint probability distribution of allele specific Xist and Tsix transcripts at the site of transcription (determined by total brightness / brightness of an individual transcript) in three different replica experiments. Rho and p values are determined by the Spearman correlation test, and significant negative correlation is found in all three replicas. **d** The joint probability distribution of the randomized data (randomized the same as Fig. 3). A two-dimensional KS test ^55^ (p-value shown) determined significant differences between the original negatively correlated data and the randomized version in all replicas. Xist and Tsix transcripts in the site of transcription is binned by 3 transcripts each. Color code corresponds to the negative logarithm base 10 of the probability. Replica 1, n=963 cells, 1045 Tsix transcription sites, 7437 nascent Tsix transcripts, 377 Xist transcription sites, 2936 nascent Xist transcripts; replica 2, n=1063 cells, 867 Tsix transcription sites, 5963 nascent Tsix transcripts, 316 Xist transcription sites, 2290 nascent Xist transcripts; replica 3, n=814 cells, 711 Tsix transcription sites, 4855 nascent Tsix transcripts, 507 Xist transcription sites, 9695 nascent Xist transcripts. Fig. 4 analyzed the transcription sites for the same cells as shown in Fig. 3.

### Transcriptional Interference of Xist and Tsix at high nascent transcription

Transcriptional Interference (TI) through antisense polymerase collision is one possible mechanism for regulation in *cis* (Fig. 1d) ^57,58^. In this model, polymerases that initiate from converging transcription units physically collide, resulting in premature polymerase termination. This model proposes a mutually exclusive expression pattern of sense and antisense transcription in overlapping regions where TI occurs, while not affecting non-overlapping regions. Without TI, overlapping regions display co-transcription ^59,60^. A recent study investigating differential effects between overlapping and non-overlapping regions of Xist and Tsix suggests that TI affects the Tsix/Xist locus to cause mutual inhibition, but the study utilized a strong artificial and inducible promoter to drive Xist ^36^. It is therefore unclear whether TI is prevalent with the native promoters of Xist and Tsix intact. To distinguish between TI and other mechanisms in wildtype cells, we designed intronic probes at the Xist and Tsix locus, which allowed us to visualize nascent transcription in new experiments (Supplementary Fig. 19. We performed these experiments on day 2 of differentiation, when there is sufficient Xist expression (Fig. 2A, Supplementary Fig. S20-27).

We first designed interlaced intronic probes targeting Xist and Tsix within the Xist transcriptional unit (Xist Intron and Tsix 3’ Intron probes, respectively) to quantify nascent expression from the same genomic location (Fig. 5a). If TI restricts overlapping co-transcription, we only expect to observe nascent transcripts of either the Xist intron or the Tsix 3’ intron but never signal from both transcripts (Fig. 5b, left). If TI is not occurring, we expect polymerases from both promotors to read through the same genomic location. This phenomenon will result in colocalization of the Xist intron and Tsix 3’ intron nascent transcripts (Fig. 5b, right). In our microscopy experiments, we observed colocalization of Xist intronic and Tsix 3’ signal (Fig. 5c, Supplementary Fig. 16, 20-23). This result indicates that overlapping Xist and Tsix regions can co-transcribe.

**Fig. 5.**
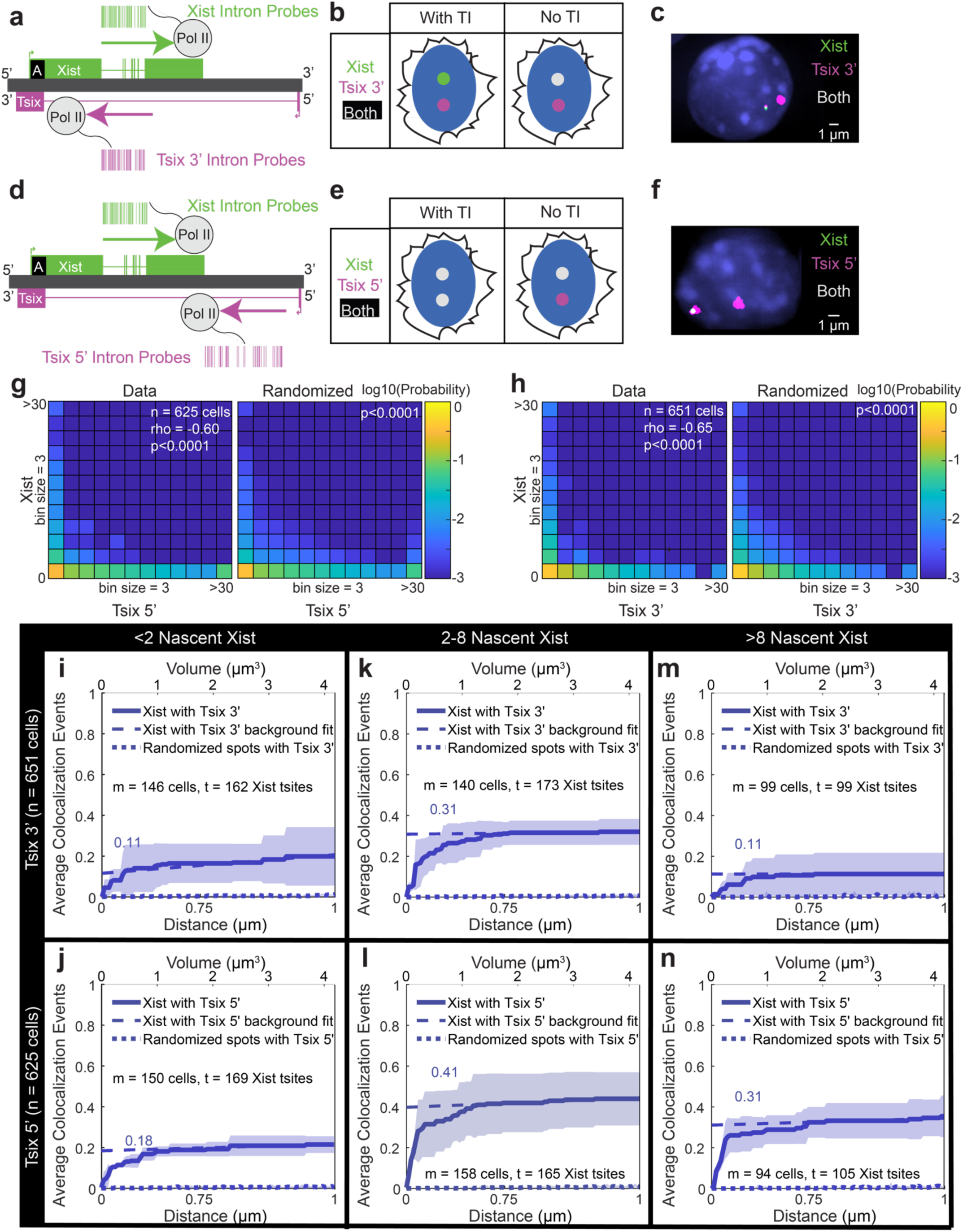
Co-transcription and Transcriptional Interference (TI) at the Xist/Tsix Locus. **a** Diagram of the Xist/Tsix locus showing Xist exons (green), Tsix exons (magenta), introns (lines), and probe positions for Xist introns (green) and Tsix 3’ introns (magenta). Polymerase positions suggest potential co-transcription of Xist and Tsix. **b** Conceptual diagram showing expected outcomes with or without TI. If TI occurs, Xist (green) and Tsix 3’ (magenta) signals won’t colocalize (left). Without TI, colocalization occurs (right). **c** Example image showing DAPI (blue), Xist (green), Tsix 3’ (magenta), and colocalization (white). **d-f** Similar diagrams and images as (A-C), but for Tsix 5’. With TI, more Xist and Tsix 5’ colocalization occurs than with Tsix 3’ (left); without TI, colocalization remains consistent (right). **g-h** Joint probability distributions of nascent Tsix (x-axis) and Xist (y-axis) transcripts. Real data is on the left, randomized data on the right. Rho and p values indicate Spearman correlations (real data) and 2D KS test results (randomized data). Analysis includes 625 cells for Tsix 5’ and 651 cells for Tsix 3’. **i-n** Quantification of colocalization at transcription sites, varying by nascent transcription levels (low, medium, high). Colocalization is plotted by search radius (x-axis, cubic scale) and average events per transcription site (y-axis). Solid line is the mean and shaded area is the standard deviation. Dashed lines is the background fit. Dotted line is the colocalization by chance. **i, k, m** Quantification of colocalization of Xist with Tsix 3’ depending on the nascent transcription of Xist. Low-level nascent Xist (i), shows colocalization with Tsix 3’ (18 tsites). A greater fraction of medium-level Xist colocalizes with Tsix 3’ (54 tsites) (k), and a lower fraction of high-level Xist colocalizes with Tsix 3’ (11 tsites) (m). A total of n = 651 cells are quantified in this analysis. **j, l, n** Quantification of colocalization of Xist with Tsix 5’ depending on the nascent transcription of Xist. Low-level nascent Xist (j), shows colocalization with Tsix 5’ (30 tsites). A greater fraction of medium-level Xist colocalizes with Tsix 5’ (68 tsites) (l), and similarly high fraction of high-level Xist colocalizes with Tsix 5’ (33 tsites) (n). A total of n = 625 cells are quantified in this analysis.

To better understand the relationship between TI and co-transcribing polymerases, we utilized smRNA-FISH with both Xist Intronic probes and Tsix 5’ Intron probes (Fig. 5d). If TI significantly affects the locus, we would expect more colocalization of Tsix 5’ Intron signal with Xist Intron signal compared to the previous data with Tsix 3’ Intron and Xist Intron (Fig. 5e, left). If there is no TI, we would expect similar colocalization as with Tsix 3’ (Fig. 5e, right). Initial visualization showed similar colocalization of Tsix 5’ Intron signal with Xist Intron signal (Fig. 5f, Supplementary Fig. 16, 24-27).

Initial analysis of the joint expression of both Tsix 5’ and Tsix 3’ nascent transcription showed negative correlation with Xist nascent transcription, and both were significantly different from random (Fig. 5g-h, Supplementary Fig. 28-31). There did not seem to be a significant difference in the negative correlation values overall (rho = -0.58 for Tsix 5’ versus rho = -0.64 for Tsix 3’). To better understand the source of this negative correlation, we analyzed the significant variability in nascent transcription, which is a consequence of stochastic gene expression, a fundamental property of single cells (Supplementary Fig. 18). This prompted us to utilize this natural variability in Xist and Tsix expression and analyze colocalization with different levels of nascent transcription to gain nascent transcription dependent insights into transcriptional interference in addition to negative correlations.

For different nascent transcription levels, we quantified the colocalization fraction of transcription sites using PICCS. Xist transcription sites with low nascent transcription (<2 nascent transcripts) had low colocalization with Tsix 3’ nascent transcription (0.11, Fig. 5I, Supplementary Fig. 32). However, low-expressing Xist sites also had low colocalization with Tsix 5’ nascent transcription (0.15, Fig. 5J, Supplementary Fig. 32). These colocalization fractions were not significantly different from each other statistically when evaluated by the Cochran-Mantel-Haenszel test, an inferential test for the association between two binary variables ^61^ (p = 0.16), suggesting that TI is not a significant regulation mechanism at low levels of nascent transcription. When nascent Xist is at medium levels (2-8 nascent transcripts), there was a significantly higher colocalization fraction with both Tsix 3’ nascent transcription (0.31, Fig. 5k, Supplementary Fig. 32) and Tsix 5’ nascent transcription (0.4, Fig. 5l) compared to the fractions when nascent Xist was at low level. These colocalization fractions were still not significantly different from each other statistically (p = 0.23), suggesting that TI is not significant at medium levels of nascent transcription. At high levels of nascent transcription (>8 nascent transcripts), Xist transcription sites colocalized less with Tsix 3’ transcription (0.12, Fig. 5m, Supplementary Fig. 32), while colocalization with Tsix 5’ transcription remained high (0.31, Fig. 5n, Supplementary Fig. 32). These colocalization fractions were significantly different from each other statistically (p = 0.02), which is consistent with predictions from TI ^36^. Another prediction for TI is low processivity due to collision with Xist polymerases, which means that Tsix will sometimes transcribe through its 5’ region but not the 3’ region that overlaps Xist. In other words, in an experiment that visualizes both the Tsix 5’ and Tsix 3’ nascent transcription (Supplementary Fig. 28-31, 33), we expect to see transcription sites that only have Tsix 5’ signal if TI is occurring. We saw this indication of low processivity when analyzing our data, especially when Tsix was lowly expressed (Supplementary Fig. 33). Overall, these results support that TI is contributing to antisense regulation at the Xist/Tsix locus, but only at high levels of nascent transcription. This data also shows a prevalence of antisense co-transcription, especially at medium levels of nascent transcription.

### Chromatin Modification during medium transcription

Another possible mechanism for gene regulation by lncRNAs is recruitment of factors involved in transcription regulation by the polymerase machinery (Fig. 1e). For instance, the C-terminal domain (CTD) of pol II recruits SET proteins that tri-methylate the 36th lysine residue of the histone H3 protein (H3K36me3) ^10,12,62^. This histone mark can then recruit factors such as histone deacetylase Rpd3 in yeast or DNA methyltransferases 3A and 3B in mammals to repress transcription ^63,64^. In the Tsix/Xist system, Tsix induction or premature termination alters chromatin modifications including H3K36me3 at the Xist promoter ^37–40^. Recent studies detailed H3K36me3 dynamics upon Tsix induction ^40^ and found that knockdowns of Setd2 caused Xist upregulation ^39^. These studies suggest that Tsix establishes H3K36me3 to inhibit Xist. However, these studies did not measure chromatin modifications and RNA expression at the locus in the same single cells, which is critical to establish a direct relationship. In new experiments, we utilized quantitative immunofluorescence (IF) and smRNA-FISH in the same cells to spatially correlate chromatin state through H3K36me3 with nascent Xist and Tsix expression in wildtype cells (Fig. 6a, Supplementary Fig. 34-36). The antibody for H3K36me3 was diluted so that individual spots could be detected ( Supplementary Fig. 36). Random spots were again generated in each nucleus as a control.

**Fig. 6.**
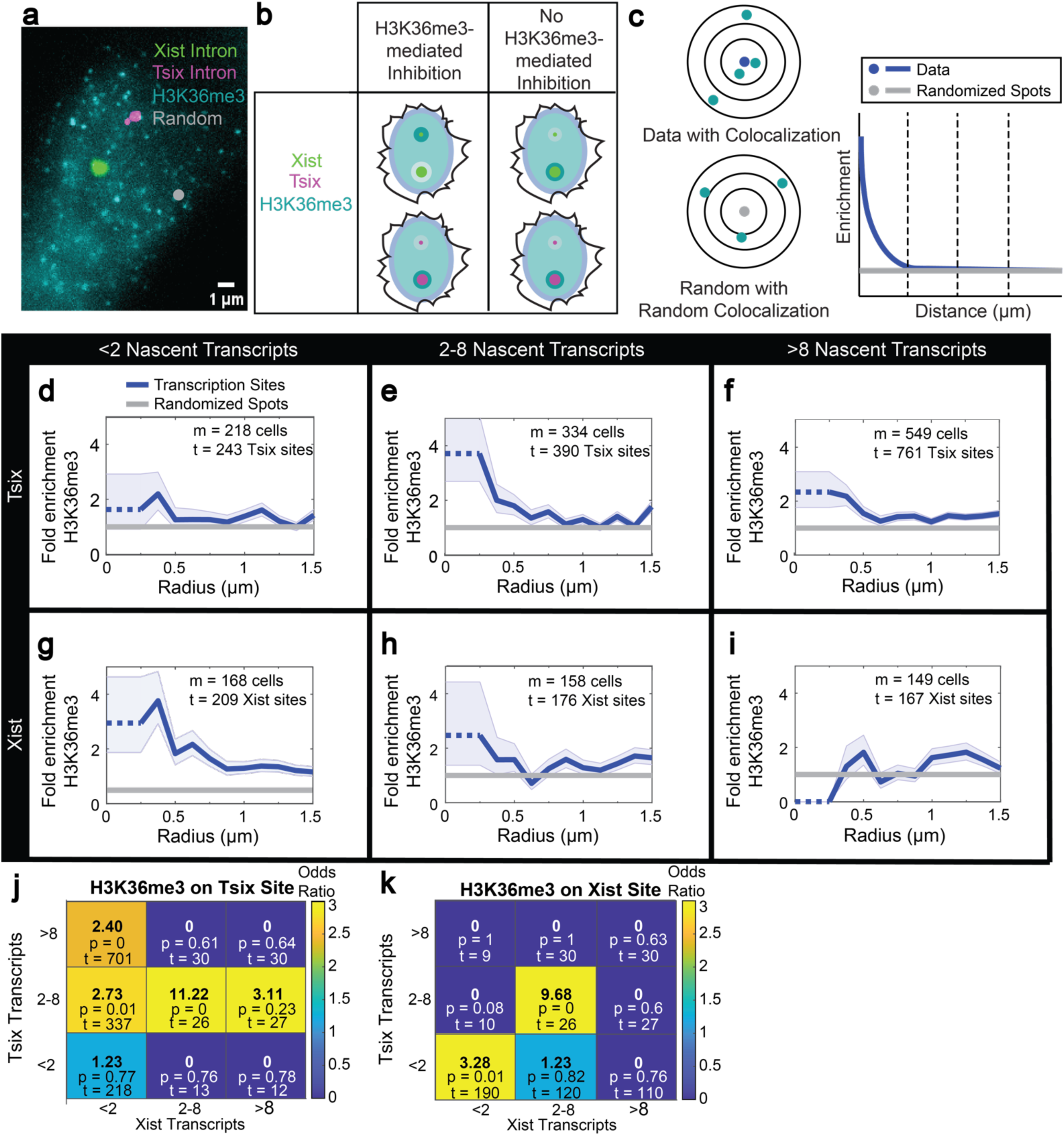
Xist and Tsix transcription is associated with different enrichment levels of trimethylated histone 3 at Lysine 36 (H3K36me3). **a** Example RNA-FISH for nascent Xist (green) and Tsix 5’ (magenta) combined with quantitative immunofluorescence for H3K36me3 (cyan) and an example random spot generated in the nucleus (gray). **b** Conceptual diagram of expected results. Nascent Xist (green) and Tsix (magenta) transcription sites are shown, with larger circles indicating higher transcription levels. Cyan represents H3K36me3, either enriched (dark cyan) or depleted (light cyan) around transcription sites. If Tsix inhibits Xist via H3K36me3, Xist will be negatively correlated with H3K36me3, and Tsix will be positively correlated (left). Without inhibition, both transcripts will be positively correlated with H3K36me3 (right). **c** Quantification of H3K36me3 enrichment (cyan) at varying distances from a transcription site (blue) or a random spot (gray). Using Cochran-Mantel-Haenszel (CMH) statistics to calculate the odds ratio, we assess the enrichment of colocalized H3K36me3 at transcription sites compared to random spots (y-axis) relative to distance from the transcription site (x-axis). **d-i** Quantification of cells at day 2 of differentiation using the approach described in (c). Fold enrichment of H3K36me3 with real transcription sites (blue) relative to random spots (gray). N = 4 biological replica experiments, n = 818 total cells. In each subpanel, m is the number of transcription sites with the designated number of nascent transcripts, t is the number of transcription sites with the number of molecules specified in each panel. **d-f** Fold enrichment of H3K36me3 at Tsix sites with low (d), medium (e), and high (f) transcription. **g-i** Fold enrichment of H3K36me3 at Xist sites with low (g), medium (h), and high (i) transcription. **j-k** Joint distributions of H3K36me3 fold enrichment at transcription site relative to random depending on the nascent transcription of Xist (x-axis) or Tsix (y-axis). t = number of transcription sites in each bin, the p value is from using the CMH test comparing the data in each bin to randomized spots. The specific localization of H3K36me3 is either on the Tsix site of transcription (j) or Xist site of transcription (k).

Because H3K36me3 is deposited by transcription and no known transcription units overlap with the Tsix promoter, we expect higher Tsix transcription to be positively correlated with enrichment of the mark (Fig. 6b, bottom). If H3K36me3 deposition inhibits Xist, transcription sites with low levels of Xist should have more enrichment of the mark nearby, and transcription sites with high levels of Xist should have less enrichment of H3K36me3 (Fig. 6b, top left). If H3K36me3 does not inhibit Xist, then we would expect the opposite relationship, because more transcription of Xist would deposit more of the mark in the gene body (Fig. 6b, top right).

To test these predictions, we quantified fold-enrichment of H3K36me3 localization to transcription sites compared to random spots in the same nuclei (Fig. 6c). In this way, we can visualize the enrichment of histone mark localization at specific distances away from the site of transcription. Similarly to our quantifications with TI, we investigated effects for low, medium, and high levels of nascent transcription at day 2 when Xist is expressed (Fig. 6d-i, Supplementary Fig. 37-39).

We first tested the prediction that Tsix transcription positively correlates with enrichment of H3K36me3. All Tsix transcription sites showed enrichment of H3K36me3 (Fig. 6d-f, Supplementary Fig. 37). Interestingly, Tsix sites with medium-level transcription showed the highest levels of enrichment of the chromatin mark (Fig. 6e). This positive correlation between Tsix transcription and H3K36me3 is consistent with the hypothesis that Tsix transcription deposits H3K36me3.

We then tested the prediction that Xist levels negatively correlate with H3K36me3 at the site of transcription. Low and medium levels of nascent Xist expression were associated with enrichment of H3K36me3 (Fig. 6g, h), while sites with high nascent Xist exhibited levels of H3K36me3 significantly below random (Fig. 6i). Notably, no H3K36me3 spots were detected within 0.25 µm of highly transcribing Xist sites (Fig. 6i, first bin). For day 0 of differentiation when Xist is largely repressed, there was high enrichment of H3K36me3 (Supplementary Fig. 40). This negative correlation between Xist transcription and H3K36me3 is consistent with the hypothesis that H3K36me3 inhibits Xist, a pattern that was previously observed in allele-specific CUT & TAG population experiments and is not observed in housekeeping genes (Supplementary Figs. 41-48) ^65^. We then compared these results to experiments where Setd2 was inhibited during two days of differentiation. We quantified the number of H3K36me3 spots in the cells, the intensity of Tsix and Xist intronic signals, and the fraction of cells with nascent Xist and Tsix expression (Supplementary Fig. 49). Our findings showed that Setd2 inhibition reduces the number of H3K36me3 spots in cells (Supplementary Fig. 49a, b), which in turn decreases the intensity of nascent Tsix expression (Supplementary Fig. 49c, d). However, there was no reduction in nascent Xist expression (Supplementary Fig. 49e, f). Interestingly, we observed an increase in the fraction of cells with nascent Xist expression upon Setd2 inhibition (Supplementary Fig. 49g), while no change was detected in the fraction of cells with nascent Tsix expression under the same conditions (Supplementary Fig. 49h).

Because Tsix and Xist can co-transcribe in the same locus (Fig. 5), we then asked how much H3K36me3 was at transcription sites with different levels of both Tsix and Xist nascent transcription. To be considered the same transcription site, Xist and Tsix signal had to be within 0.75 µm (Supplementary Fig. 50), which is less than the 1 µm minimum distance found with X chromosome pairing ^66^. Surprisingly, we observed the strongest and most significant enrichment of H3K36me3 when Xist and Tsix co-transcribed at medium levels (Fig. 6j, k, center bin, Supplementary Figs. 37). Our findings suggest that medium levels of transcription, which allows for the maximum amount of overlapping co-transcription (Fig. 5k, l), coincide with the most deposition of histone marks when this co-transcription occurs.

Overall, our results show different mechanisms of regulation by lncRNAs at the Xist/Tsix locus depending on the level of nascent transcription. At high levels of transcription, TI causes inhibition in overlapping transcriptional regions. At medium-level transcription, antisense regions can co-transcribe and deposit maximum histone marks for chromatin regulation.

## DISCUSSION

In this study, we established a methodology to test mechanisms of gene regulation by lncRNAs and applied it to the Xist/Tsix system. With our quantitative experimental and analytical framework, we tested multiple mechanisms by investigating single cells and utilizing the natural variability of stochastic gene expression in the system to probe the effects of different expression levels. These different levels of nascent transcription had a profound impact on what mechanisms of regulation were being utilized.

Multiple proposed mechanisms of Tsix regulation of Xist involved binding of the antisense pair (Fig. 1a-b), but it was unknown to what extent they bound in cells. This gap in knowledge resulted from previous studies using either population-based measurements or long RNA-FISH probes that could not detect individual Xist and Tsix molecules ^22,36^. Our smRNA-FISH data analyzed with PICCS revealed that Tsix almost never bound to Xist in mESCs, providing strong evidence against any mechanisms involving their binding (Fig. 2). Our analysis, which shows a high degree of colocalization, co-expression, and correlation between Xist and RepA, suggests that RepA is not an independent transcript but instead a segment of the Xist transcript (Fig. 2, Supplementary Fig. 8, 11, 13). It is improbable that rapid RNA degradation via RNAi would prevent detection in our assays. Previous research has demonstrated that RNA-FISH is capable of detecting even short-lived RNA species with lifespans shorter than three minutes ^52,53^.

Our single-cell quantification of Xist and Tsix also determined that whole-cell levels of Xist and Tsix were not correlated (Fig. 3). The lack of correlation provided additional evidence against *trans* regulation by these lncRNAs at the locus. It was essential to employ rigorous statistical methods because simple visualization or incorrect statistical tests can suggest correlations that are not present (Supplementary Fig. 6). In contrast to previous studies, our methodology did not necessitate the production of mutants that can introduce artifacts leading to misinterpretation of results. Similar analyses can be done on any two genes if there is single-cell variability in the expression of the genes of interest.

Identifying transcription sites as bright RNA-FISH signals in the nucleus also enabled us to test, even with exonic probes, whether the lncRNAs regulated each other in *cis*. Regulation *in cis* had been tested with mutants of specific alleles altering Tsix transcription ^15,17,22,39^, but it had not been tested by quantification of nascent transcription in wildtype cells. We saw a strong negative correlation between Xist and Tsix at the site of transcription, which provided additional evidence that regulation was occurring in *cis* (Fig. 4). Analysis of signal from intronic RNA-FISH probes further supported that regulation occurs in *cis* (Fig. 5) and prompted us to investigate mechanisms involving the act of transcription.

One possible mechanism of transcription-mediated regulation by lncRNAs is Transcriptional Interference (TI) (Fig. 1d). A previous study suggested that TI is an active mechanism at the Xist/Tsix locus ^36^. However, they utilized a doxycycline-inducible system to control Xist expression instead of the native promoter. We instead took advantage of the natural variability in RNA expression and found that TI only occurs at high levels of nascent transcription (Fig. 5). This result is consistent with the findings of Mutzel et al., as inducible promoters tend to be strong and would mimic the cells with high nascent transcription.

Interestingly, we found that medium-level transcription was associated with the highest co-transcription of Xist and Tsix (Fig. 5k, l). Because evidence suggests eukaryotic polymerases cannot pass each other ^57^, antisense co-transcription in overlapping regions has been explained by polymerases completing transcription during short times when the opposite strand is unoccupied. Completed transcripts are then retained for some time nearby ^60^. Our results are consistent with this explanation since the opposite strand would be unoccupied for longer lengths of time during medium transcription levels compared to lengths during high levels. The low colocalization at weakly-transcribing transcription sites (Fig. 5i, j) might be explained by lower local concentration of transcription machinery. Our discovery that antisense co-transcription peaks at moderate nascent transcription levels is a starting point to determine if this observation applies to other antisense transcripts.

Another possible mechanism of transcription-mediated regulation by lncRNAs is through chromatin modification (Fig. 1e) ^2,3^. By quantifying H3K36me3 in the vicinity of transcription sites using quantitative immunofluorescence and smRNA-FISH, we found enrichment of H3K36me3 at Tsix transcription sites and depletion of H3K36me3 at highly transcribed Xist sites (Fig. 6). These results are consistent with the hypothesis that Tsix deposits H3K36me3 to inhibit Xist in single cells ^39,40^.

We found that maximum enrichment of H3K36me3 occurs when Tsix and Xist are co-transcribed at medium levels. This unexpected outcome may be a function of varying local concentrations of polymerases and histones as illustrated in Fig. 7. While higher nascent transcription levels would have more polymerases to recruit chromatin modifiers, strong transcription can also remove histones and create nucleosome-depleted regions (Fig. 7, top and bottom) ^67^. In contrast, we speculate that medium transcription levels enable polymerases to transcribe around histone proteins without removing them ^68^. As a result, the highest local density of modifiable histone proteins is observed at medium transcription levels (Fig. 7, middle).

**Fig. 7.**
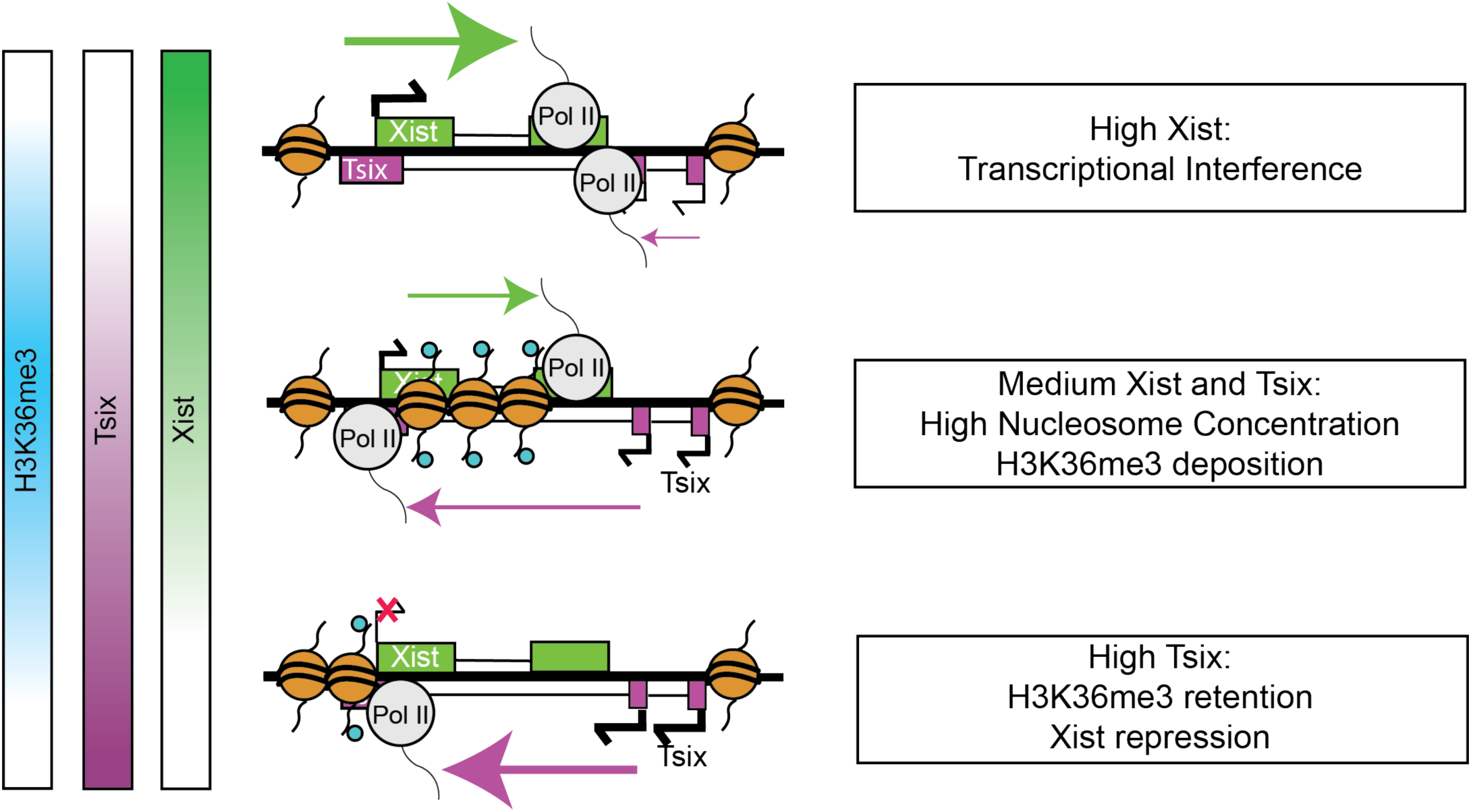
Natural variability of transcription enables different mechanisms of Xist and Tsix regulation within a population of cells. Bars to the left indicate the H3K36me3 levels (cyan) that are maximized near the site at medium transcription of both Tsix (magenta) and Xist (green). The conceptual diagram showing DNA (black lines) wrapped around histone proteins (orange spheres) with and without the H3K36me3 (cyan circles) modification at the Xist and Tsix locus depending on the amount of Tsix and Xist transcription. (Top) High nascent transcription of Xist results in transcriptional interference and depletion of H3K36me3 near the site of Xist transcription. (Middle) Medium nascent transcription of both Xist and Tsix is correlated with the highest enrichment of H3K36me3 near the site of Xist and Tsix co-transcription. (Bottom) High nascent Tsix transcription results in transcriptional interference and modest enrichment of H3K36me3 levels.

When both Xist and Tsix are transcribing, there is also a higher density of polymerases to recruit chromatin modifiers (Fig. 7, middle). Although transcription sites with medium Xist and Tsix levels represent a small subset of all detected transcription sites, they might signify a crucial temporary state for establishing H3K36me3 that persists, allowing for further regulation. Through Setd2 inhibitor studies (Supplementary Fig. 49), we observed that the intensity of Tsix transcription sites decreased, suggesting that the number of polymerases transcribing Tsix is reduced. However, the number of cells initiating Tsix transcription did not change. In contrast, while the intensity of Xist transcription sites remained unchanged during Setd2 inhibition, the fraction of cells exhibiting nascent Xist expression increased. This indicates that Setd2 regulates the frequency of polymerase initiation responsible for transcribing Xist. In summary, our methodology provided single-cell evidence for Xist inhibition involving Tsix-induced H3K36me3 and identified an interaction with medium-level co-transcription that would have been undetectable in population-level experiments or those with artificial promotors.

Our quantitative, high-resolution methodology contributes novel and critical insights into the regulation at the Xist and Tsix locus despite it being a focus of significant research for over two decades. We were able to test a variety of mechanisms in a single study. Since this methodology does not necessitate the generation of mutants, it is a promising way to rapidly investigate the many other antisense transcripts whose function and mechanism remain unknown in their native context. Our methodology can also be used to test other mechanisms involving RNA-RNA interactions. Overall, our results support two different regulatory mechanisms within the same population of cells: chromatin modification at medium transcription and TI at high transcription (Fig. 7).

Some limitations of this study are that we only quantified snapshots of different cells instead of following the same cells over time, so we do not know when or how long different states of transcription occur. Future studies utilizing timelapse microscopy could lead to complementary insights ^60^. Although we did not find any evidence of Xist and Tsix binding together, we cannot exclude that Xist and Tsix binding results in a short-lived complex that is only detectible in small amounts in our assay. However, previous studies suggest that the timescale of double-stranded RNA degradation by Dicer is in minutes ^69^, which should make the complex more visible in our data. Xist and Tsix might also bind to each other at the site of transcription only but then dissociate later. Although we determined locations and distances in three dimensions, we did not determine the precise genomic locations of the nascent transcripts and histone marks.

Our findings have general implications for future studies or therapeutics targeting lncRNAs because the lncRNA of interest might have multiple mechanisms, which could confound results or lead to suboptimal outcomes. For instance, strategies such as antisense oligonucleotides (ASOs) that lower RNA expression are promising avenues for disease treatments ^70^. However, lowering the expression of a lncRNA might only eliminate mechanisms that are employed at high transcription levels (such as TI at the Xist/Tsix locus), while retaining other mechanisms requiring lower transcription (such as chromatin modification at medium levels of Tsix). Similarly, inhibiting a protein such as Setd2, which causes H3K36me3 and is a target of recent clinical studies ^11,12^, might only affect some of the mechanisms of regulation and may not interfere with TI at high expression. Our study suggests that a combinatorial approach targeting multiple mechanisms might be necessary for effective future mechanistic lncRNA studies and the development and application of therapeutics.

## METHODS

### Experimental methods

#### Cell culture

F1 2-1 mESCs ^17^ were obtained from the lab of Dr. Rudolf Jaenisch and validated by karyotyping. The F1 2-1 mESCs were thawed with 1 million seeded cells on 75 cm^2^ tissue culture flasks with vented caps (Falcon 353110) gelatinized with EmbryoMax 0.1% Gelatin Solution (Millipore ES-006-B) for 30 minutes at 37°C. Cells were grown with 2i media composed of: DMEM with high glucose (Life Technologies 11960-044), 0.5x N2 Supplement (Gibco 17502-048), 0.5x B27 supplement (Gibco 17504044), 25 mM HEPES (Gibco 15630-030), 0.5x MEM NEAA (Life technologies 11140-050), 1x (100 U/mL) Penicillin-Streptomycin (Gibco 15140122), 100 μM 2-mercaptoethanol (Life Technologies 21985-023), 1000 U/mL LIF (EMD Millipore ESG1106), 0.25x GlutaMAX^TM^ (Gibco, Catalog#: 35050-061), and 1x (1 mM) sodium pyruvate (Gibco 11360-070), 20 μg/mL human insulin (Sigma I9278-5ML), 1 μM (Sigma PD0325901), 3 μM (Sigma CHIR99021), 1000 U/mL LIF (EMD Millipore ESG1107) at 37°C in a 5% CO_2_ humidity-controlled environment for two passages before experiments. Media was changed every day, and cells were grown 3 days before passaging.

To begin undirected differentiation in adherent conditions, cells were washed twice with 1x PBS and dissociated from the flasks with accutase (Merck SF006). 1 million cells were then plated on 15 cm tissue culture dishes gelatinized with 0.1% Embryomax gelatin for 30 minutes at 37°C. They were grown in differentiation media consisting of: DMEM with high glucose (Life Technologies 11960-044), 15% Heat Inactivated FBS (Gibco 16140-071), 25 mM HEPES (Gibco 15630-030), 1x MEM NEAA (Sigma M7145), 1x (100 U/mL) Penicillin-Streptomycin (Gibco 15140122), 100 μM 2-mercaptoethanol (Life Technologies 21985-023), 1x GlutaMAX^TM^ (Gibco, Catalog#: 35050-061), and 1x (1 mM) sodium pyruvate (Gibco 11360-070) at 37°C in a 5% CO_2_ humidity-controlled environment. Differentiation plates were split if >70% confluent.

Setd2 inhibitor studies were conducted using 500 nM of SETD2i (inhibitor EPZ-719, MedChemExpress) in 2i media during two days of differentiation, as previously described. RNA-FISH probes and H3K36me3 antibodies were used as outlined above.

#### Cell Fixation

mESCs were dissociated after washing with 1x PBS using 0.25x accutase when cultured in 2i media and 0.05% trypsin (Gibco 25300054) when in FBS-containing media. The cell suspension was centrifuged for 5 minutes at 300 g, washed with 1x PBS, and then fixed for 8-10 minutes at room temperature with a 3.7% formaldehyde solution in 1x PBS. The cells were washed twice with 1x PBS and then permeabilized with 70% ethanol at 4°C for at least one hour, usually overnight.

#### DAPI, RNA-FISH, and IF staining

After cell permeabilization, use BSA coated tips to transfer cells to a low bind tube. After centrifugation and EtOH aspiration, resuspend cells in 100ul wash buffer (PBS with 10% formamide), centrifuge at 300g, aspirate wash buffer completely, and then resuspend cells in 50 ul hybridization buffer (10% dextran (Sigma Aldrich, D8906), 10 mM vandyl-ribonucleoside complex (NEB, S1402S), 0.02% RNAse-free BSA (Ambion, AM2616), 1mg /ml E.coli tRNA (Sigma, R4251), 2x SSC (Ambion, AM9763), 10% formamid (Ambion, AM9342)) that contains RNA-FISH probes (1:1000 dilution) and antibody if necessary and incubate at 37C overnight. After hybridization resuspend cells in 1ml wash buffer, centrifuge at 300g, aspirate solution, resuspend cells in 1ml wash buffer with DAPI (Sigma, D9564, 150 ng/ml) and secondary antibody if required. Incubate for 30min at 37C. Repeat wash step one more time without DAPI and antibodies. ^52^ Primary antibodies were added at the same time as the RNA-FISH probes, and secondary antibody was added during the DAPI staining (see table for antibodies and concentrations). The primary antibody concentrations were determined with an optimization experiment for when individual spots were visible and minimal background fluorescence was detected (Fig. 5, Supplementary Fig. 7).

#### Microscopy

Cells were imaged with epifluorescence at 100x with a 500ms exposure time, at 300nm per z-step, from the top to the bottom of the cells ^50^. Micro-Manager software version 1.4 was used to control a Nikon Ti Eclipse epifluorescent microscope equipped with perfect focus (Nikon), a 100× VC DIC lens (Nikon), an X-cite fluorescent light source (Excelitas), and an Orca Flash 4v2 CMOS camera (Hamamatsu). Mammalian cells were on the ibidi 15 μ-Slide Angiogenesis (81506) in wells coated with 0.01% poly-D-lysine (Cultrex 3439-100-01) for 10 minutes. Fluorescent signal was imaged with Semrock Brightline filters for DAPI (DAPI-5060C-NTE-ZERO), CY5 (NIK-0013 RevA-NTE-ZERO), TAMRA (SpGold-B-NTE-ZERO), and AF488 (FITC-2024B-NTE-ZERO).

### Computational methods

#### Cell Segmentation

Nuclei were segmented in 3D and cells were segmented in 2D by utilizing CellDissect, an automatic segmentation pipeline ^50^. The DAPI signal of the nucleus was automatically thresholded to maximize the number of non-overlapping signals presenting individual nuclei. The nuclei are used as seeds to segment the cell boundary in 2D automatically.

#### Probe Design

Sequences for Xist and Tsix were downloaded from the UCSC genome browser, and genomic sequences specific to the two background strains present in the cell line (129S1/SvImJ and Castaneous) were downloaded from the Mouse Genomes Project from the Sanger Institute (http://www.sanger.ac.uk/resources/mouse/genomes/). For the Xist, Tsix, and RepA exonic probes, Stellaris software (Biosearch technologies) was utilized with certain considerations for sequence input and downstream probe pruning. For Xist, the input sequence was designed to avoid the repeat sequences known to be important for Xist function, since secondary structure could inhibit probe binding. For Tsix, the region overlapping Xist was avoided. Since the RepA region overlaps with Tsix and is part of the Repeat A region of Xist critical for X chromosome inactivation, neither of these could be avoided. These sequences were input into the Stellaris software with the maximum masking level of 5 to generate 20mer probes with 3 base spacing. Probes with tandem repeats were removed by utilizing a tandem repeat finder (Tandem Repeats Finder Advanced Submit Page (bu.edu) on the original sequence (except for with the RepA probes), and probes with SNPs that were different between 129 and Cast backgrounds were avoided by comparing their genomic sequences. Custom MATLAB software that utilized online BLASTing and the "Mouse Genomic+Transcript" option determined the number of off-targets. Any probes with 19 or more matching bases of an off-target were excluded, and probes with lower numbers of off-targets were prioritized to be in the final probe set.

The intronic probes were designed using custom MATLAB software that utilized BLAST+ and a local BLAST database generated from the mm10 genome, mouse rRNAs from RNACentral.org, and mouse transcripts from RefSeq. Genomic and transcript sequences for Xist and Tsix were again downloaded from the UCSC genome browser, and the strain-specific sequences were again used from the Sanger institute. For Xist and Tsix 3’ intronic probes, the intronic regions of Xist were used for the probe design, again avoiding the SNPs that differ between 129 and Castaneous mouse strains. Probes were BLASTed against the mouse database, and probes were chosen based on minimal off-target binding. Probes with matches against Hsp90 and any rRNAs were excluded. All probes that had matches against highly expressed genes in differentiating mESCs with the same background (the top 465 expressed genes from the RNA-Seq dataset in Marks Genome Biology 2015) ^49^ were also excluded. The resulting probes for Xist and Tsix 3’ were interlaced, with half designed against the Tsix strand and half against the Xist strand. The same procedure was performed for Tsix 5’ intronic probes, and the region with known DXPas34 transcription was avoided.

#### Probe Creation

Probes were ordered from Biosearch Technologies with a 3’ Amine modification. All probes in a set were then pooled, conjugated to the appropriate dye, and then purified using HPLC. Xist Exon probes were coupled with CY5, RepA probes were coupled to TAMRA, Tsix Exon Probes were coupled to TAMRA, Xist Intron probes were coupled with TAMRA (for example in Supplementary Fig. 7) and CY5 (for experimental data), and Tsix 5’ Intron and Tsix 3’ Intron were coupled to TAMRA.

#### Image Analysis

Data analysis was performed using custom MATLAB software. Spot detection for RNA-FISH and IF experiments was done using manual thresholding of each channel in images put through Gaussian (to remove noise) and Laplacian of a Gaussian (edge detection) filters. A single slice was used for visualization during thresholding for the IF images due to high densities of spots, while a maximum intensity projection of slices within the bottom and top 10 slices was used for the RNA-FISH images with a low density of spots. For all channels with RNA-FISH signal, a custom algorithm was used to determine the presence of clouds or transcription sites. This algorithm searched for the brightest determined spot, and then determined a 3D connected volume of signal above background attached to this spot. If another spot was present in the cell, the algorithm was used again for a total of two connected volumes determined. These were used for quantification of possible transcription sites and clouds in downstream analysis. For each determined spot and cloud, signal above background was calculated by subtracting background determined using a broad Gaussian filter from the raw images.

#### Data Analysis

The intensity of single RNA spots for each channel was determined by plotting the probability distribution of total signal above background for each determined spot and cloud. The intensity with the highest probability density was used in downstream analyses as the intensity of an individual RNA molecule (Supplementary Fig. 5).

To determine the colocalization of individual RNA molecules visualized with Exonic RNA-FISH probes (Fig. 2), transcription sites were filtered out by ignoring spots with two times the estimated intensity of a single spot. The two brightest spots in each cell were also excluded since they could be transcription sites even if they were not bright. The Euclidian distance was determined between every spot in one channel (e.g. Xist) with every spot in the other channel (e.g. Tsix) in 3D and plotted the total average colocalized spots against distance. The distance axis is on cubic scale for a linear increase in volume from 0-1.5 µm. To determine background colocalization, we fit the data from 0.75-1.5 µm to a line. The y intercept of this line corresponds to the fraction of colocalization as described previously ^51^. Analysis using intronic RNA-FISH probes to visualize nascent transcription (Fig. 5 and 6) utilized the same method, except the two brightest spots in each cell were considered transcription sites.

To calculate the total RNA in each cell, the number of transcripts in every cloud was added to the number of individual puncta within the same cell for Xist, while the total number of puncta was used for Tsix (Fig. 3). We estimated the number of transcripts in every cloud or transcription site by taking the total fluorescence intensity and dividing by the intensity of a single transcript (Fig. 4-6).

To determine the enrichment of histone marks with Xist and Tsix transcription sites, we first calculated the distances between Xist or Tsix transcription sites and spots of histone marks. For every transcription site, 500 random spots were generated inside the same nuclei of the same cells, and the distances between them and the histone marks were also calculated. The number of colocalized histone marks within each bin was then calculated for the data as well as the randomized spots, and the odds ratio of the Cochran Mantel-Haenszel (CMH) test was used to calculate enrichment. For colocalized Xist and Tsix nascent transcription, Xist and Tsix were considered colocalized if they were within 0.75 µm, which is within the 1 µm minimum distance shown previously for paired chromosomes ^66^ (Supplementary Fig. 15). Enrichment and p values were again calculated with CMH statistics.

## Supporting information

Supp Mat

## ACKNOWLEDGMENTS

We like to thank members of the Neuert lab for their feedback on the manuscript. We like to thank Dr. Rudolf Jaenisch for sharing the female F1 2-1 mESCs and we like to thank Drs. Jeannie T. Lee and Rudolf Jaenisch for sharing protocols for these mESC. We also like to thank Dr. Mark Magnuson and the Vanderbilt Stem Cell core for guidance on culturing F1 2-1 mESCs, and Dr. Frank Mason for sharing the SETD2 inhibitor. G.N. is supported by NIH, United States, DP2 GM11484901, NIH R01GM115892, NIH R01GM140240, Vanderbilt Deans Faculty Fellow award, and Vanderbilt Startup Funds. B.K.K. was supported by T32GM008320. This study used resources at the Advanced Computing Center for Research and Education (ACCRE) at Vanderbilt University, Nashville, TN (NIH, United States, S10 Shared Instrumentation Grant 1S10OD023680-01).

## AUTHOR CONTRIBUTIONS

Conceptualization, G.N., B.K.K.; methodology, G.N., B.K.K.; software, G.N., B.K.K.; validation, G.N., B.K.K.; formal analysis, B.K.K.; investigation, B.K.K.; data curation, B.K.K., J.A.; writing – original draft, B.K.K.; writing – review & editing, G.N., B.K.K.; visualization, G.N., B.K.K.; supervision, G.N., project administration, G.N.; funding acquisition, G.N.

