## Supplementary material for "Transcriptional stochasticity reveals multiple mechanisms of long non-coding RNA regulation at the Xist – Tsix locus": Supp Mat

Supplementary Figures 1-50

Supplementary Table 1

### SUPPLEMENTARY METHODS

CUT & TAG data analysis:

For the analyses shown in Supplementary Fig. 41–49, we obtained CUT&TAG for H3K36me3 and RNA-seq data from reference 65. These data were converted into WIG format using the bigWigToWig command line tool, and the resulting WIG files were imported into MATLAB 2024b. We defined genomic regions as described in Supplementary Fig. 41 and calculated the read density per kilobase of gene length and per million sequencing reads (RPKM) for the H3K36me3 CUT&TAG data and RNA-seq data at various time points. The samples from reference 65 were collected under different conditions for the XXΔXic genotype, at days 0, 2, and 4, of differentiation with biological replicates for each condition.

**SUPPLEMENTARY FIGURES**

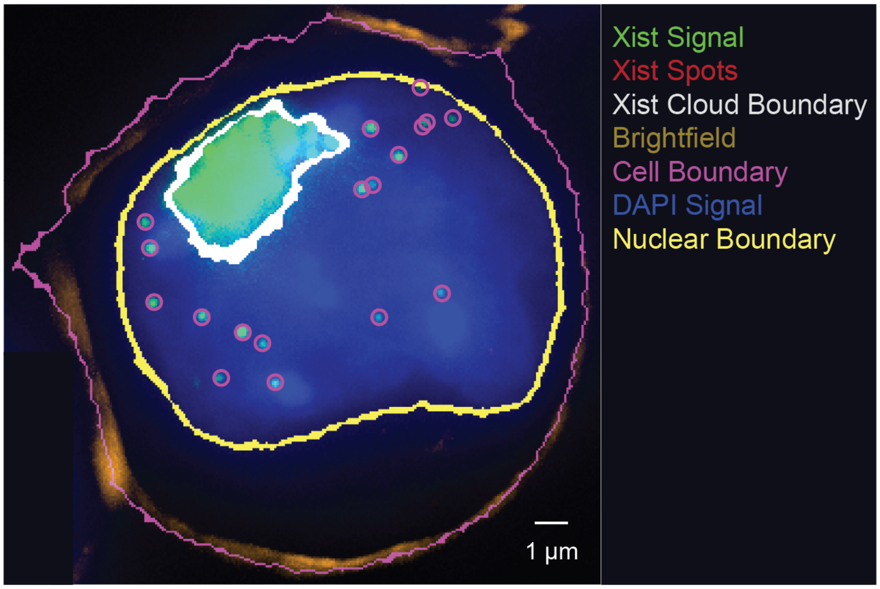


**Supplementary Fig. 1. Example of Boundary detection and Spot Determination**. For cellular and nuclear segmentation, we utilized CellDissect (Kesler et al., 2019). This software uses the brightfield signal of the cell boundary (gold) to determine cell boundaries in 2D (magenta line), and DAPI signal (blue) to segment the nuclear boundaries in 3D (yellow line). RNA-FISH signal (green) is used to determine a boundary of a cloud (while line) and the locations of individual RNA molecules (red circles).

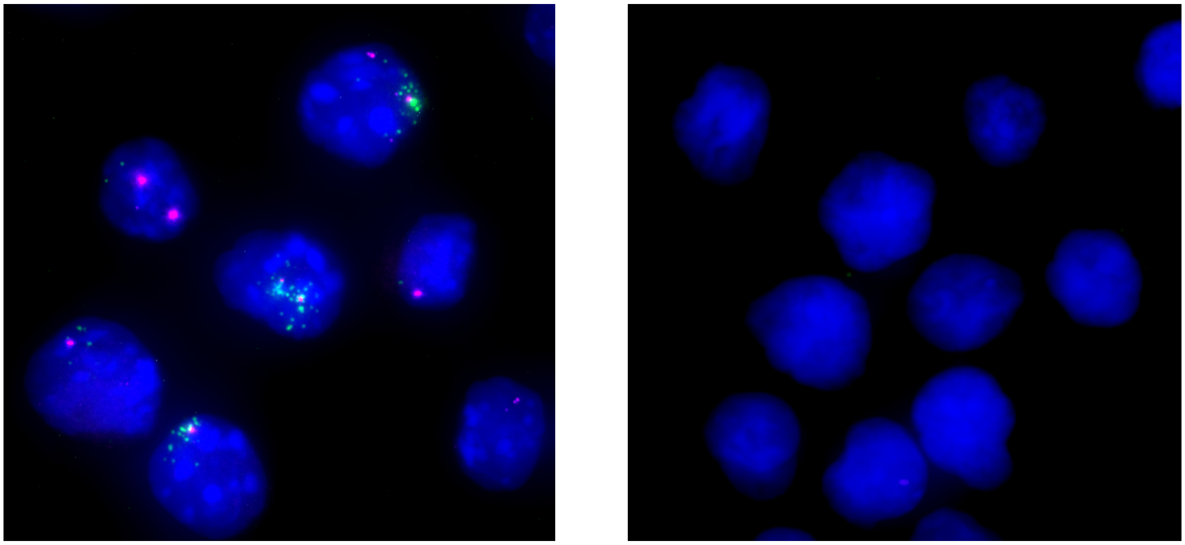

**Supplementary Fig. 2. Specificity of RNA-FISH probes.** (Left) Xist (green) and Tsix (magenta) exon probes in mESC after 2 days of differentiation. (Right) Same probes in human Jurkat cells demonstrating probe specificity. Nucleus stained with DAPI (blue).

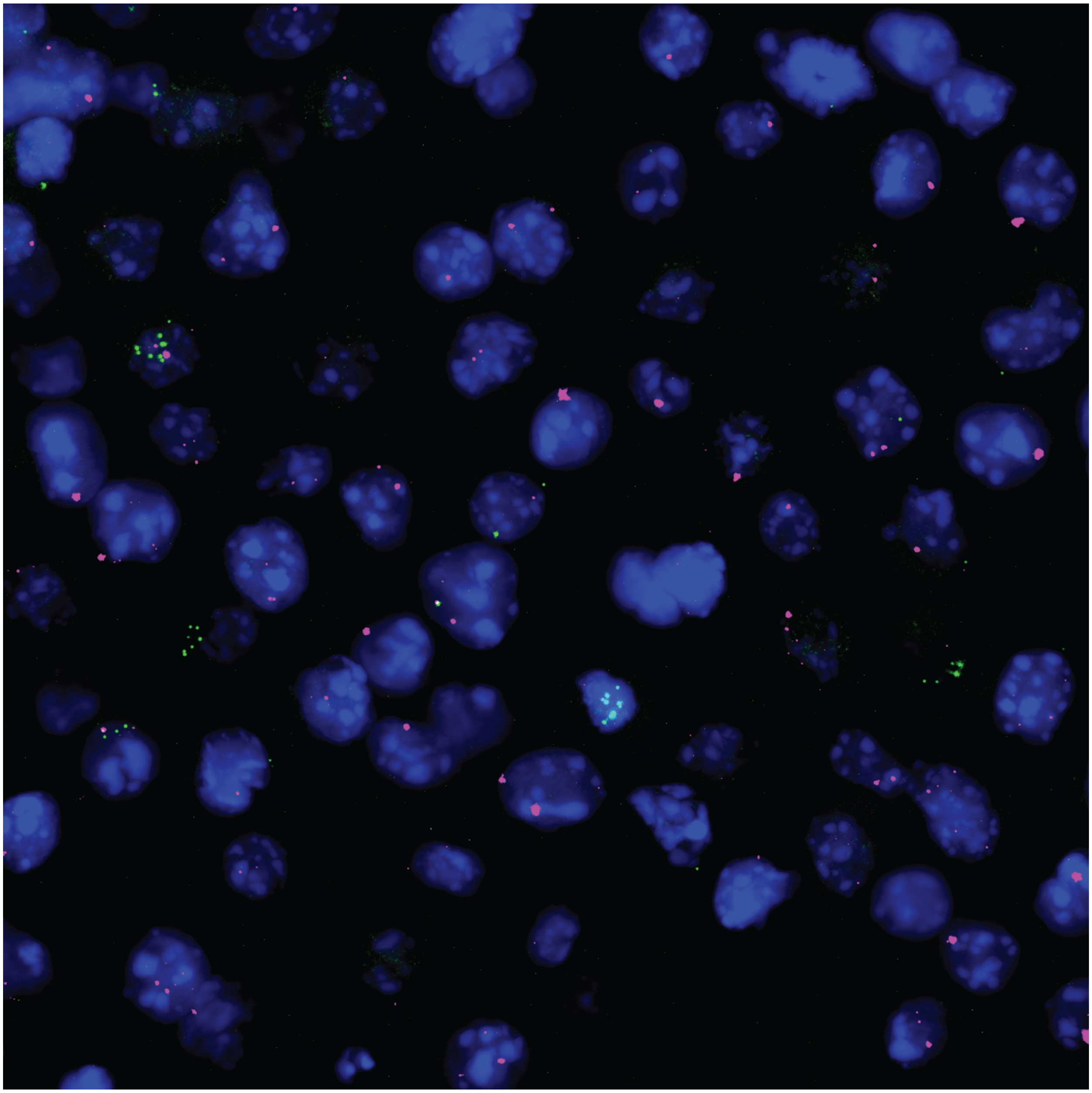

**Supplementary Fig. 3. Example Image of Xist and Tsix Exonic Probes at day 0.** An example image of RNA-FISH signal in mESCs at 0 days. Xist (green) and Tsix exonic probe signals (magenta) visualizes both mature transcripts and the site of transcription. Colocalization of Xist and Tsix signal is shown in white. The nucleus is stained with DAPI (blue).

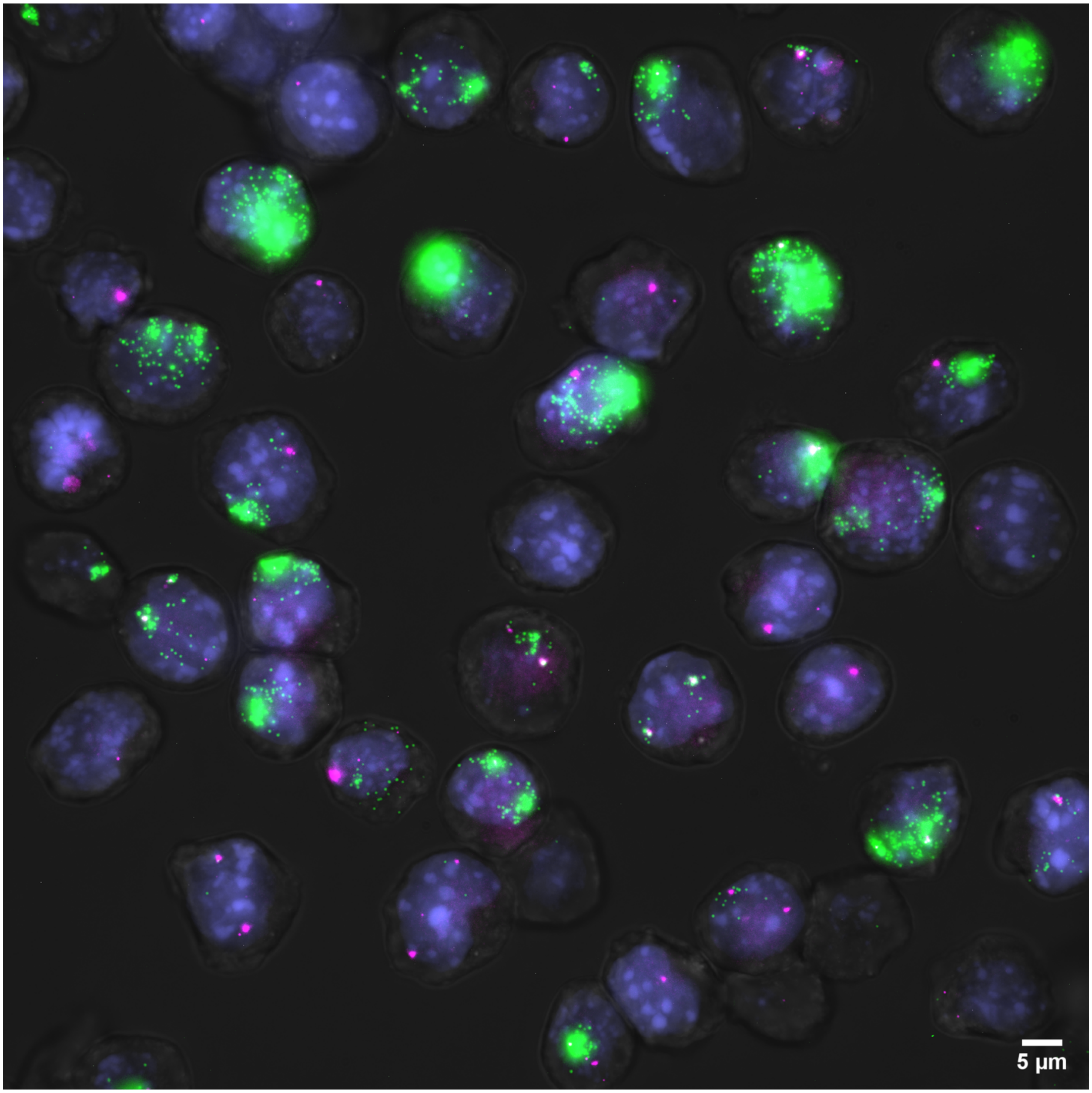


**Supplementary Fig. 4. Example Image of Xist and Tsix Exonic Probes.** An example image of RNA-FISH signal in mESCs differentiated for 2 days. Xist Exonic probe signal (green, log10 scale) and Tsix exonic probe signal (magenta) visualizes both mature transcripts and the site of transcription. Colocalization of Xist and Tsix signal is shown in white. The nucleus is stained with DAPI (blue).

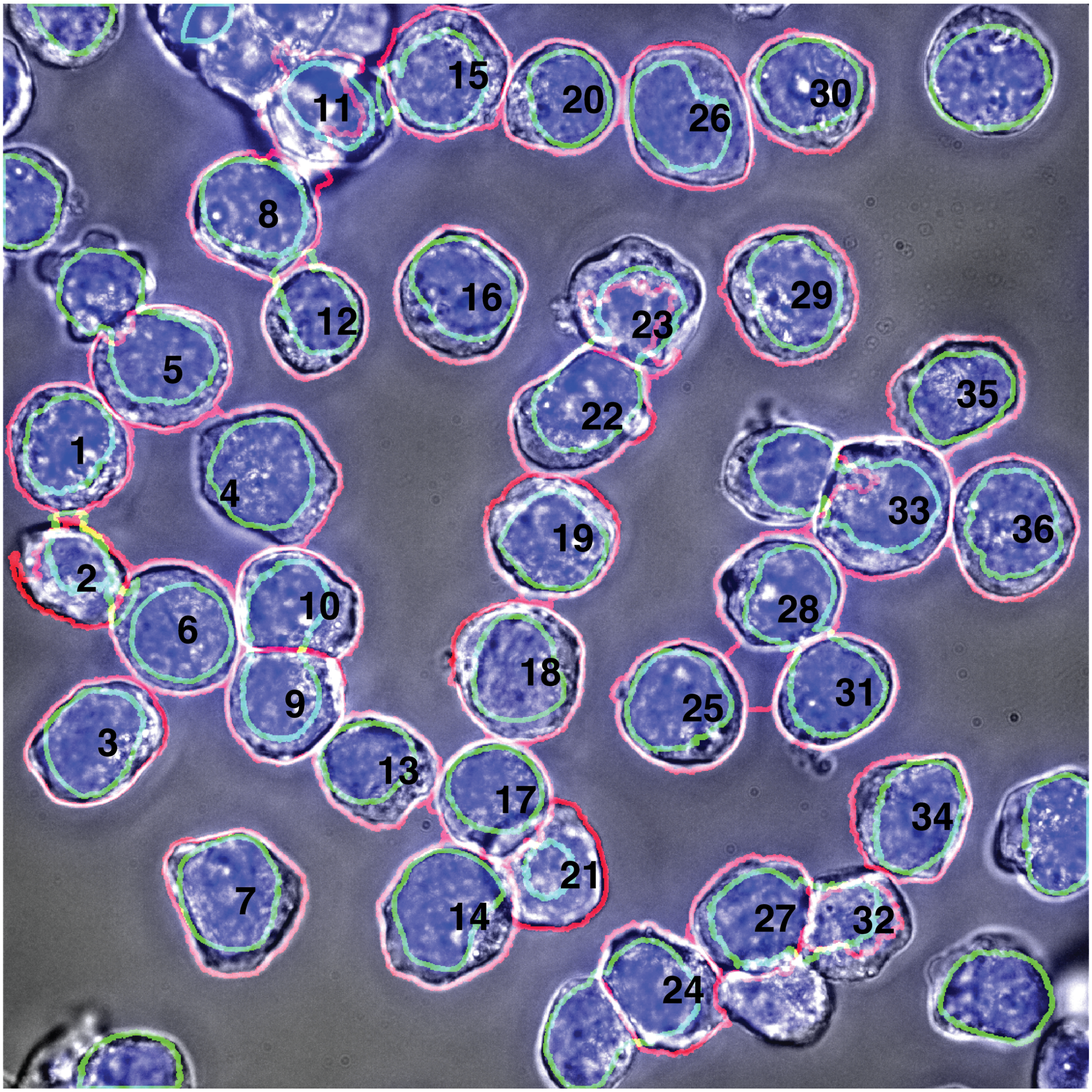


**Supplementary Fig. 5. Example Image of segmented cells from Supplementary Fig. 4.** This image shows segmented cells from Supplementary Fig. 4. The brightfield image is displayed in grey. The nucleus is stained with DAPI and appears in blue. The segmented nucleus is highlighted in green, while the segmented cell boundary is outlined in red. Black numbers are used to identify the segmented cells. Cells located on the border of the image and cells with only nuclear segmentation are not included in the analysis.

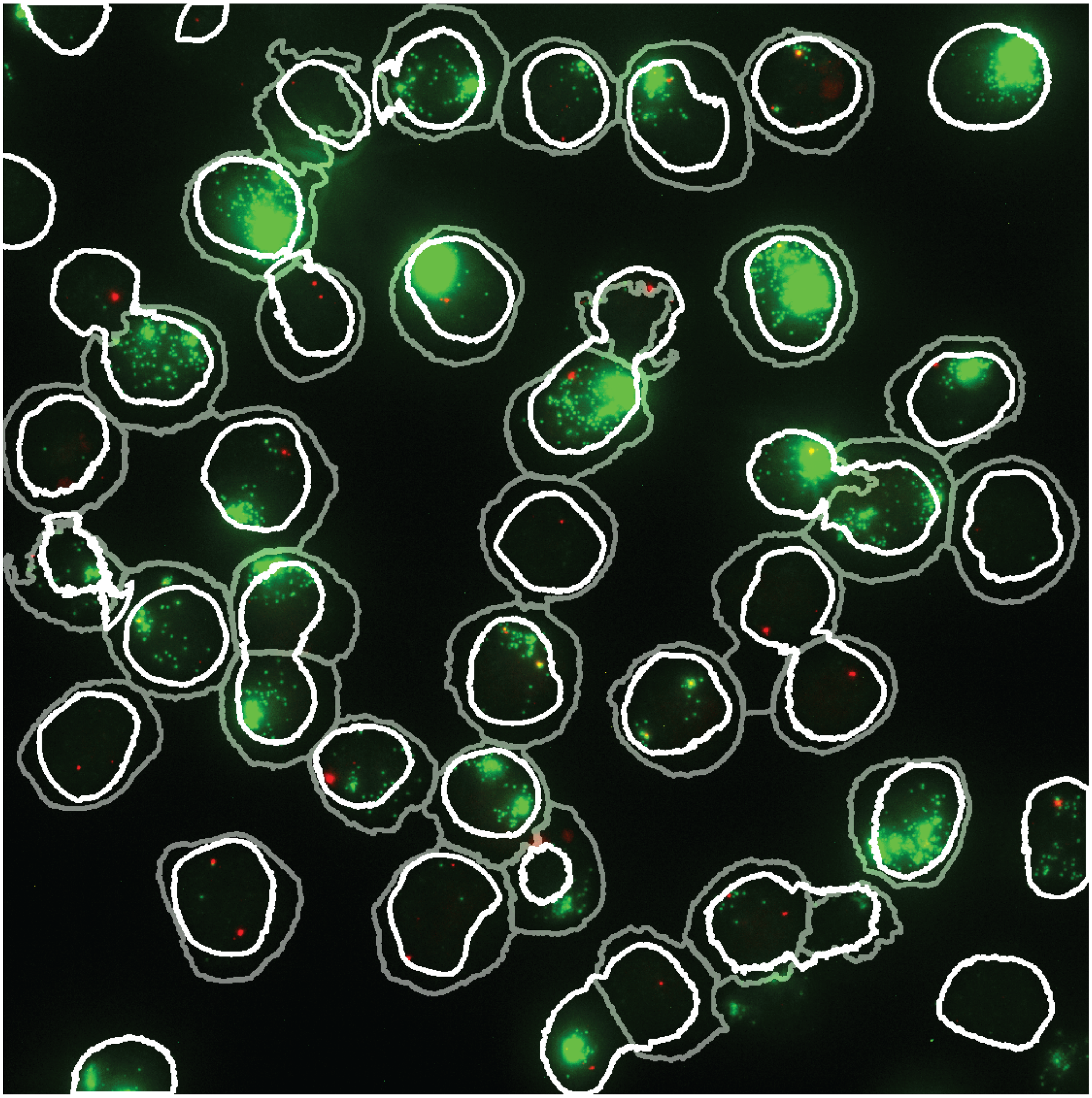

**Supplementary Fig. 6. Example Image segmented cells expressing Xist and Tsix using exonic probes from Supplementary Fig. 4.** This image shows an RNA-FISH signal in mouse embryonic stem cells (mESCs) differentiated for 2 days. The Xist exonic probe signal is shown in green, and the Tsix exonic probe signal is shown in red, visualizing both mature transcripts and the site of transcription. Colocalization of Xist and Tsix signals is displayed in yellow. The nuclear segmentation boundary is marked in white, and the cell segmentation boundary is marked in grey.

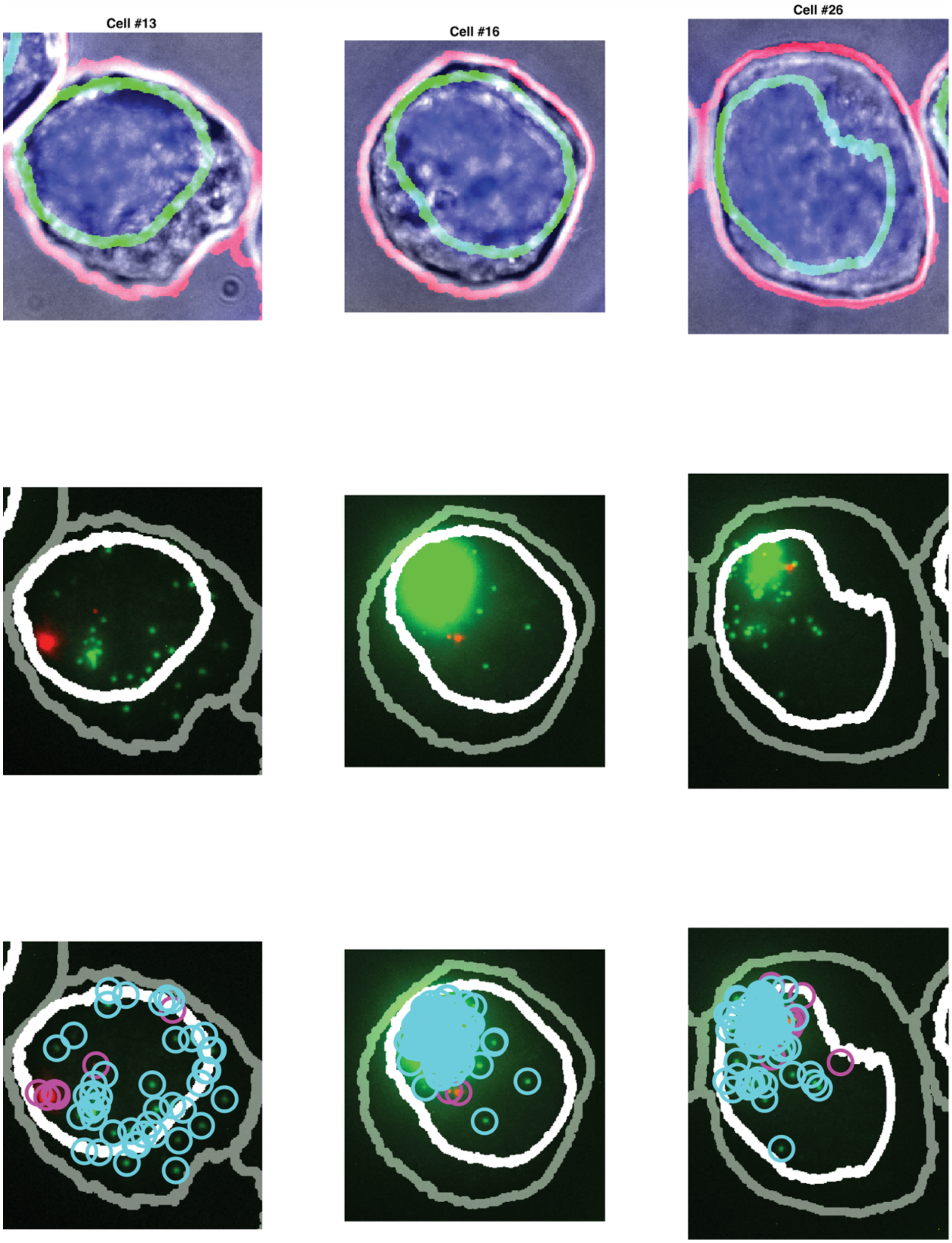


**Supplementary Fig. 7. Example Cells from Supplementary Fig. 5 Demonstrating Cell Segmentation, Nuclear Segmentation, and RNA Spot Counting.** The top panel shows nuclear and cell segmentation. The middle panel displays the RNA-FISH signal for mature Xist transcripts in green and Tsix transcripts in red. The bottom panel illustrates RNA spot detection, with Xist transcripts marked by cyan circles and Tsix transcripts marked by magenta circles.

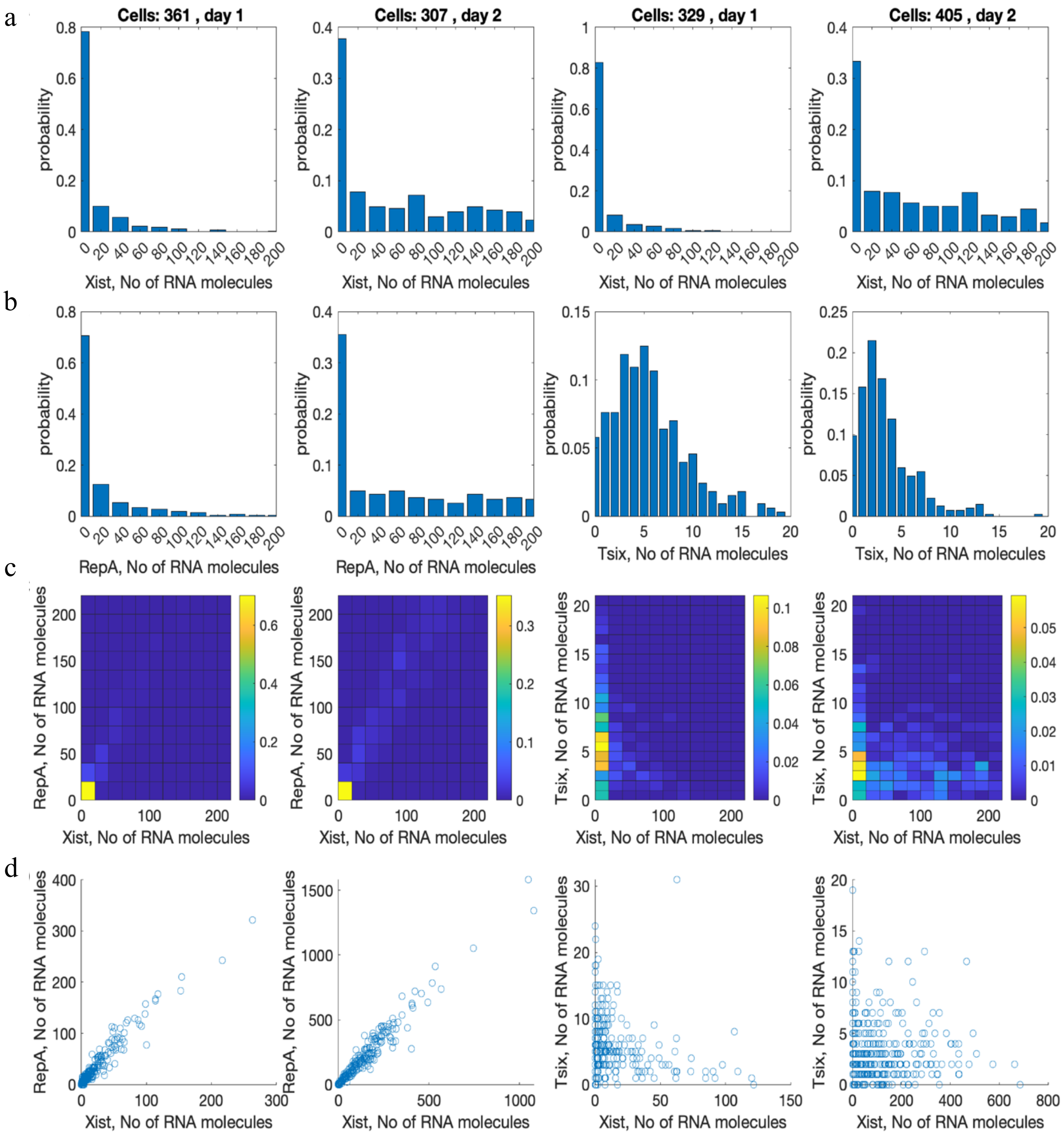


**Supplementary Fig. 8. Detailed Distributions from Fig. 2. a-b** Show the marginal probability distributions for each of the replica experiments for Xist exonic transcript (a), and RepA transcript and Tsix exonic transcript (b). c-d Display the joint probability distributions and scatter plots for each of the replica experiments for Xist and RepA, as well as Xist and Tsix, for days 1 and 2.

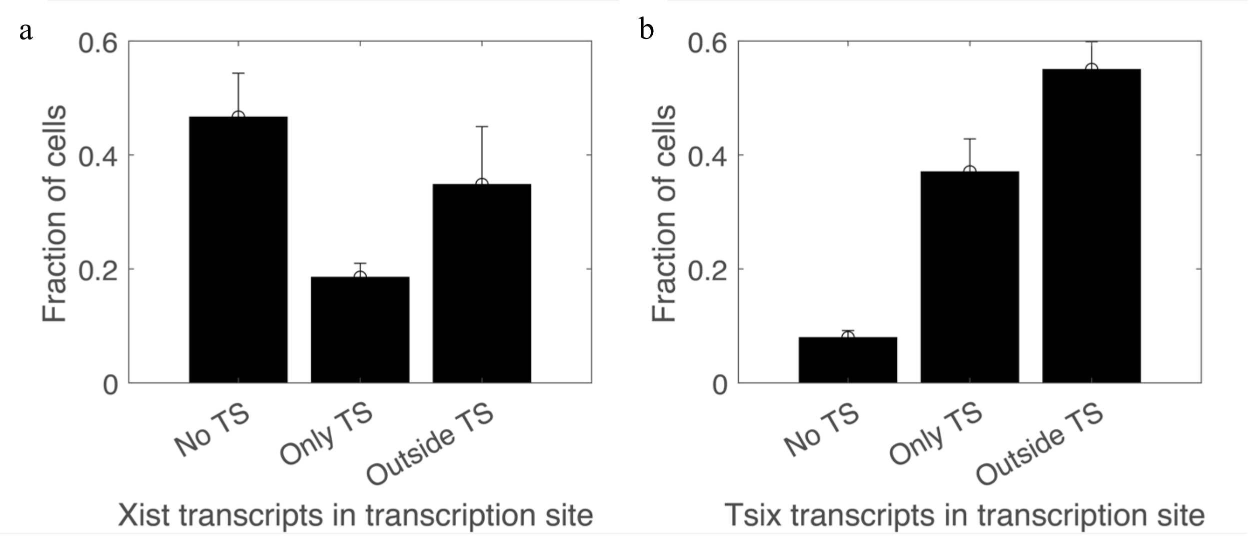


**Supplementary Fig. 9. Spatial Distribution of Xist and Tsix Exonic Transcripts. a** Shows the fraction of cells that have Xist exonic transcripts. **b** shows the fraction of cells that have Tsix exonic transcripts. The categories include cells with transcripts not present in the transcription site (no TS), only present in the transcription site (only TS), and present outside the transcription site (outside TS).

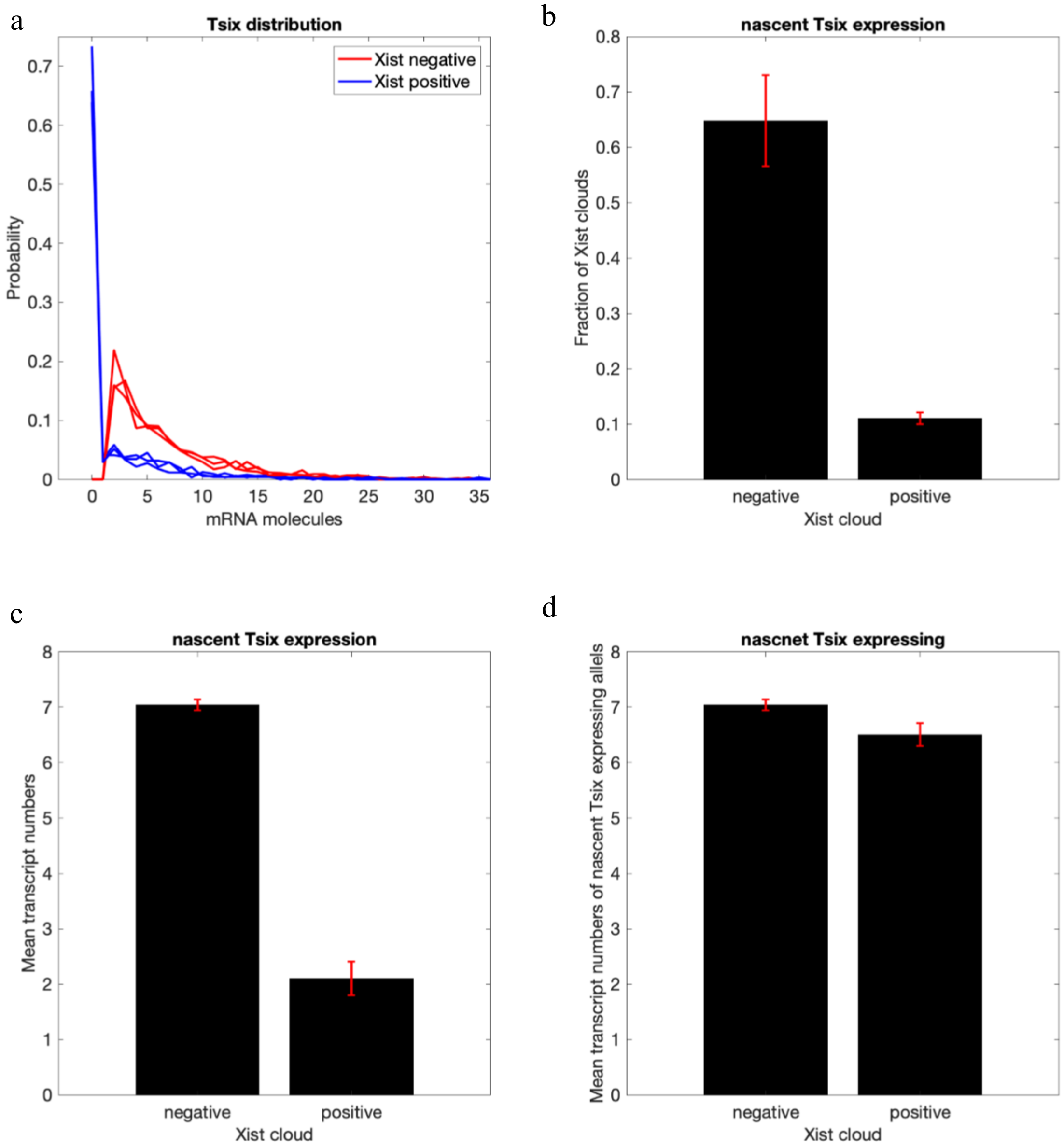


**Supplementary Fig. 10. Tsix Conditional Expression in Xist Cloud Positive and Negative Cells. a** Shows the probability distributions of nascent exonic Tsix expression in cells with (blue) and without (red) an exonic Xist cloud. Each line represents a biological replicate experiment. **b** Displays the fraction of cells with nascent exonic Tsix expression in exonic Xist cloud positive and negative cells. **c** Illustrates the mean nascent exonic Tsix transcript expression in exonic Xist cloud positive and negative cells. **d** Shows the mean number of nascent transcripts in cells that express nascent exonic Tsix, indicating that if cells express nascent exonic Tsix, their mean nascent expression levels are only slightly different between exonic Xist cloud positive and negative cells.

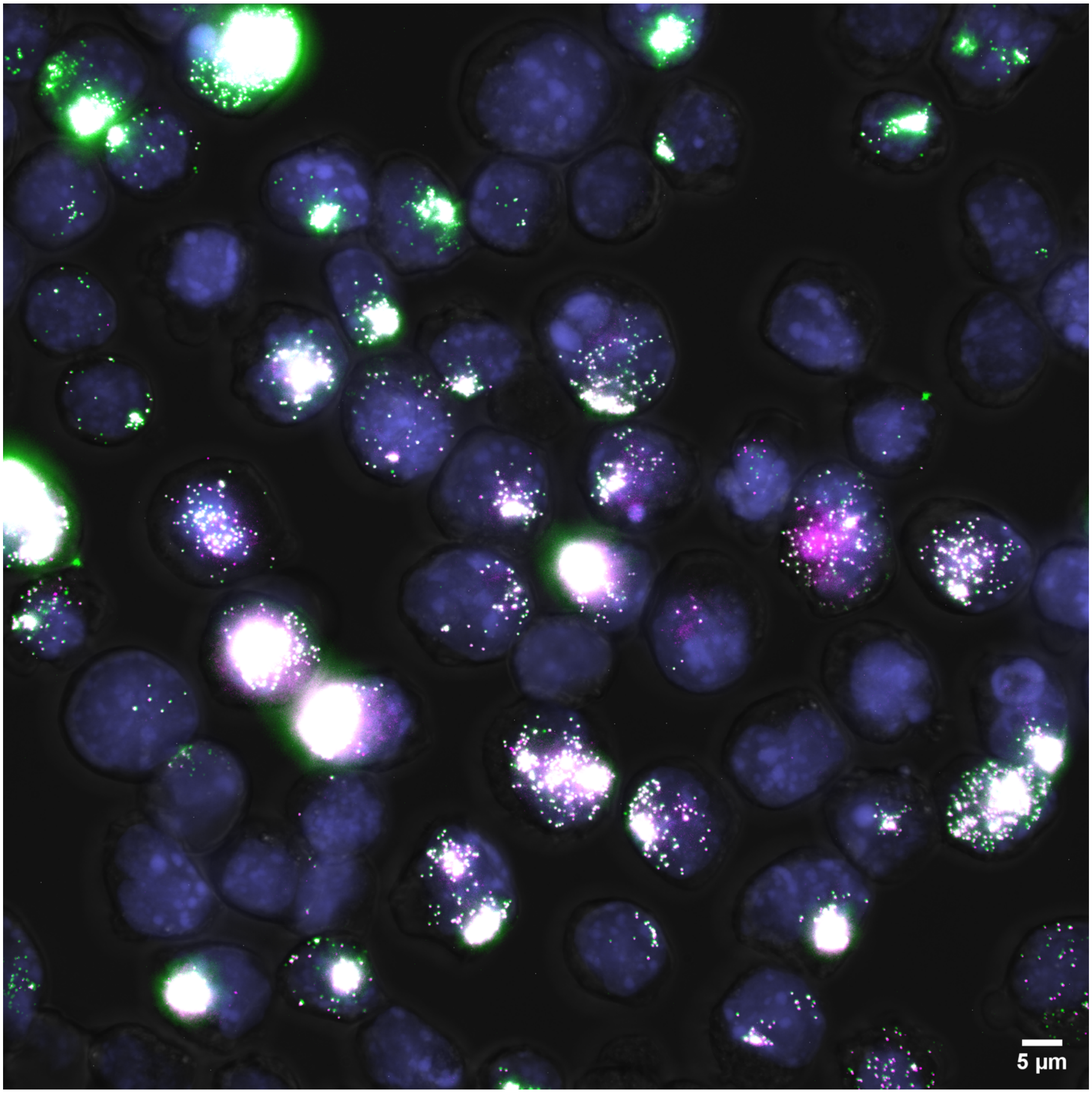


**Supplementary Fig. 11. Example Image of Xist Exonic and RepA Probes.** An example image of RNA-FISH signal in mESCs differentiated for 2 days. Xist Exonic probe signal (green) and RepA probe signal (magenta) visualizes both mature transcripts and the site of transcription. Colocalization of Xist and RepA signal is shown in white. DAPI stained nucleus in blue.

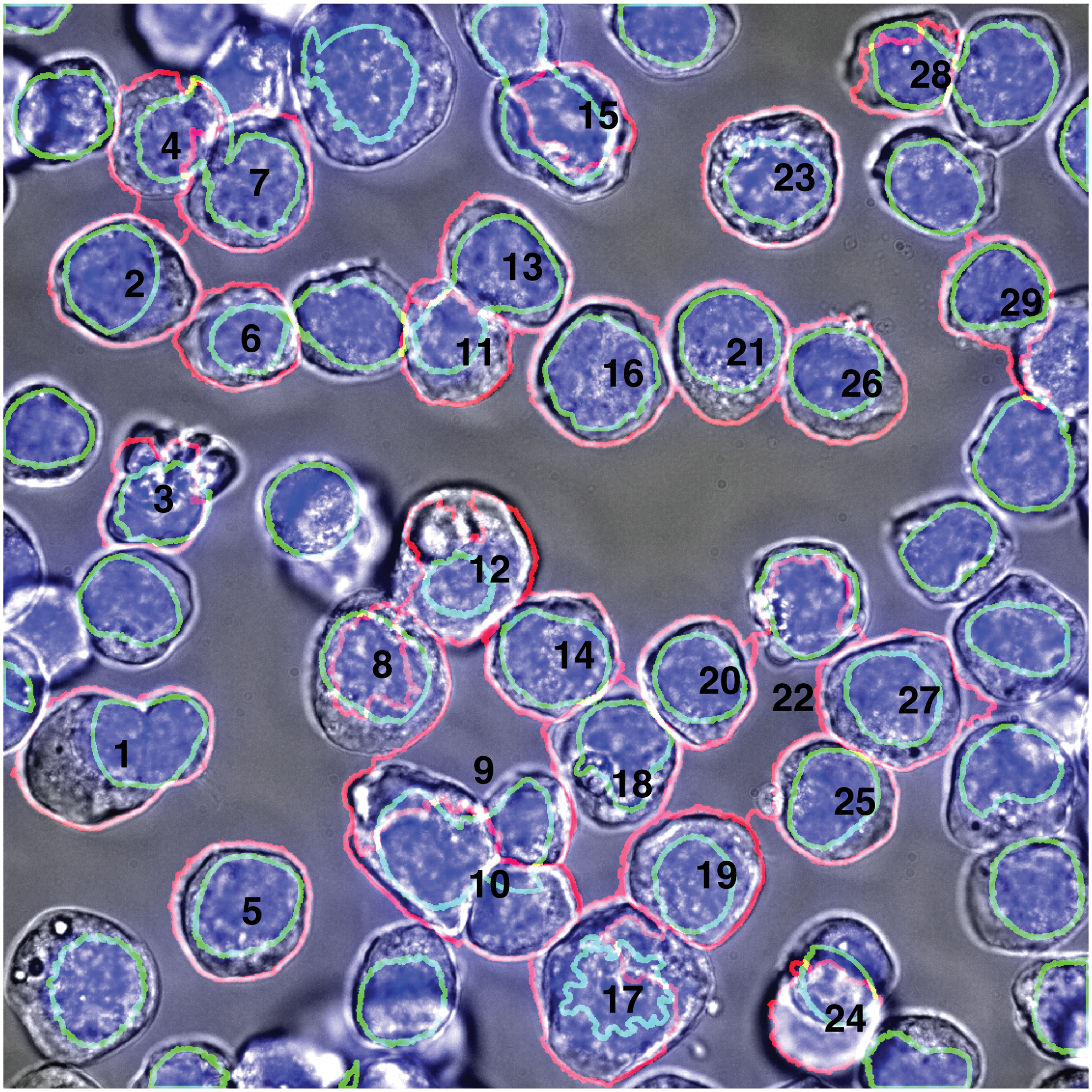


**Supplementary Fig. 12. Example Image of segmented cells from Supplementary Fig. 11.** The brightfield image is displayed in grey. The nucleus is stained with DAPI and appears in blue. The segmented nucleus is highlighted in green, while the segmented cell boundary is outlined in red. Black numbers are used to identify the segmented cells. Cells located on the border of the image and cells with only nuclear segmentation are not included in the analysis.

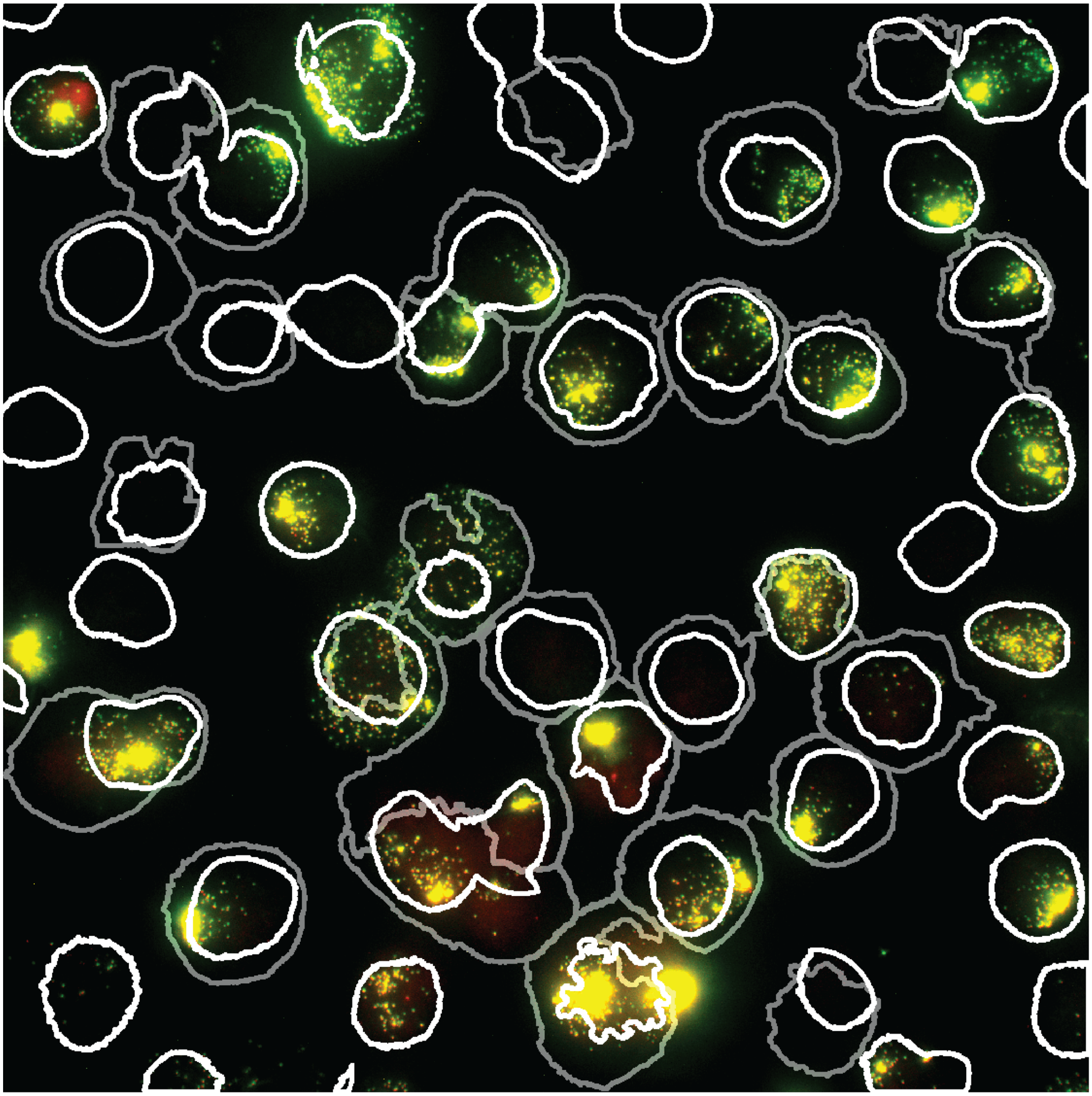

**Supplementary Fig. 13. Example Image segmented cells expressing Xist and RepA using exonic probes from Supplementary Fig. 11.** This image shows an RNA-FISH signal in mouse embryonic stem cells (mESCs) differentiated for 2 days. The Xist exonic probe signal is shown in green, and the RepA exonic probe signal is shown in red, visualizing both mature transcripts and the site of transcription. Colocalization of Xist and RepA signals is displayed in yellow. The nuclear segmentation boundary is marked in white, and the cell segmentation boundary is marked in grey.

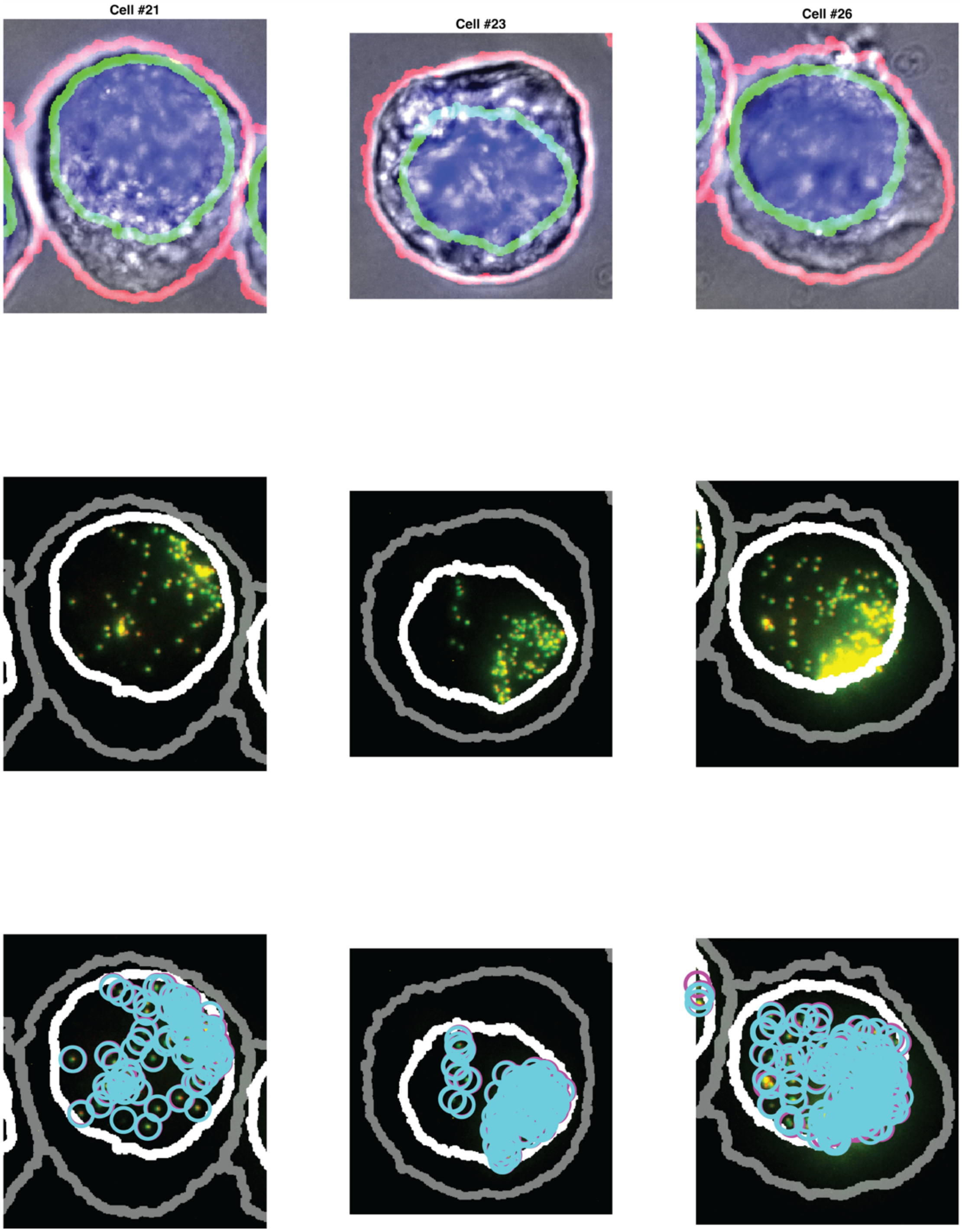


**Supplementary Fig. 14. Example Cells from Supplementary Fig. 12 Demonstrating Cell Segmentation, Nuclear Segmentation, and RNA Spot Counting.** The top panel shows nuclear and cell segmentation. The middle panel displays the RNA-FISH signal for mature Xist transcripts in green and RepA transcripts in red. The bottom panel illustrates RNA spot detection, with Xist transcripts marked by cyan circles and RepA transcripts marked by magenta circles.

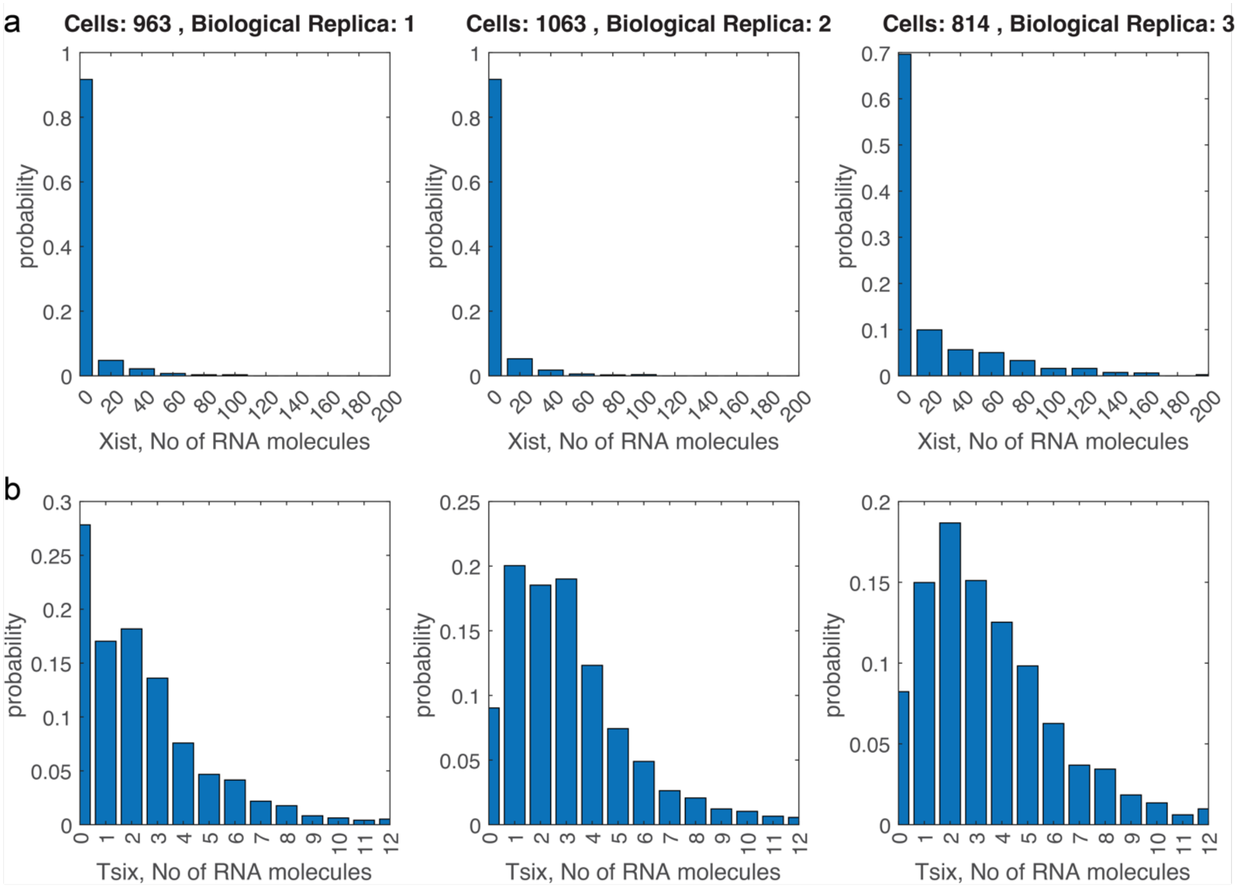


**Supplementary Fig. 15. Detailed distributions from Fig. 3. a-b)** Marginal probability distributions for each of the replica experiments of exonic Xist (a) and exonic Tsix (b) using the same binning as in Fig. 3. The bin size is 20 RNA molecules in (a).

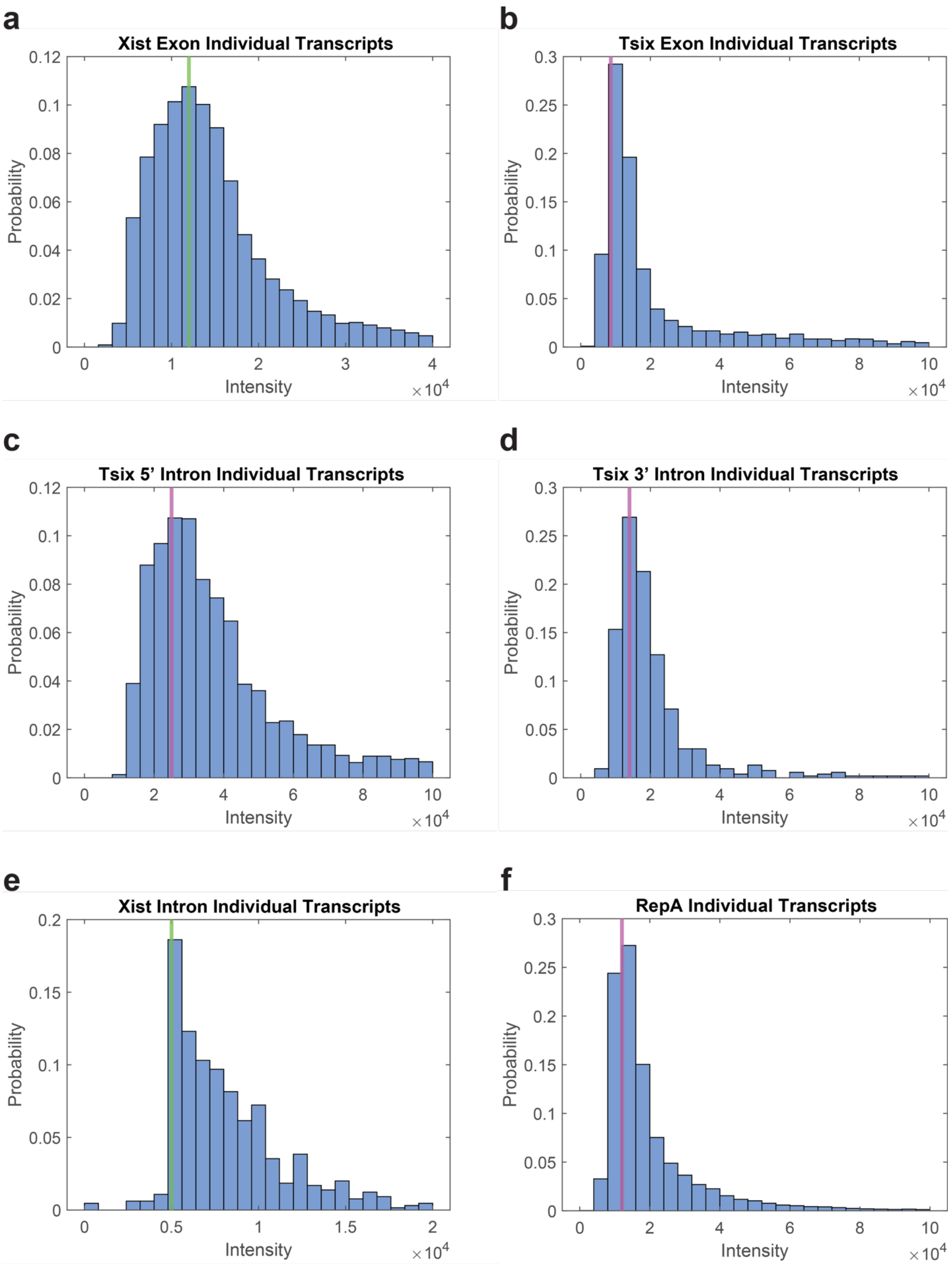


**Supplementary Fig. 16. Probability distribution of the intensities of individual transcripts**.

Histograms that show the probability density of the intensities of individual spots (spots that are not the two brightest in each cell) reveal the intensity of a single spot with the highest probability (green or magenta vertical lines). Shown are example densities for individual transcripts visualized by Xist Exon probes (734 cells, 12762 transcripts) (A), Tsix Exon probes (734 cells, 1560 transcripts) (B), Tsix 5' Intron probes (625 cells, 784 transcripts) (C), Tsix 3' Intron probes (651 cells, 505 transcripts) (D), and Xist Intron probes (651 cells, 194 transcripts) (E).

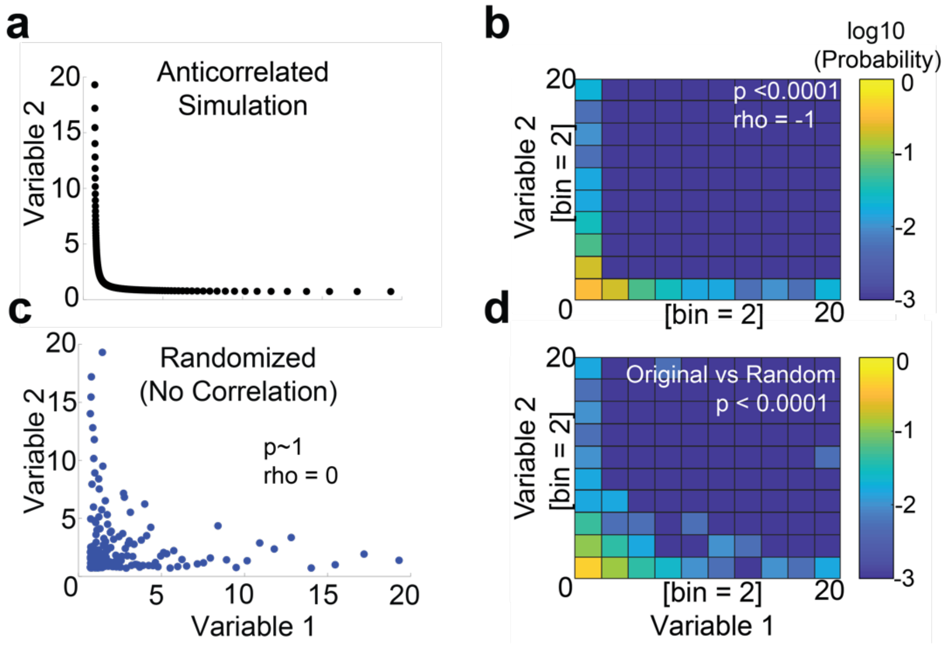


**Supplementary Fig. 17. Quantitative framework for evaluation of correlation using simulated data**. **(A)** An example of a negatively correlated data with an L-shaped expression pattern. **(B)** Visualization of the data with a joint probability distribution, with correlation values assessed by the non-parametric Spearman test of correlation (white text). **(C)** The same data as in A, but with randomized interactions between the variables, shows no correlation when assessed by the Spearman test (black text). **(D)**. A visualization of the randomized data with a joint probability distribution shows differences from the non-randomized distribution (B) that is significantly different when assessed by the 2D Kolmogorov-Smirnov test (white text).

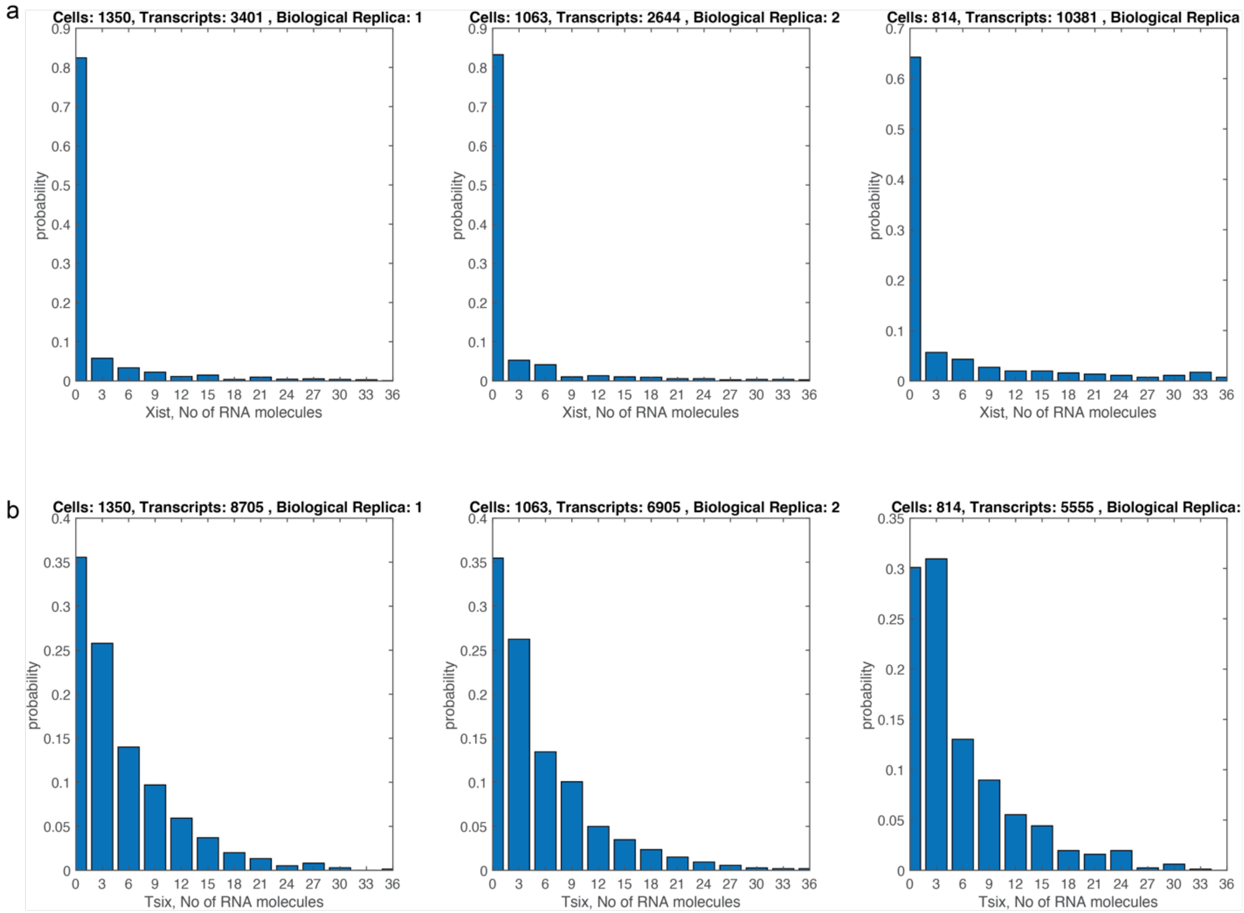


**Supplementary Fig. 18. Detailed distributions from Figure 4. a-b** Marginal probability distributions for each of the replica experiments of nascent Xist (a) and Tsix (b) using exonic probes .

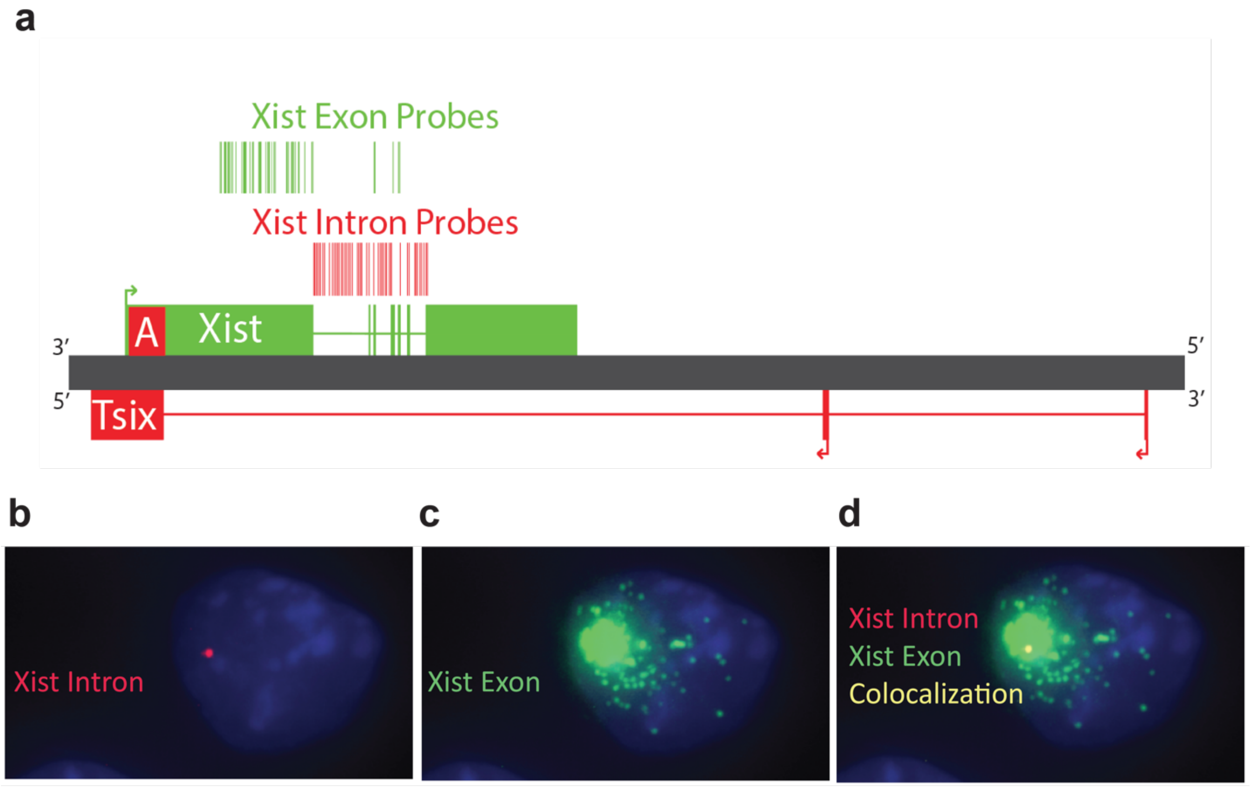


**Supplementary Fig. 19. Intronic probes enrich for nascent transcription**. **a** A diagram (to scale) showing the probe binding locations of Xist exonic probes (green) and Xist intronic probes (red). **b** Example image of a cell with nascent Xist and Xist Intron probe signal (red). **c** Example image of the same cell with mature Xist transcripts visualized by Xist Exon signal (green). **d** The combination of images b and c, with Xist intron signal in red, Xist Exon signal in green, and colocalization in yellow. DAPI stained nucleus in blue.

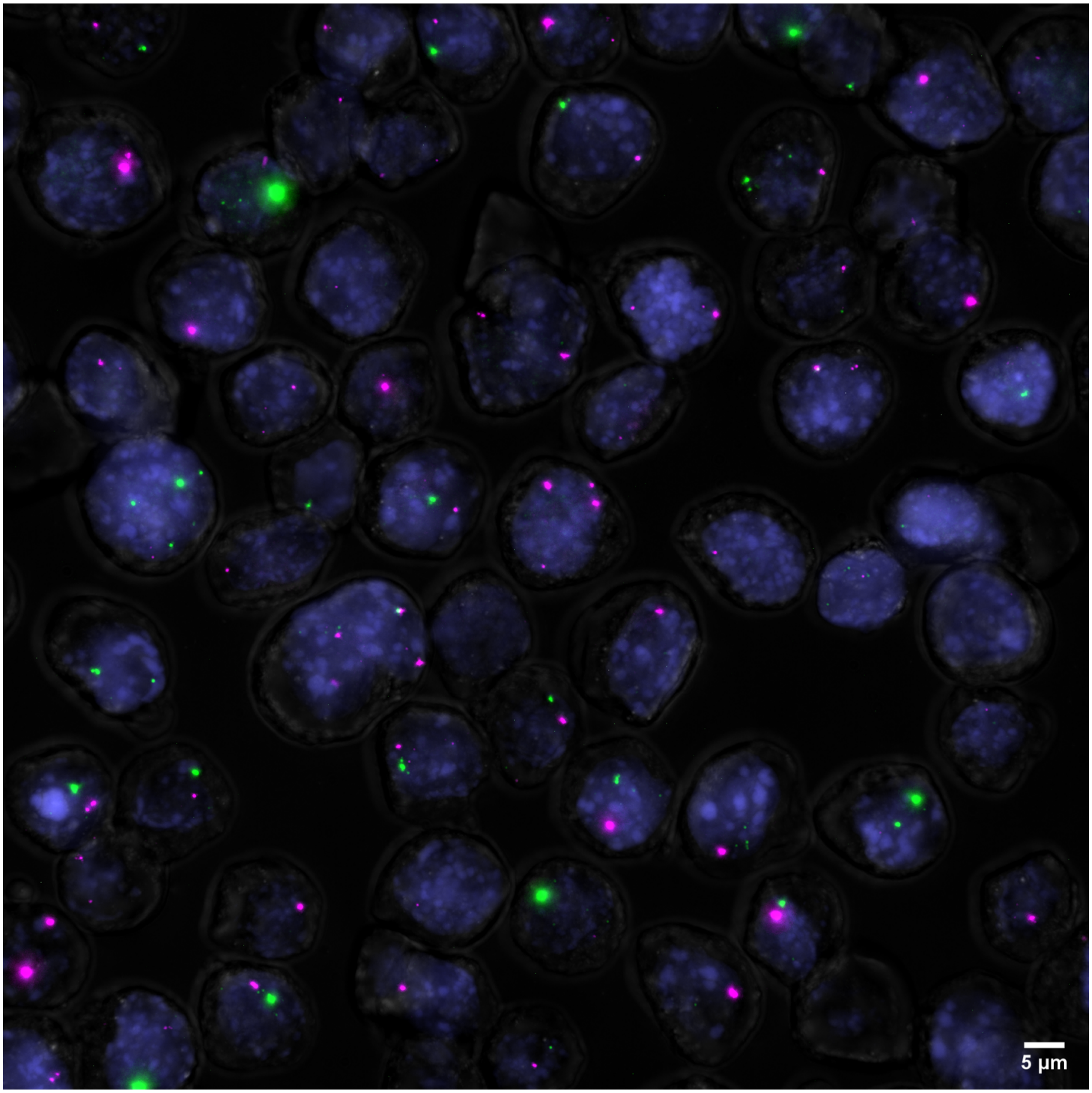


**Supplementary Fig. 20. Example Image of Xist Intronic and Tsix 3' Intronic Probes.** An example maximum intensity projection image of RNA-FISH signal in mESCs differentiated for 2 days. Xist Exonic probe signal (green) and Tsix 3' probe signal (magenta) visualizes both mature transcripts and the site of transcription. Colocalization of Xist and Tsix 3' is shown in white. DAPI stained nucleus in blue.

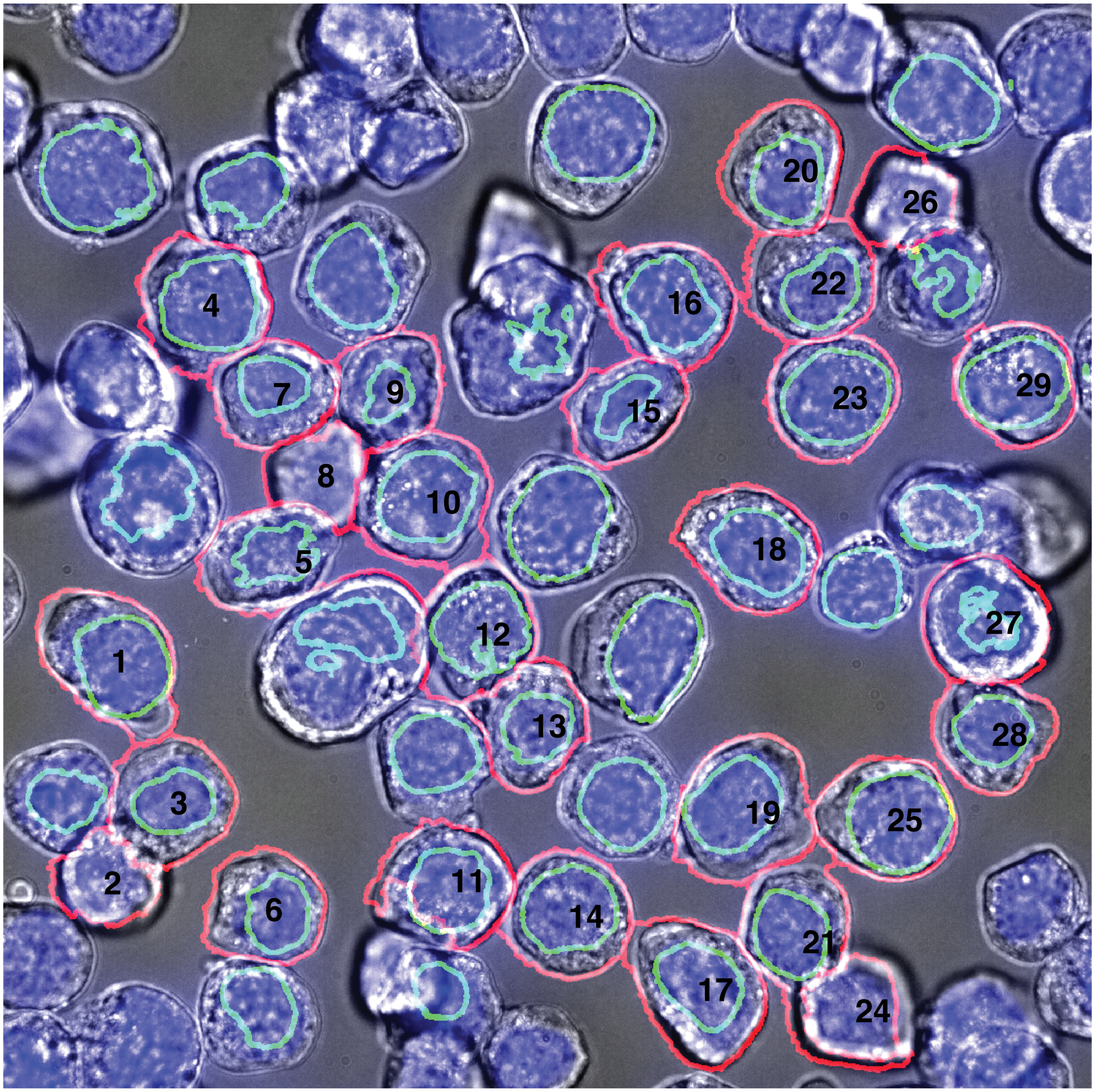


**Supplementary Fig. 21. Example Image of segmented cells from Supplementary Fig. 20.** The brightfield image is displayed in grey. The nucleus is stained with DAPI and appears in blue. The segmented nucleus is highlighted in green, while the segmented cell boundary is outlined in red. Black numbers are used to identify the segmented cells. Cells located on the border of the image and cells with only nuclear segmentation are not included in the analysis.

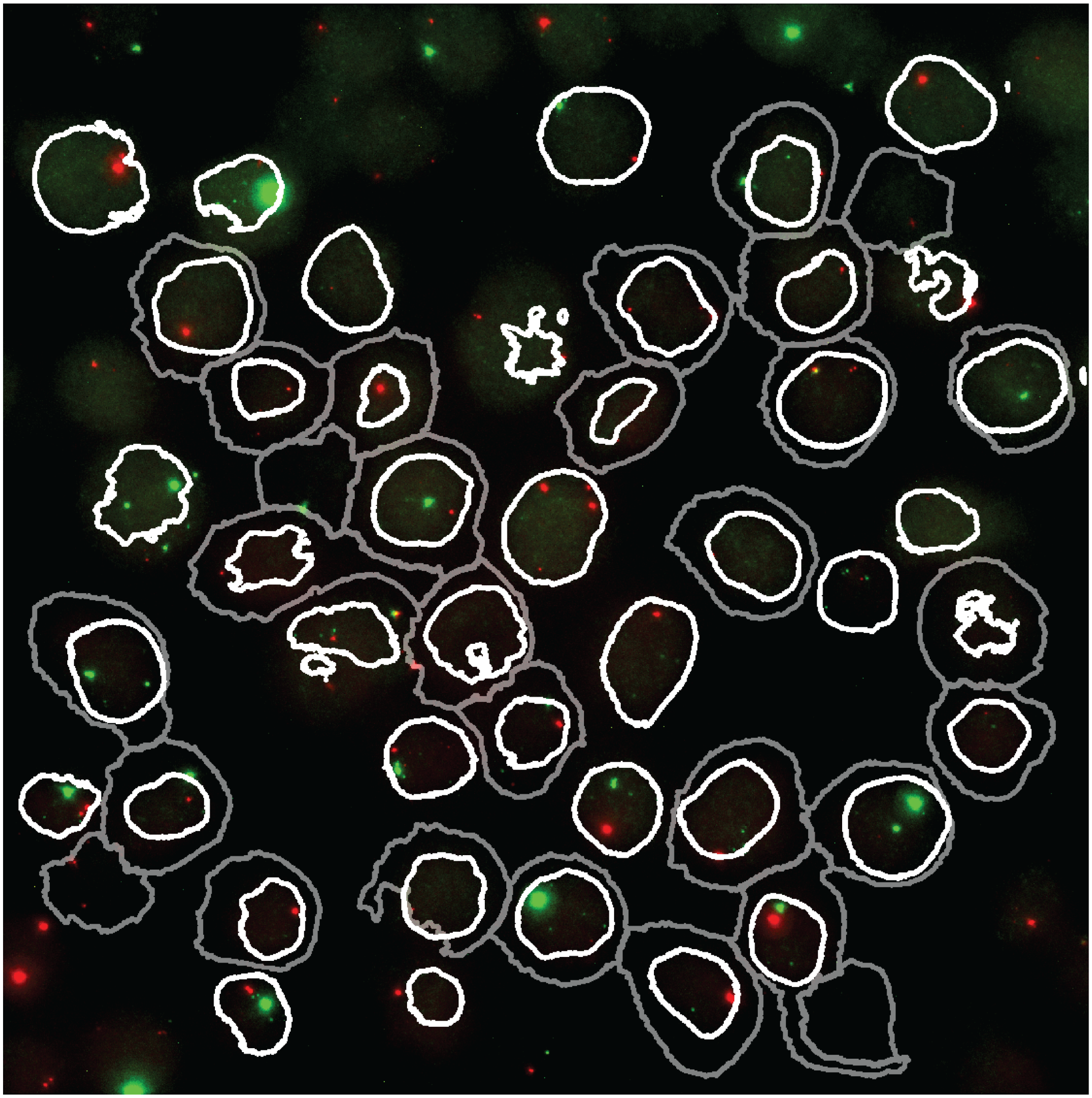

**Supplementary Fig. 22. Example Image segmented cells expressing Xist and Tsix 3' using Intronic probes from Supplementary Fig. 20.** This image shows an RNA-FISH signal in mouse embryonic stem cells (mESCs) differentiated for 2 days. The Xist intronic probe signal is shown in green, and the Tsix 3' intronic probe signal is shown in red, visualizing both mature transcripts and the site of transcription. Colocalization of Xist and Tsix 3' signals is displayed in yellow. The nuclear segmentation boundary is marked in white, and the cell segmentation boundary is marked in grey.

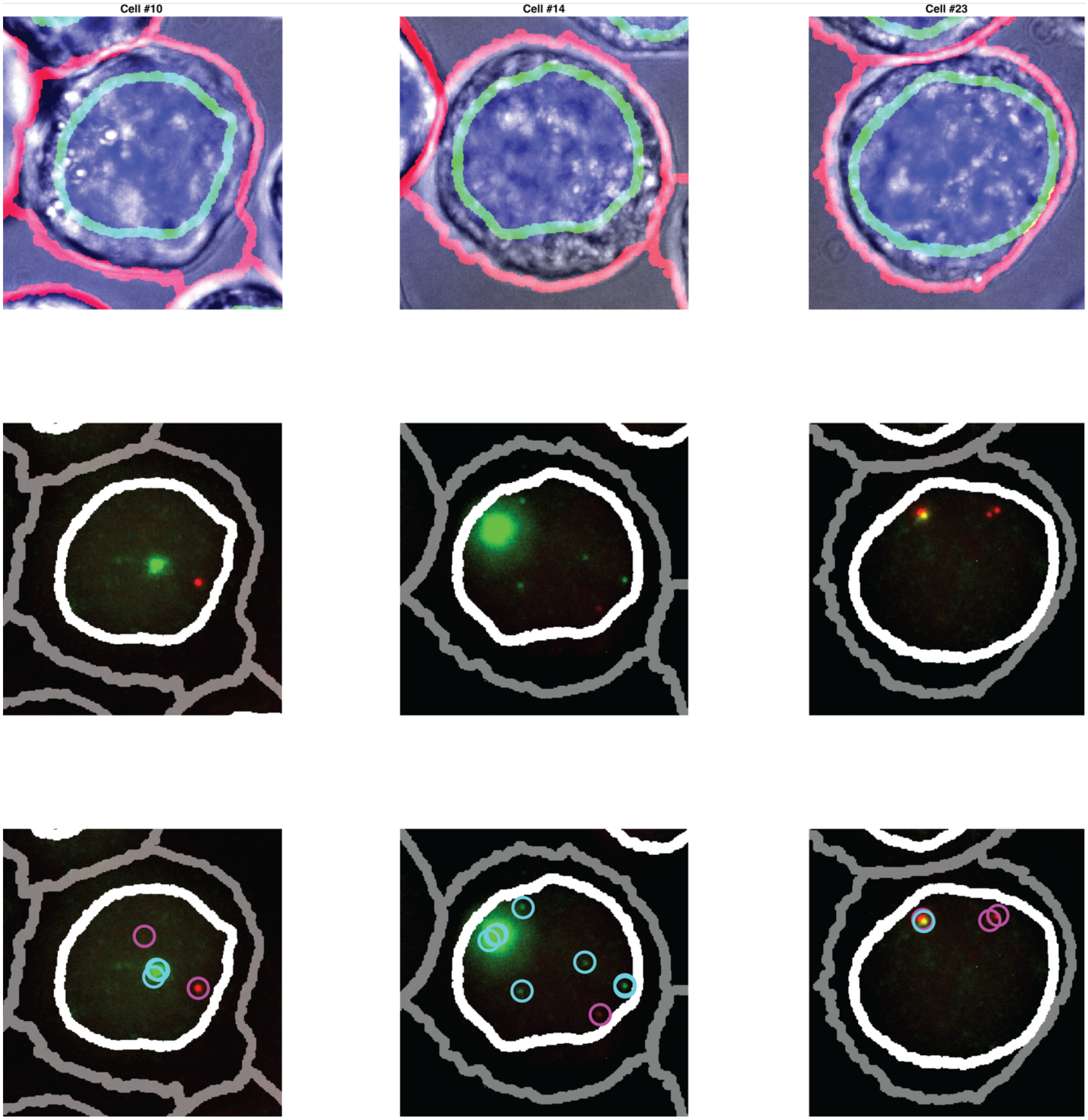


**Supplementary Fig. 23. Example Cells from Supplementary Fig. 21 Demonstrating Cell Segmentation, Nuclear Segmentation, and RNA Spot Counting.** The top panel shows nuclear and cell segmentation. The middle panel displays the RNA-FISH signal for mature Xist transcripts in green and Tsix 3’ transcripts in red. The bottom panel illustrates RNA spot detection, with Xist transcripts marked by cyan circles and Tsix 3’ transcripts marked by magenta circles.

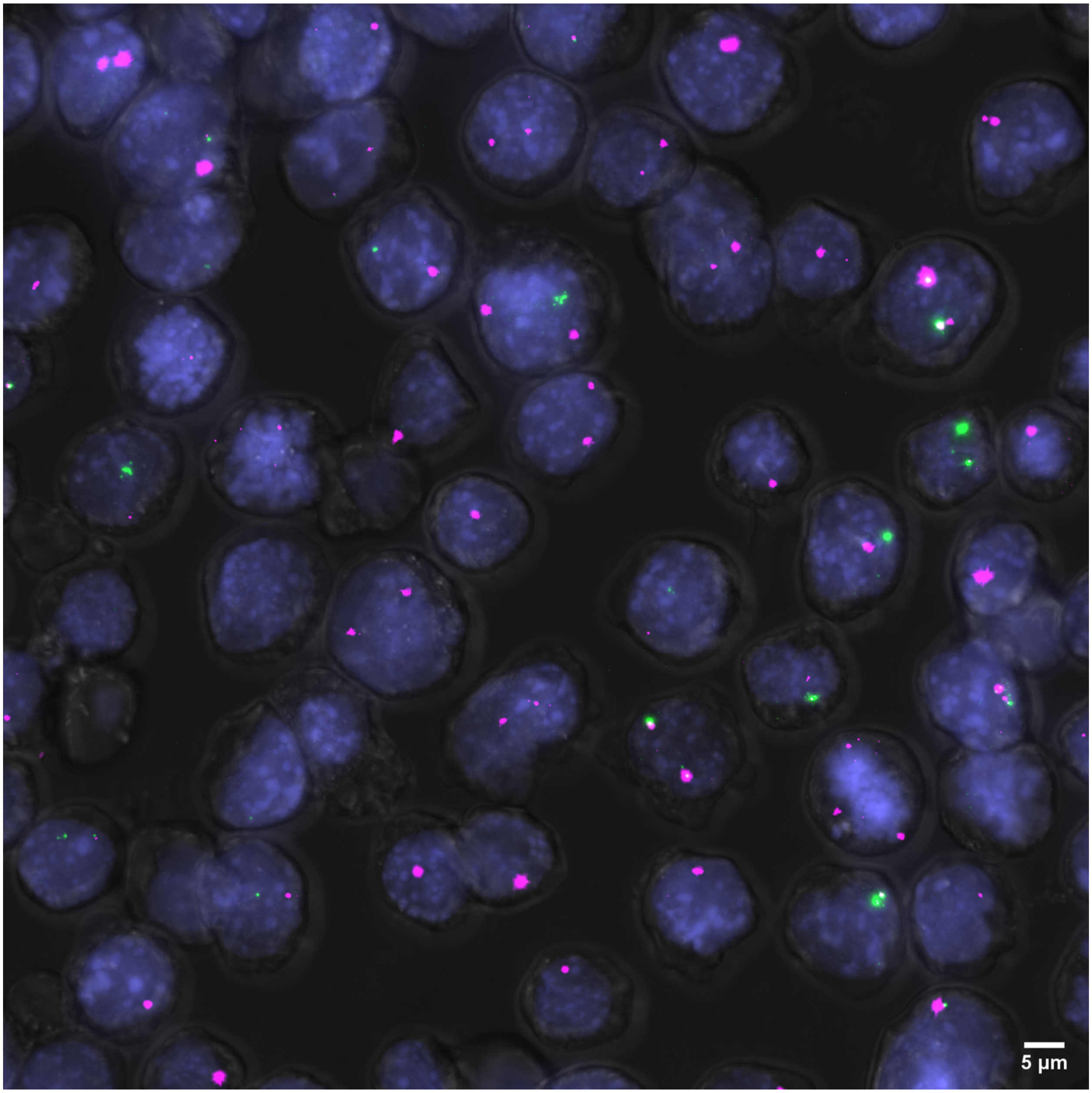


**Supplementary Fig. 24. Example Image of Xist Intronic and Tsix 5’ Intronic Probes.** An example maximum intensity projection image of RNA-FISH signal in mESCs differentiated for 2 days. Xist Exonic probe signal (green) and Tsix 5’ intronic probe signal (magenta) visualizes both mature transcripts and the site of transcription. Colocalization of Xist and Tsix 5’ signal is shown in white. DAPI stained nucleus in blue.

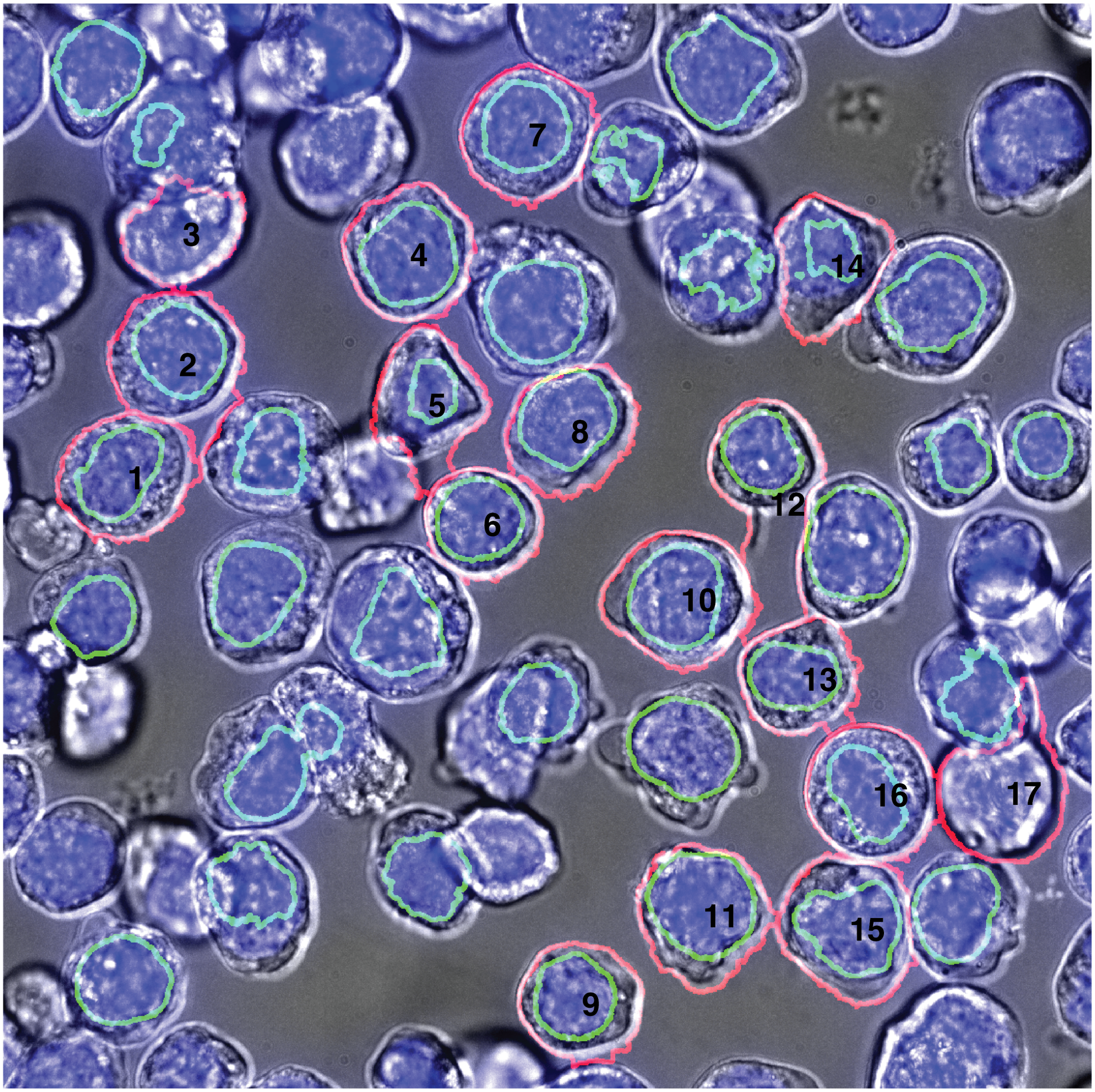


**Supplementary Fig. 25. Example Image of segmented cells from Supplementary Fig. 24.** The brightfield image is displayed in grey. The nucleus is stained with DAPI and appears in blue. The segmented nucleus is highlighted in green, while the segmented cell boundary is outlined in red. Black numbers are used to identify the segmented cells. Cells located on the border of the image and cells with only nuclear segmentation are not included in the analysis.

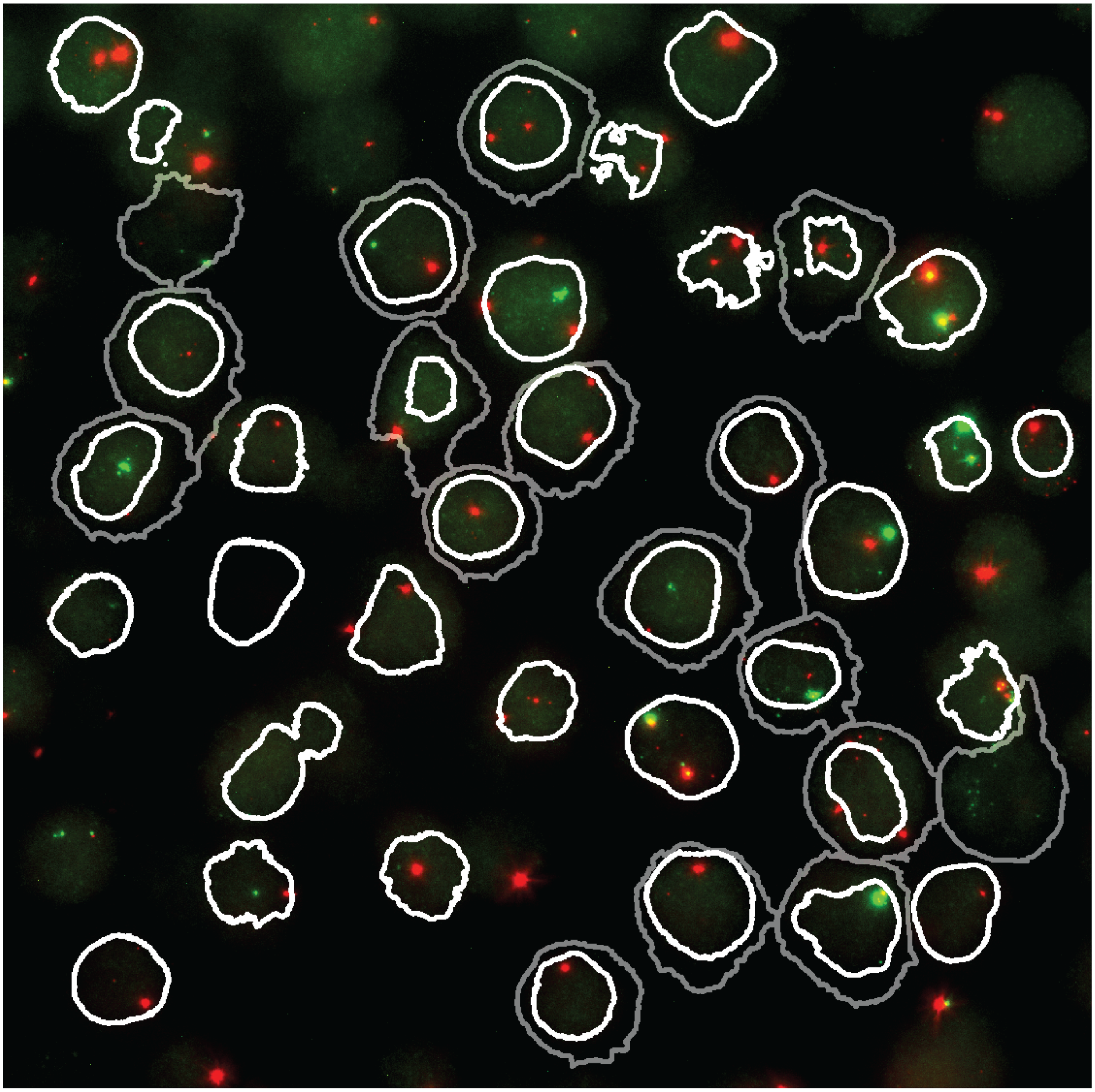

**Supplementary Fig. 26. Example Image segmented cells expressing Xist and Tsix 3’ using Intronic probes from Supplementary Fig. 24.** This image shows an RNA-FISH signal in mouse embryonic stem cells (mESCs) differentiated for 2 days. The Xist intronic probe signal is shown in green, and the Tsix 5’ intronic probe signal is shown in red, visualizing both mature transcripts and the site of transcription. Colocalization of Xist and Tsix 5’ signals is displayed in yellow. The nuclear segmentation boundary is marked in white, and the cell segmentation boundary is marked in grey.

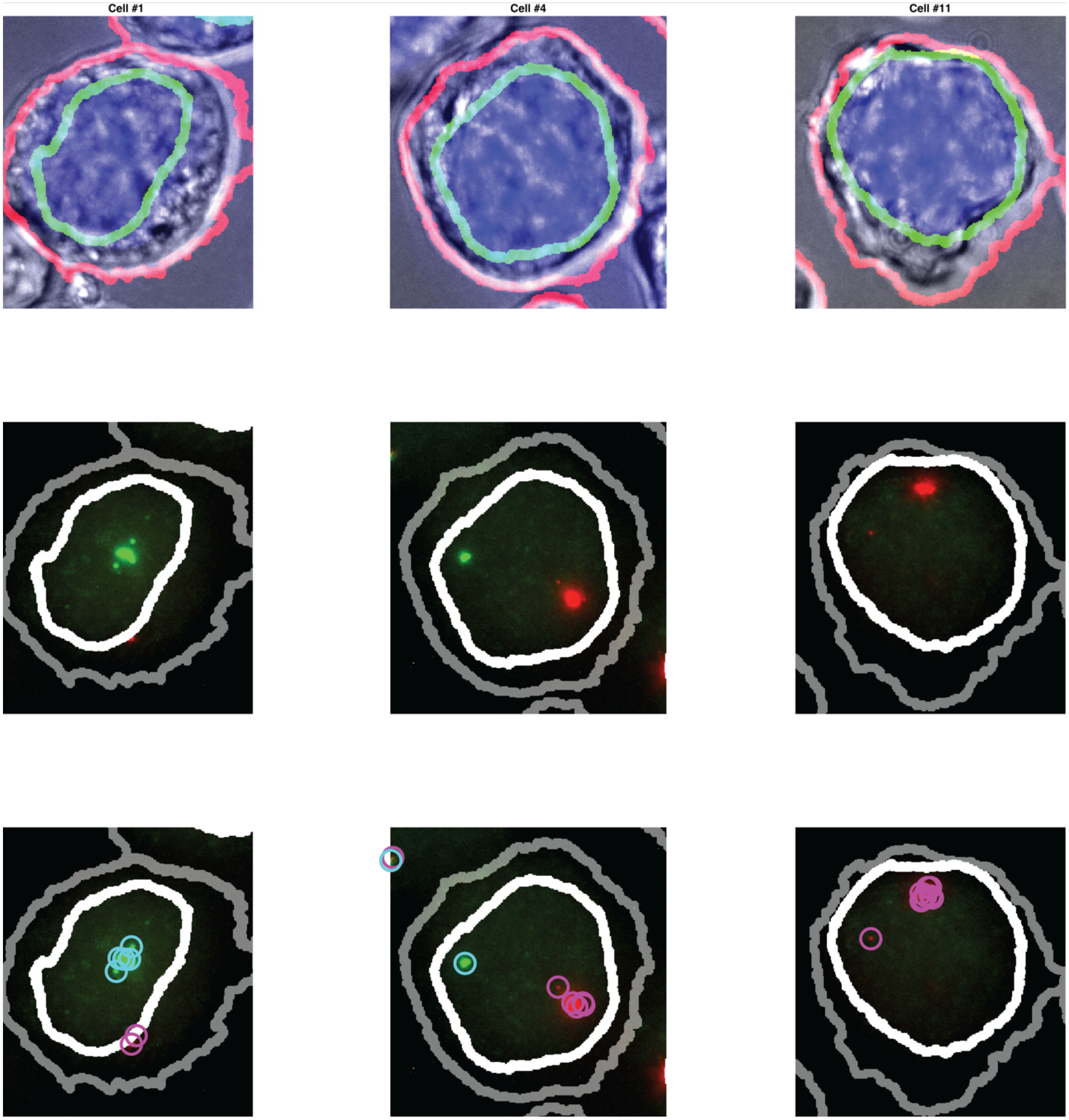


**Supplementary Fig. 27. Example Cells from Supplementary Fig. 25 Demonstrating Cell Segmentation, Nuclear Segmentation, and RNA Spot Counting.** The top panel shows nuclear and cell segmentation. The middle panel displays the RNA-FISH signal for mature Xist transcripts in green and Tsix 5' transcripts in red. The bottom panel illustrates RNA spot detection, with Xist transcripts marked by cyan circles and Tsix 5' transcripts marked by magenta circles.

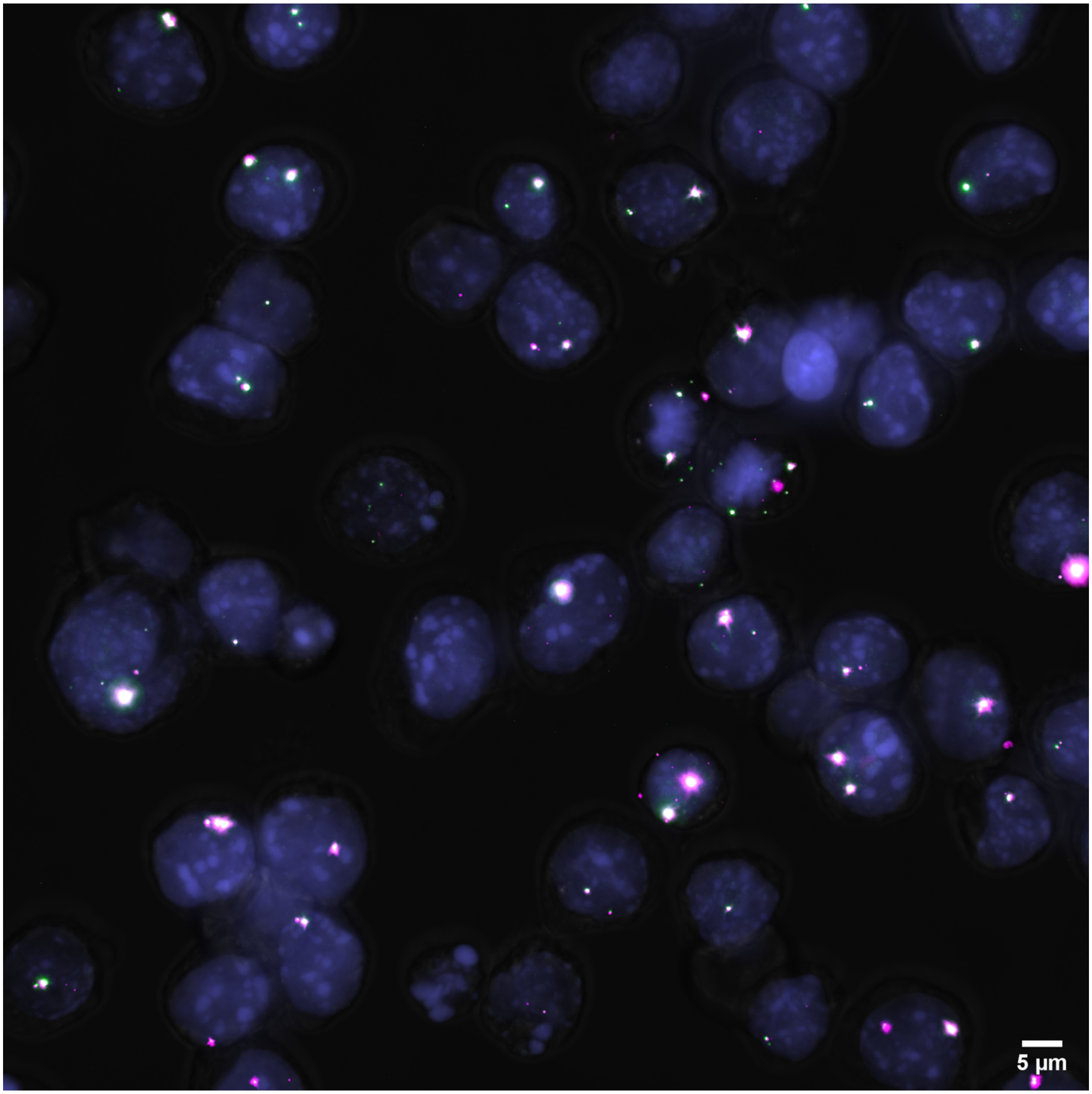


**Supplementary Fig. 28. Example Image of Tsix 5' Intronic and Tsix 3' Intronic Probes.** An example image of RNA-FISH signal in mESCs differentiated for 2 days. Tsix 3' intronic probe signal (green) and Tsix 5' intronic probe signal (magenta) visualizes both mature transcripts and the site of transcription. Colocalization of Tsix 5' and 3' is shown in white. DAPI stained nucleus in blue.

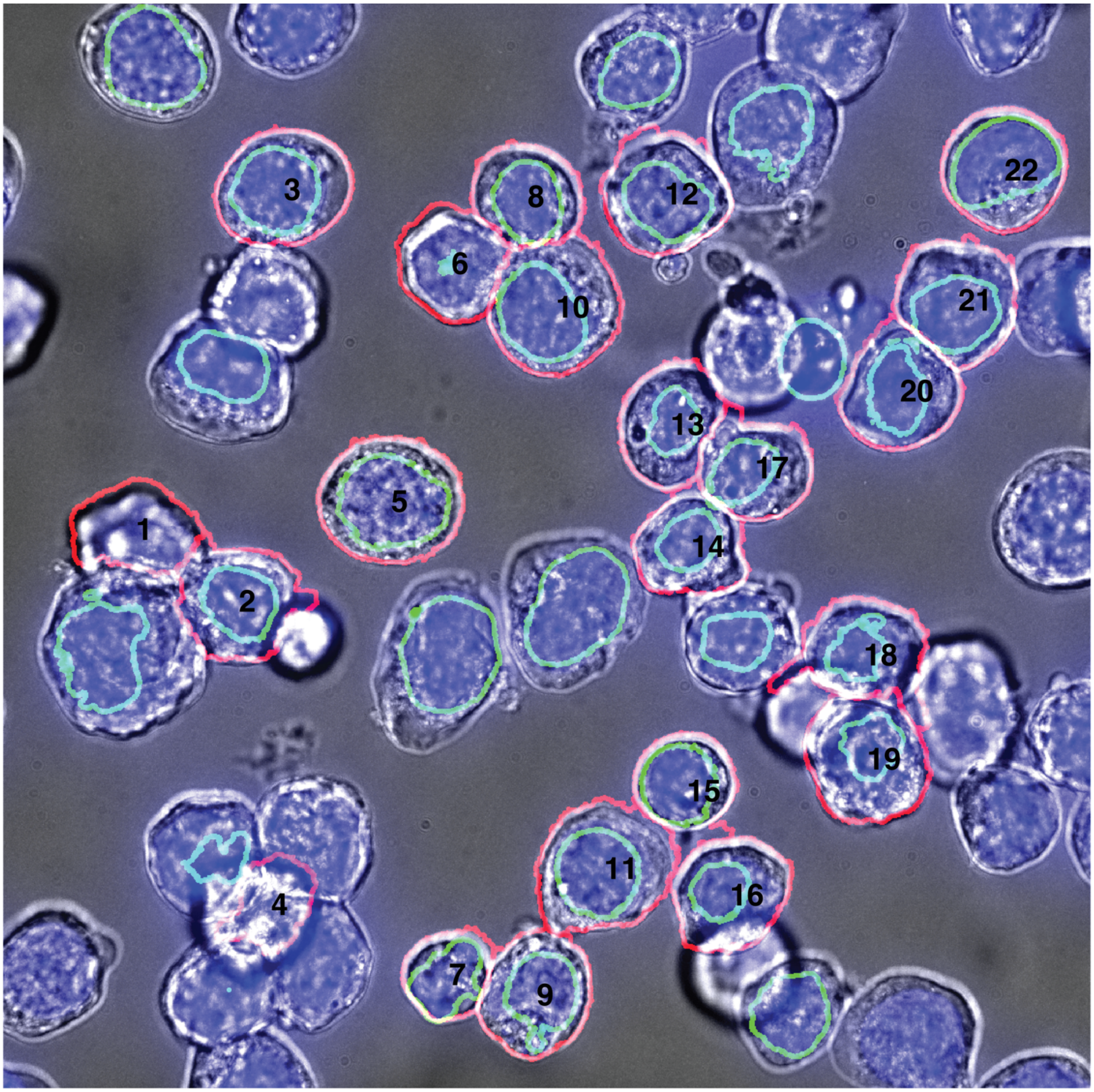


**Supplementary Fig. 29. Example Image of segmented cells from Supplementary Fig. 29.** The brightfield image is displayed in grey. The nucleus is stained with DAPI and appears in blue. The segmented nucleus is highlighted in green, while the segmented cell boundary is outlined in red. Black numbers are used to identify the segmented cells. Cells located on the border of the image and cells with only nuclear segmentation are not included in the analysis.

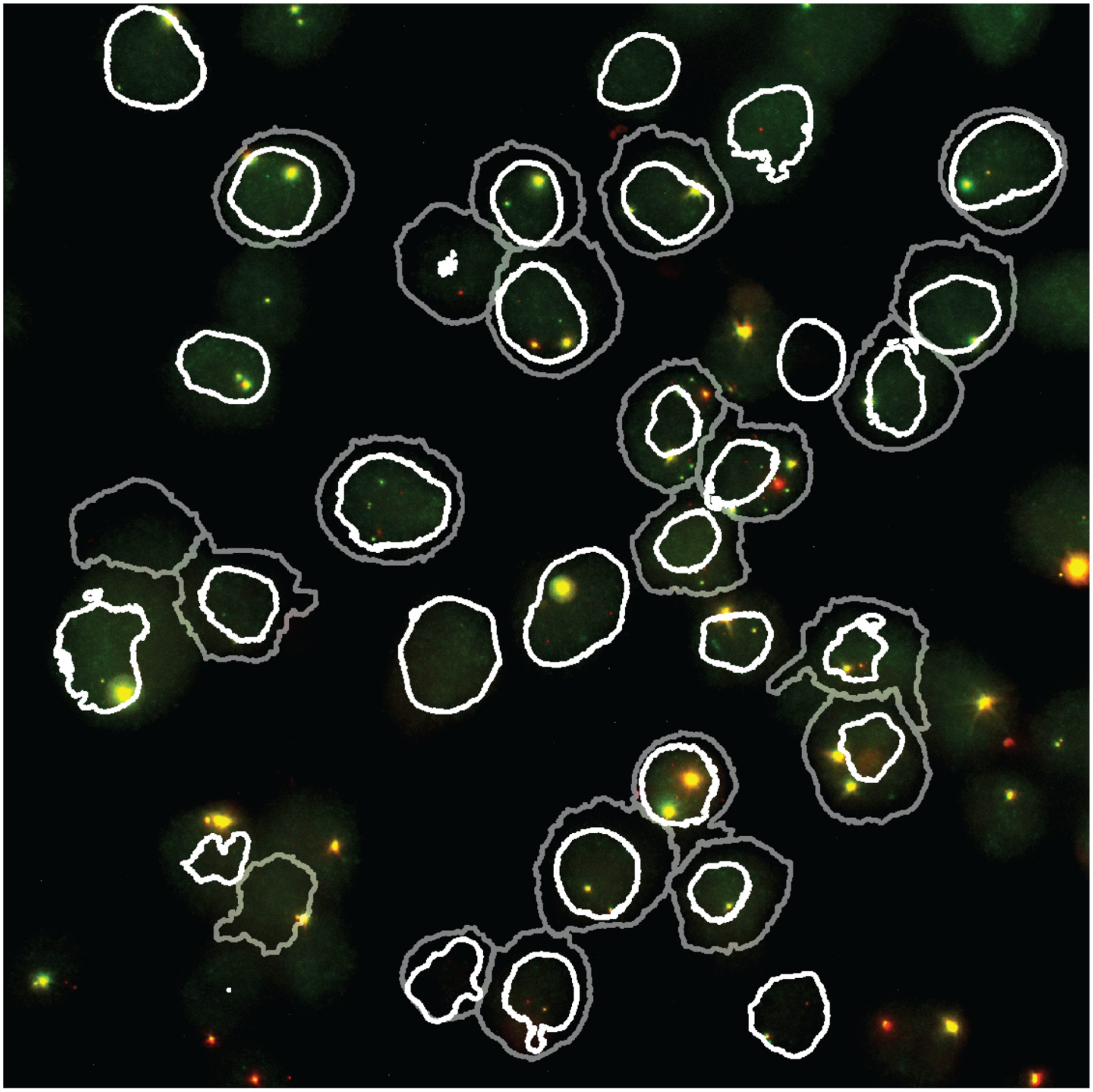

**Supplementary Fig. 30. Example Image segmented cells expressing Tsix 5’ and Tsix 3’ using Intronic probes from Supplementary Fig. 28.** This image shows an RNA-FISH signal in mouse embryonic stem cells (mESCs) differentiated for 2 days. The Tsix 5’ intronic probe signal is shown in green, and the Tsix 3’ intronic probe signal is shown in red, visualizing both mature transcripts and the site of transcription. Colocalization of Tsix 5’ and Tsix 3’ signals is displayed in yellow. The nuclear segmentation boundary is marked in white, and the cell segmentation boundary is marked in grey.

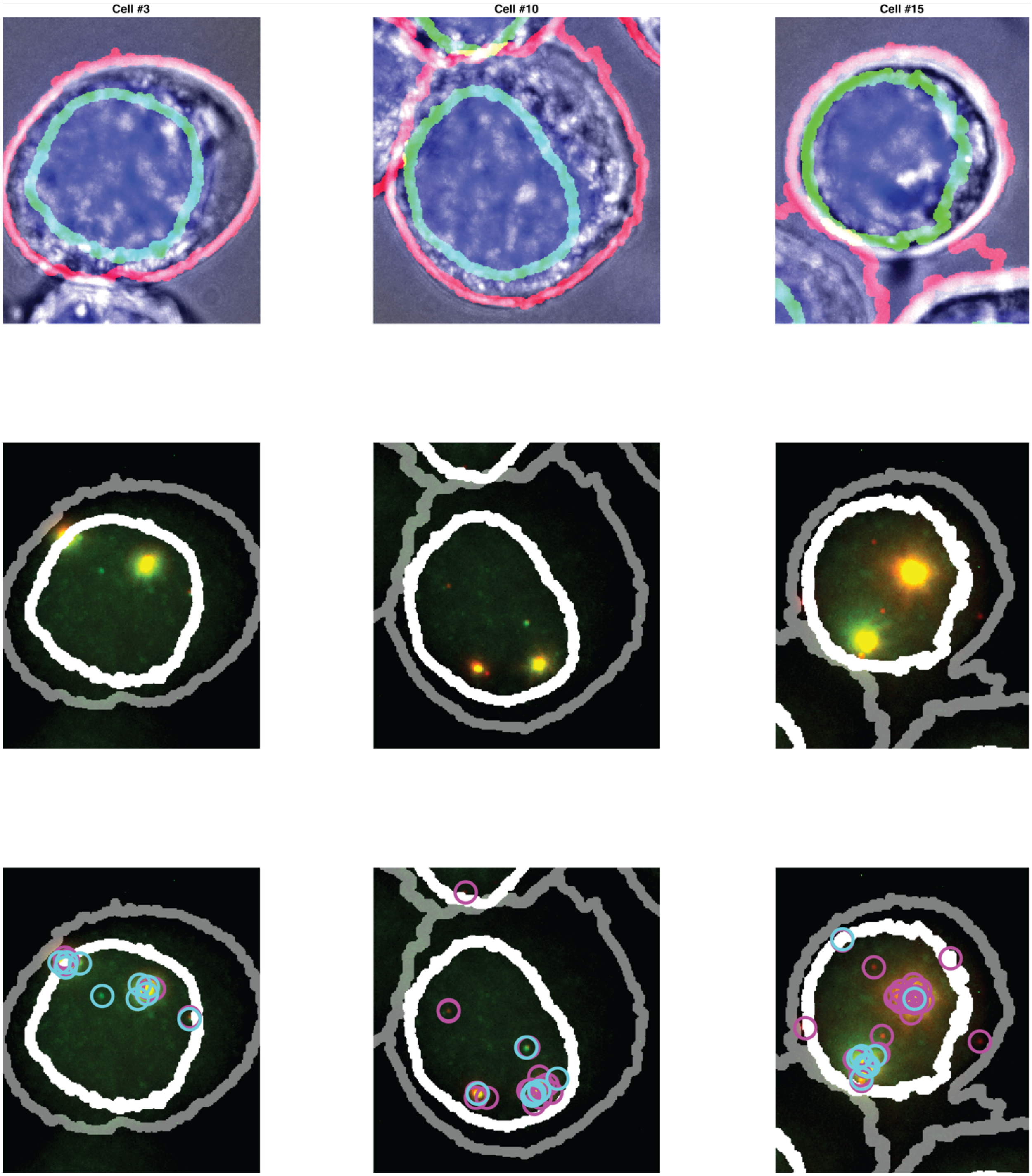


**Supplementary Fig. 31. Example Cells from Supplementary Fig. 29 Demonstrating Cell Segmentation, Nuclear Segmentation, and RNA Spot Counting.** The top panel shows nuclear and cell segmentation. The middle panel displays the RNA-FISH signal for mature Tsix 5' transcripts in green and Tsix 3' transcripts in red. The bottom panel illustrates RNA spot detection, with Tsix5’ transcripts marked by cyan circles and Tsix 3’ transcripts marked by magenta circles.

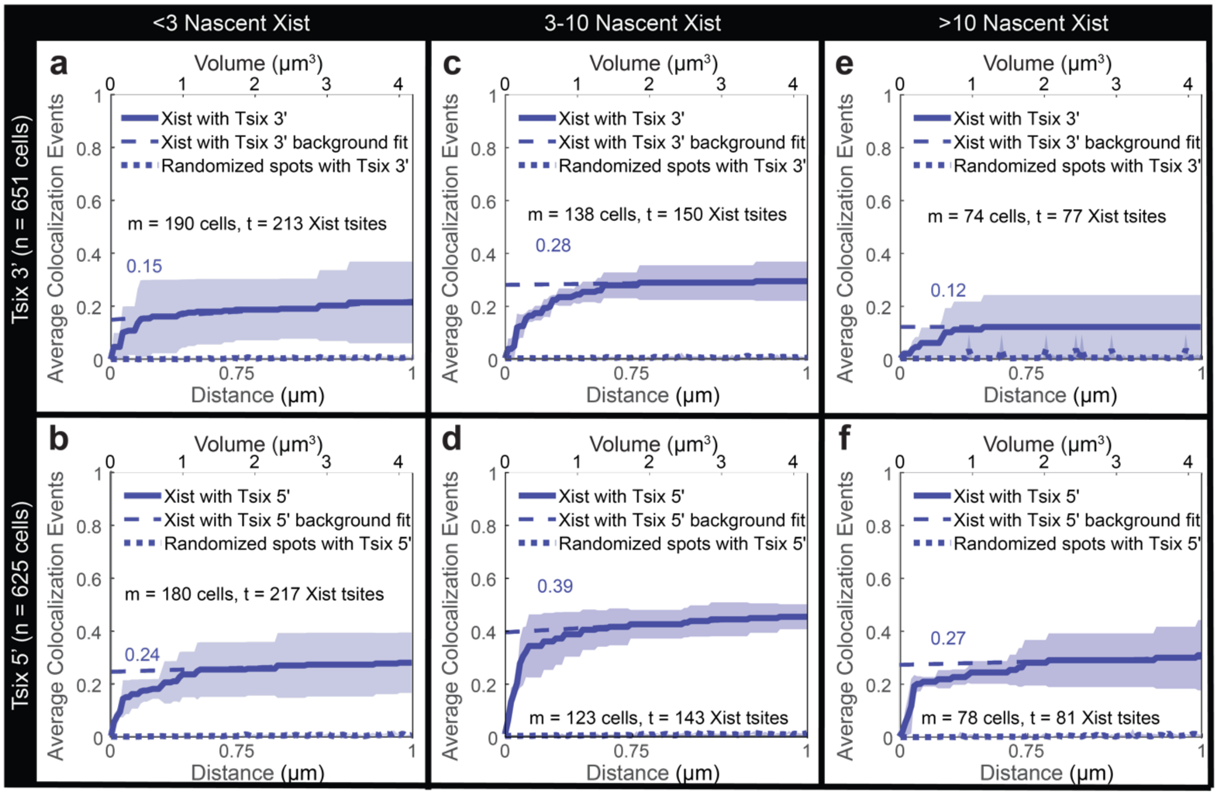
<3 Nascent Xist
3-10 Nascent Xist
>10 Nascent Xist
Tsix 3' (n = 651 cells)
Tsix 5' (n = 625 cells)
a
Volume (μm$^3$)
Average Colocalization Events
Distance (μm)
Xist with Tsix 3'
Xist with Tsix 3' background fit
Randomized spots with Tsix 3'
m = 190 cells, t = 213 Xist tsites
0.15
c
m = 138 cells, t = 150 Xist tsites
0.28
e
m = 74 cells, t = 77 Xist tsites
0.12
b
Xist with Tsix 5'
Xist with Tsix 5' background fit
Randomized spots with Tsix 5'
m = 180 cells, t = 217 Xist tsites
0.24
d
m = 123 cells, t = 143 Xist tsites
0.39
f
m = 78 cells, t = 81 Xist tsites
0.27
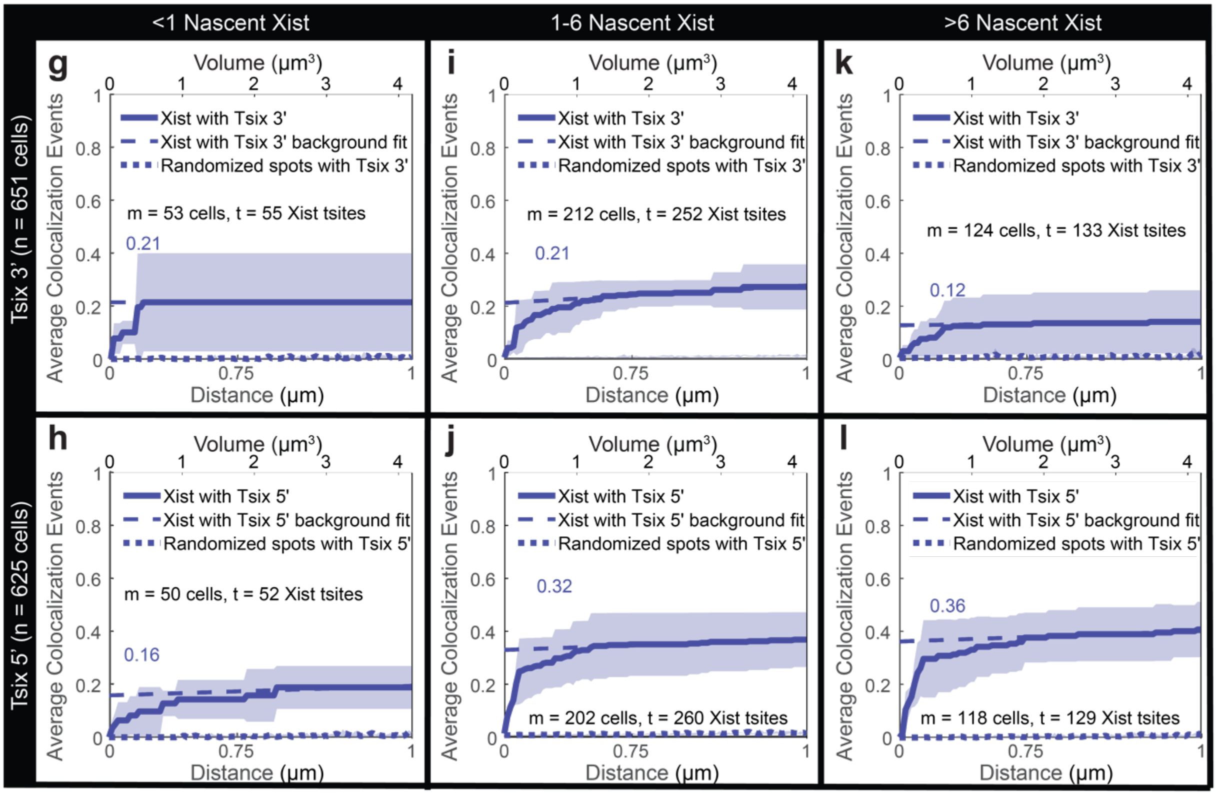
<1 Nascent Xist
1-6 Nascent Xist
>6 Nascent Xist
Tsix 3' (n = 651 cells)
Tsix 5' (n = 625 cells)
g
Volume (μm$^3$)
Average Colocalization Events
Distance (μm)
Xist with Tsix 3'
Xist with Tsix 3' background fit
Randomized spots with Tsix 3'
m = 53 cells, t = 55 Xist tsites
0.21
i
m = 212 cells, t = 252 Xist tsites
0.21
k
m = 124 cells, t = 133 Xist tsites
0.12
h
Xist with Tsix 5'
Xist with Tsix 5' background fit
Randomized spots with Tsix 5'
m = 50 cells, t = 52 Xist tsites
0.16
j
m = 202 cells, t = 260 Xist tsites
0.32
l
m = 118 cells, t = 129 Xist tsites
0.36

**Supplementary Fig. 32. Quantification of Colocalization at Sites of Transcription (tsites) at Different Cutoffs. a-f** show the same data as in Figure 5i-n but with a higher cutoff for low (<3 transcripts), medium (3 to 10 transcripts), and high (>10 transcripts) nascent Xist expression. **g-l** display the same data as in Figure 5i-n but with a lower cutoff for low (<1 transcript), medium (2 to 6 transcripts), and high (>6 transcripts) nascent intronic Xist expression. The results are independent of the cutoffs.

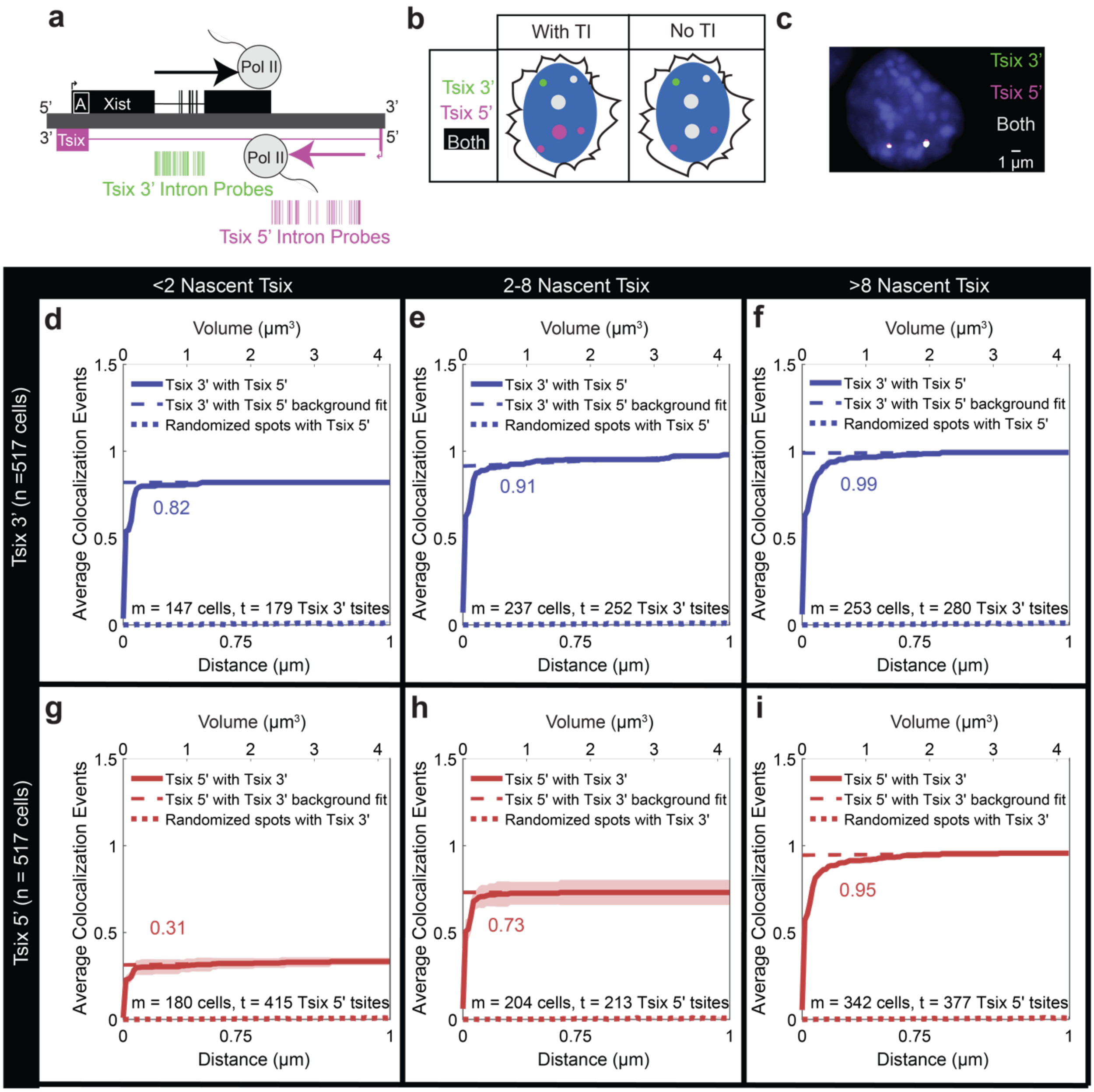


**Supplementary Fig. 33. Colocalization of signal from Tsix 5' and Tsix 3' intronic probes shows incomplete processivity**. **a** A to-scale diagram showing the locations of Tsix 5' intron probes (magenta) and Tsix 3' intron probes (green). **b** A conceptual diagram showing transcription sites (large circles) and individual molecules (small circles) of Tsix 5' signal (magenta) and Tsix 3' signal (green). These show the expected results with TI (left) and the increased colocalization without TI (right). **c** An example microscopy image showing Tsix 5'

signal (magenta), Tsix 3’ signal (green), and colocalization (white). **d-i** Quantification of colocalization at sites of transcription. On the y-axis is the total average colocalization events per transcription site. On the x-axis is the search radius (in µm) from the transcription site. This axis is on a cubic scale, so volume increases linearly (top x-axis). On the left column, n = total cells quantified in the experiment, within the panels m = total cells with the specified amount of nascent transcripts and t = the total transcription sites (tsites) with the specified amount of nascent transcripts. **d-f** Quantification of colocalization of Tsix 3’ with Tsix 5' depending on the nascent transcription of Tsix 3’. **g-i** Quantification of colocalization of Tsix 5' with Tsix 3’ depending on the nascent transcription of Tsix 5’. Low-level nascent Tsix 5’ (g), shows low colocalization with Tsix 3’, indicating low processivity. A greater percentage of medium-level Tsix 5’ colocalizes with Tsix 3’ (h), and a very high percentage of highly transcribing Tsix 5’ colocalizes with Tsix 3’ (i).

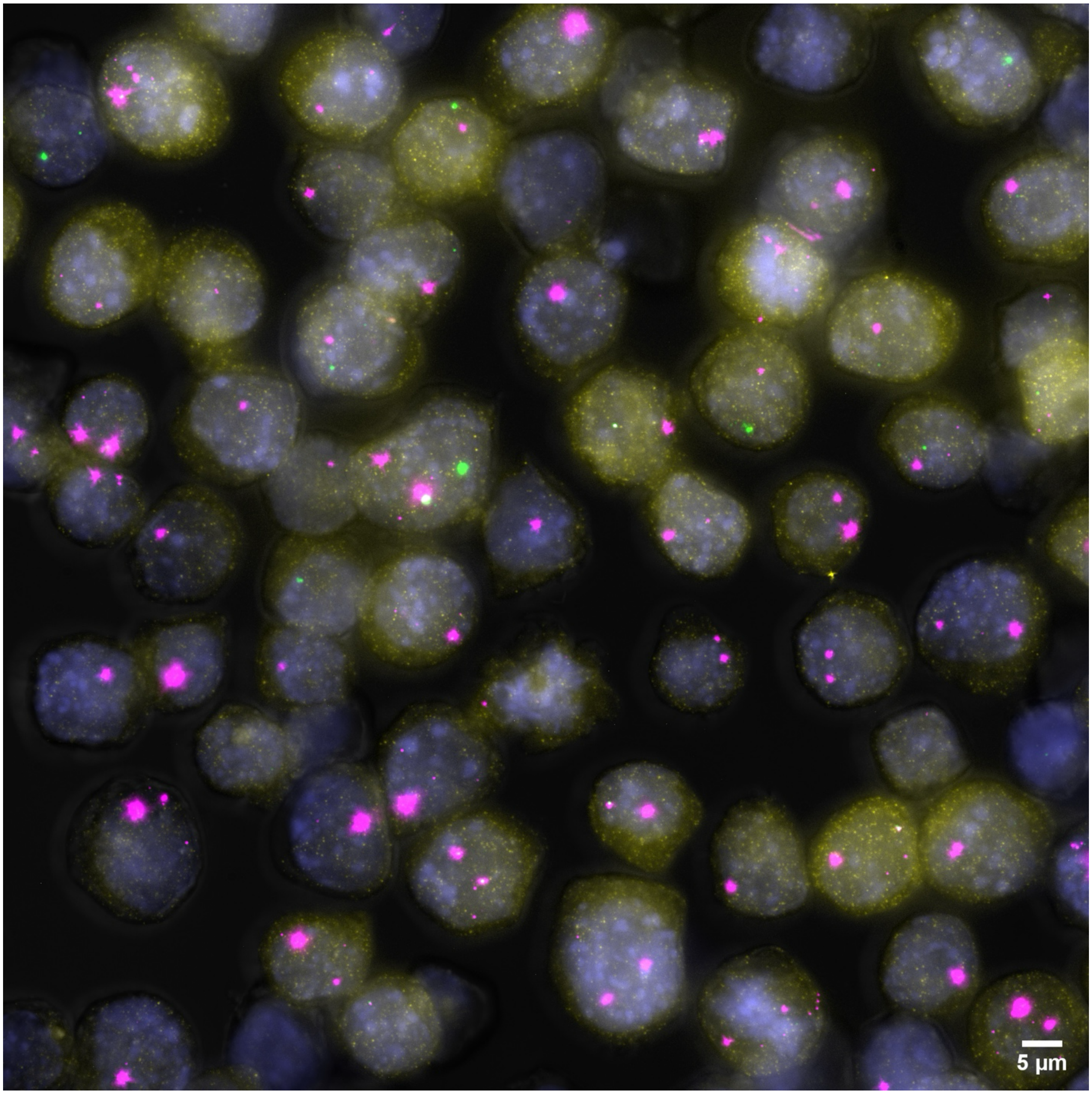


**Supplementary Fig. 34. Example Image of Xist Intronic Signal, Tsix 5' Intronic Signal, and H3K36me3 immunofluorescence.** An example image of RNA-FISH signal in mESCs differentiated for 2 days. Xist Intronic probe signal (green) and Tsix 5' intronic probe signal (magenta) visualizes the site of transcription. H3K36me3 immunofluorescence is shown in yellow. Colocalization of Xist, Tsix 5', and H3K36me3 is shown in white. DAPI stained nucleus in blue.

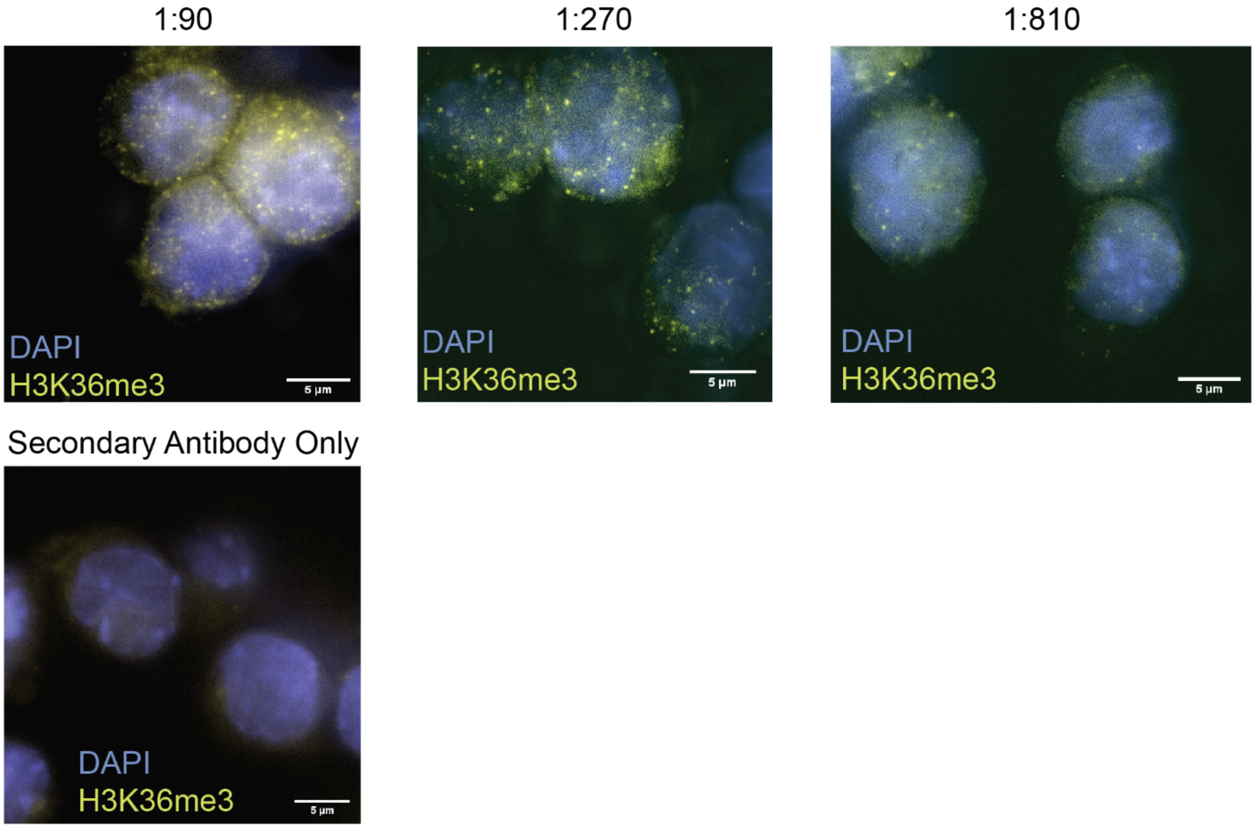


**Supplementary Fig. 35. Dilution of immunofluorescence antibody allows for single spot detection**. Example images of a single slice of F1 2-1 mESCs with the H3K36me3 antibody at different dilutions. DAPI is in blue, and signal from the secondary antibody is in yellow. The concentration used in experiments was 1:270.

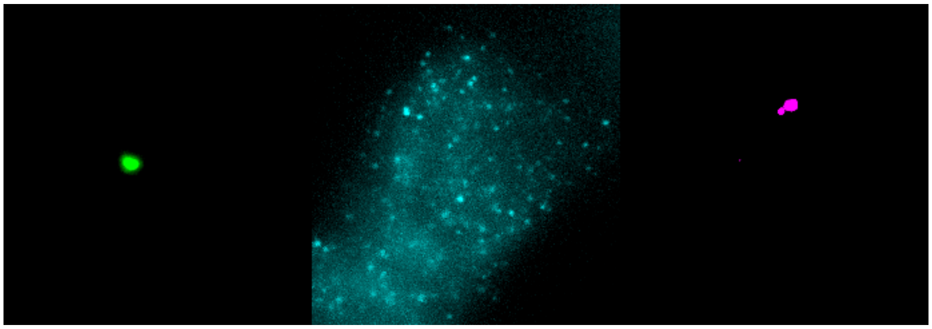

**Supplementary Fig. 36. Same image as Figure 6a but as individual channels.** (left) Xist intronic signal, (middle) H3K36me3, (right) Tsix 5’ intronic probes. mESC after 2 days of differentiation.

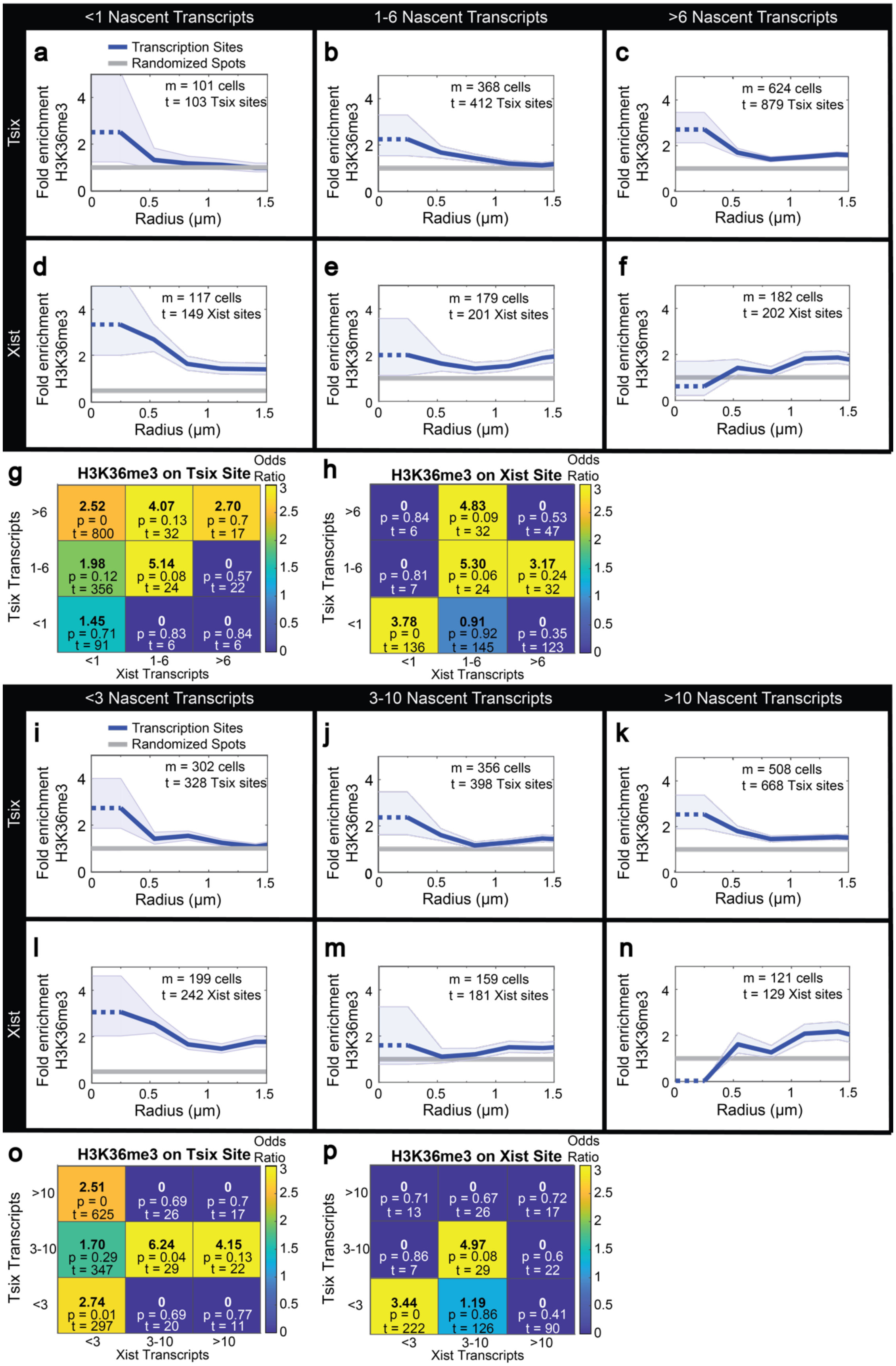

<1 Nascent Transcripts
1-6 Nascent Transcripts
>6 Nascent Transcripts
Tsix
Xist
a
b
c
d
e
f
Transcription Sites
Randomized Spots
Fold enrichment H3K36me3
Radius (μm)
m = 101 cells
t = 103 Tsix sites
m = 368 cells
t = 412 Tsix sites
m = 624 cells
t = 879 Tsix sites
m = 117 cells
t = 149 Xist sites
m = 179 cells
t = 201 Xist sites
m = 182 cells
t = 202 Xist sites
g
H3K36me3 on Tsix Site
Odds Ratio
Tsix Transcripts
Xist Transcripts
>6
1-6
<1
2.52 p = 0 t = 800
4.07 p = 0.13 t = 32
2.70 p = 0.7 t = 17
1.98 p = 0.12 t = 356
5.14 p = 0.08 t = 24
0 p = 0.57 t = 22
1.45 p = 0.71 t = 91
0 p = 0.83 t = 6
0 p = 0.84 t = 6
h
H3K36me3 on Xist Site
0 p = 0.84 t = 6
4.83 p = 0.09 t = 32
0 p = 0.53 t = 47
0 p = 0.81 t = 7
5.30 p = 0.06 t = 24
3.17 p = 0.24 t = 32
3.78 p = 0 t = 136
0.91 p = 0.92 t = 145
0 p = 0.35 t = 123
<3 Nascent Transcripts
3-10 Nascent Transcripts
>10 Nascent Transcripts
i
j
k
l
m
n
m = 302 cells
t = 328 Tsix sites
m = 356 cells
t = 398 Tsix sites
m = 508 cells
t = 668 Tsix sites
m = 199 cells
t = 242 Xist sites
m = 159 cells
t = 181 Xist sites
m = 121 cells
t = 129 Xist sites
o
H3K36me3 on Tsix Site
>10
3-10
<3
2.51 p = 0 t = 625
0 p = 0.69 t = 26
0 p = 0.7 t = 17
1.70 p = 0.29 t = 347
6.24 p = 0.04 t = 29
4.15 p = 0.13 t = 22
2.74 p = 0.01 t = 297
0 p = 0.69 t = 20
0 p = 0.77 t = 11
p
H3K36me3 on Xist Site
0 p = 0.71 t = 13
0 p = 0.67 t = 26
0 p = 0.72 t = 17
0 p = 0.86 t = 7
4.97 p = 0.08 t = 29
0 p = 0.6 t = 22
3.44 p = 0 t = 222
1.19 p = 0.86 t = 126
0 p = 0.41 t = 90

**Supplementary Fig. 37. Quantification of Colocalization at Sites of Transcription (tsites) with histone modification signal H3K36me3 at Different Cutoffs. a-p** Show the same data as in Figure 6d-k but with different thresholds. **a-h** panels show a lower cutoff for low (<1 transcript), medium (2 to 6 transcripts), and high (>6 transcripts) nascent expression. The results are independent of the cutoffs. Xist and Tsix are labeled with intronic probes. **i-p** panels show a higher cutoff for low (<3 transcripts), medium (3 to 10 transcripts), and high (>10 transcripts) nascent expression.

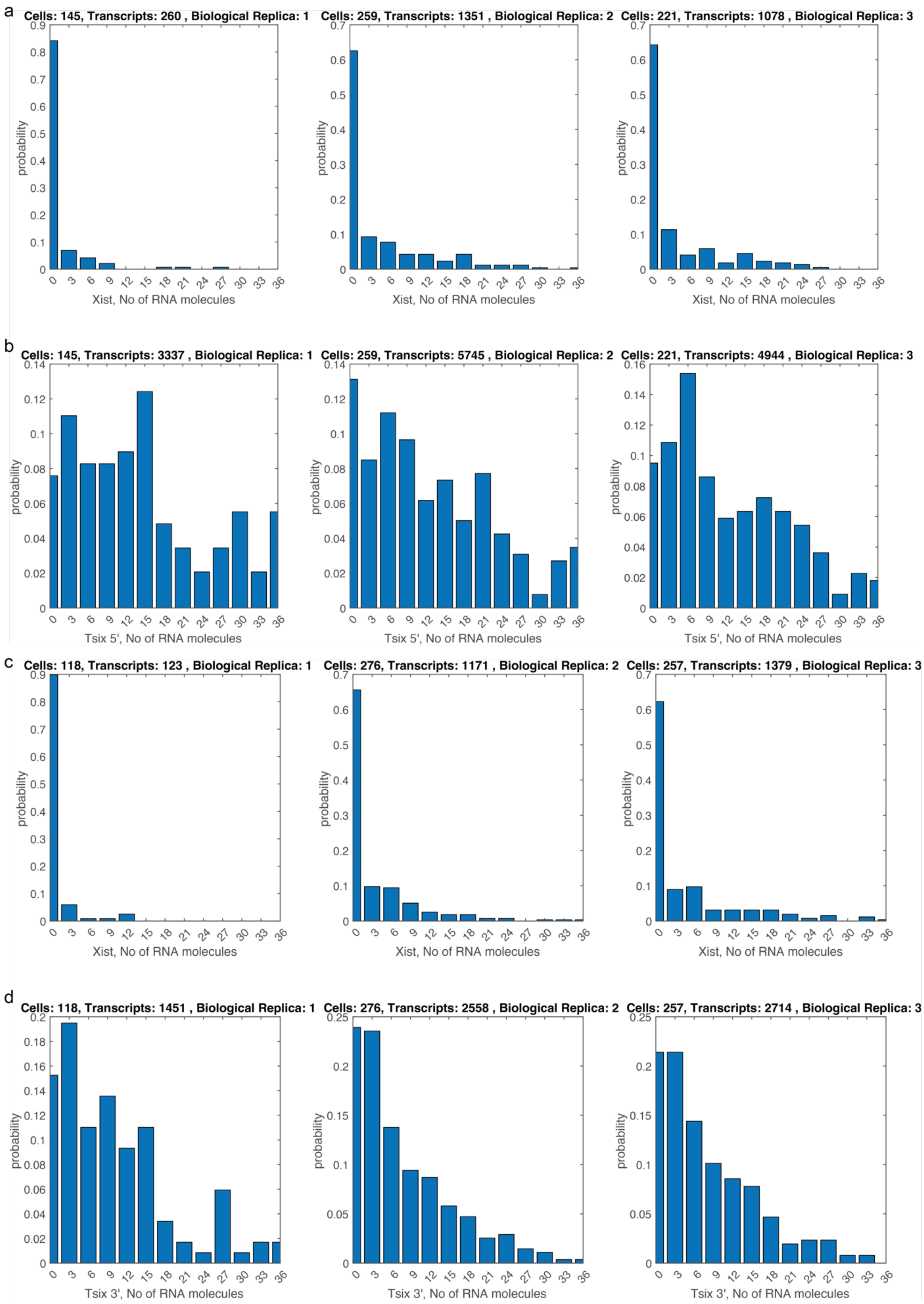
a
Cells: 145, Transcripts: 260 , Biological Replica: 1
Cells: 259, Transcripts: 1351 , Biological Replica: 2
Cells: 221, Transcripts: 1078 , Biological Replica: 3
probability
Xist, No of RNA molecules
b
Cells: 145, Transcripts: 3337 , Biological Replica: 1
Cells: 259, Transcripts: 5745 , Biological Replica: 2
Cells: 221, Transcripts: 4944 , Biological Replica: 3
Tsix 5', No of RNA molecules
c
Cells: 118, Transcripts: 123 , Biological Replica: 1
Cells: 276, Transcripts: 1171 , Biological Replica: 2
Cells: 257, Transcripts: 1379 , Biological Replica: 3
Xist, No of RNA molecules
d
Cells: 118, Transcripts: 1451 , Biological Replica: 1
Cells: 276, Transcripts: 2558 , Biological Replica: 2
Cells: 257, Transcripts: 2714 , Biological Replica: 3
Tsix 3', No of RNA molecules

**Supplementary Fig. 38. Detailed distributions from Figure 5.** Marginal probability distributions for each of the replica experiments of nascent intronic Xist (A, C),  Tsix 5 '(B), and Tsix 3 ' (D).

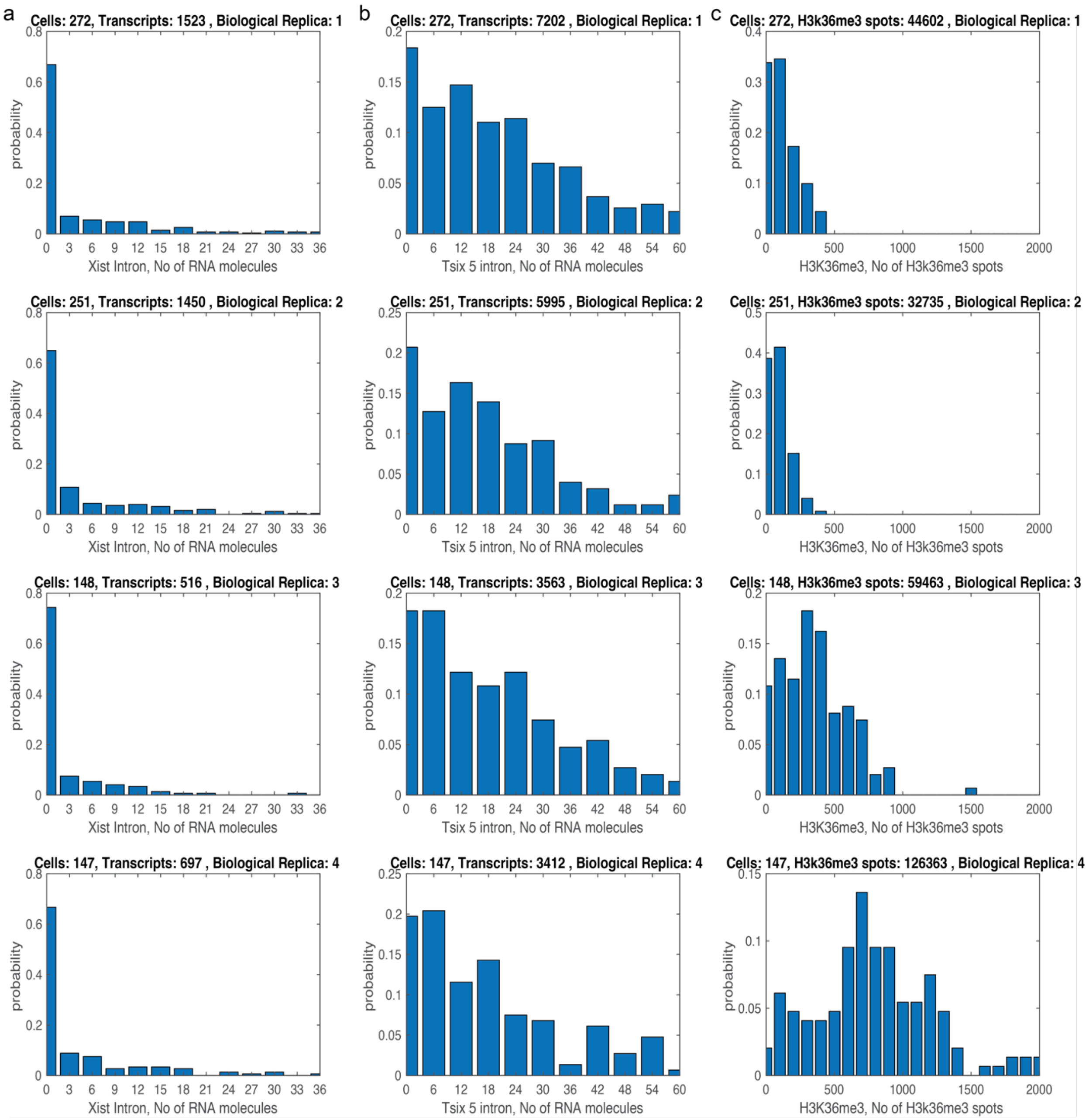


**Supplementary Fig. 39. Detailed distributions from Figure 6. a-c** Marginal probability distributions for each of the replica experiments of nascent intronic Xist (a), nascent intronic Tsix 5' (b), and H3K36me3 (c).

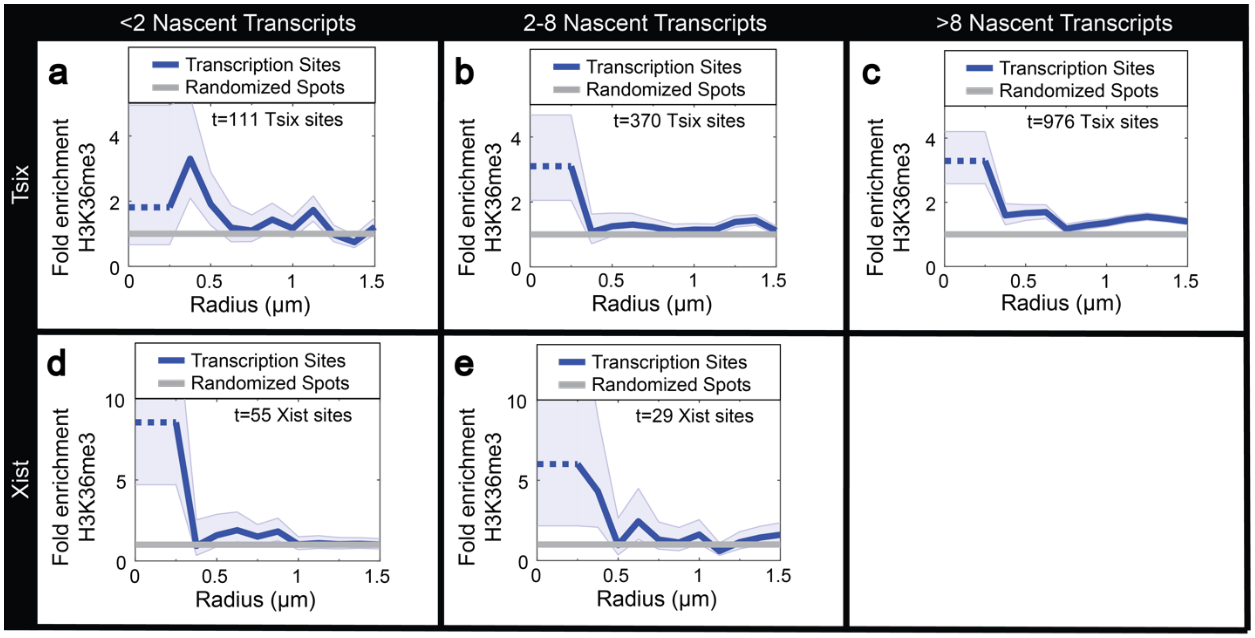


**Supplementary Fig. 40. Day 0 H3K36me3 enrichment**. Quantification of cells at day 0 of differentiation using the approach described in (Figure 6c). Fold enrichment of H3K36me3 calculated by Cochran–Mantel–Haenszel statistics with real transcription sites (blue) relative to random (gray). Shaded error bars are approximately one standard deviation. N = 3 biological replica experiments, n = 791 total cells. In each subpanel, t is the number of transcription sites with the designated number of nascent transcripts, t is the number of transcription sites with the number of molecules specified in each panel. **a-c** Fold enrichment of H3K36me3 at sites with low, medium, and high Tsix transcription. **d-e** Fold enrichment of H3K36me3 at sites with low and medium Xist transcription. Xist and Tsix are labeled with intronic probes.

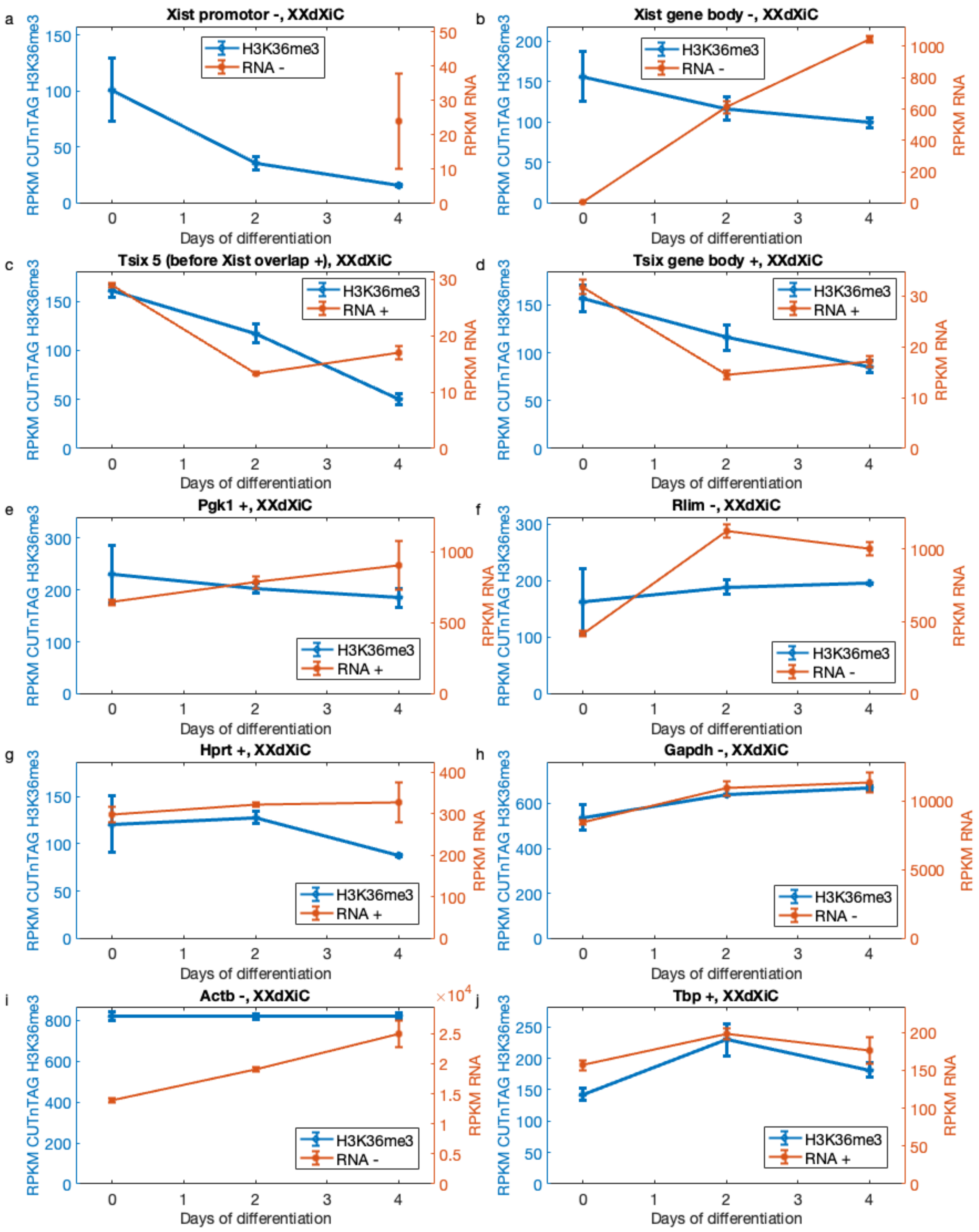


**Supplementary Fig. 41. Allele-specific re-analysis of CUT & TAG of H3K36me3 and strand specific RNA-seq in mouse embryonic stem cells (mESC) during differentiation (day 0, 2 ,4)**

**in the XXΔXic genotype published in [65].** H3K36me3 and RNA-seq read density in RPKM for **a** Xist promotor (Chr X:103483234-103486633), **b** Xist gene body (Chr X, 103460374-103483233), **c** Tsix 5' before Xist (Chr X, 103431518-103460373), **d** Tsix gene body (Chr X, 103431518-103484957), **e** Pgk1 gene body (Chr X, 106185101-106205699), **f** Rlim gene body (Chr X, 103957163-103981285), **g** Hprt gene body (Chr X, 52988078-53021660), **h** Gapdh gene body (Chr 6:125161852-125166467), **i** Actb gene body (Chr 5:142903115-142906754), **j** Tbp gene body (chr17:15499888-15517427). 'RNA +' represents the plus strand, and 'RNA –' represents the minus strand. Statistical analysis details are provided in Supplementary Table 2.

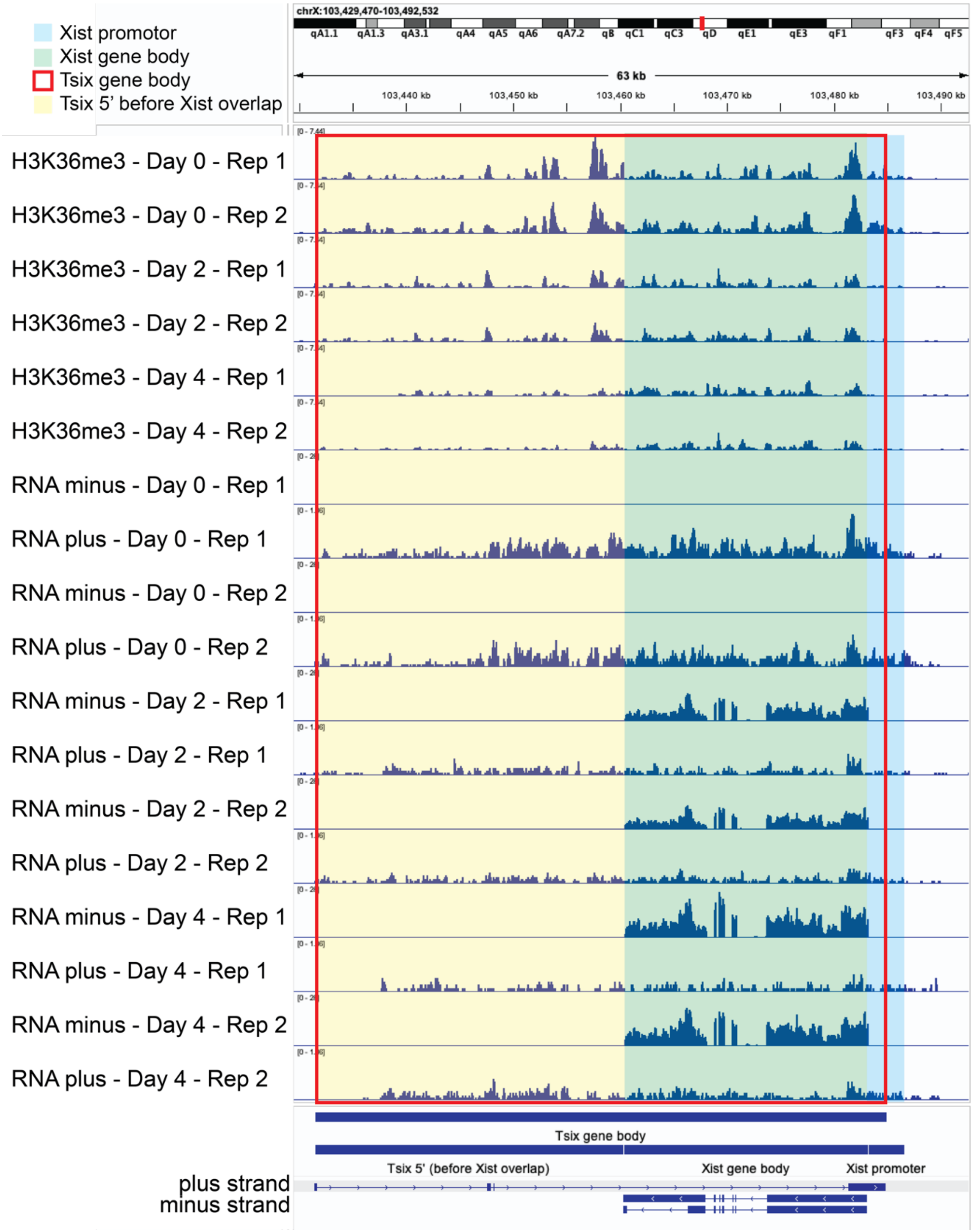


**Supplementary Fig. 42. IGV density plots of CUT & TAG H3K36me3 and strand specific RNA-seq for Xist and Tsix in the XXΔXic genotype published in [65].** Shaded area is used for quantification in Supplementary Fig. 41. Xist promotor (light blue), Xist gene body (light green), Tsix gene body (red line), and Tsix 5' before Xist overlap (light yellow). H3K36me3, RNA plus and RNA minus strand tracks are each grouped to the same height.

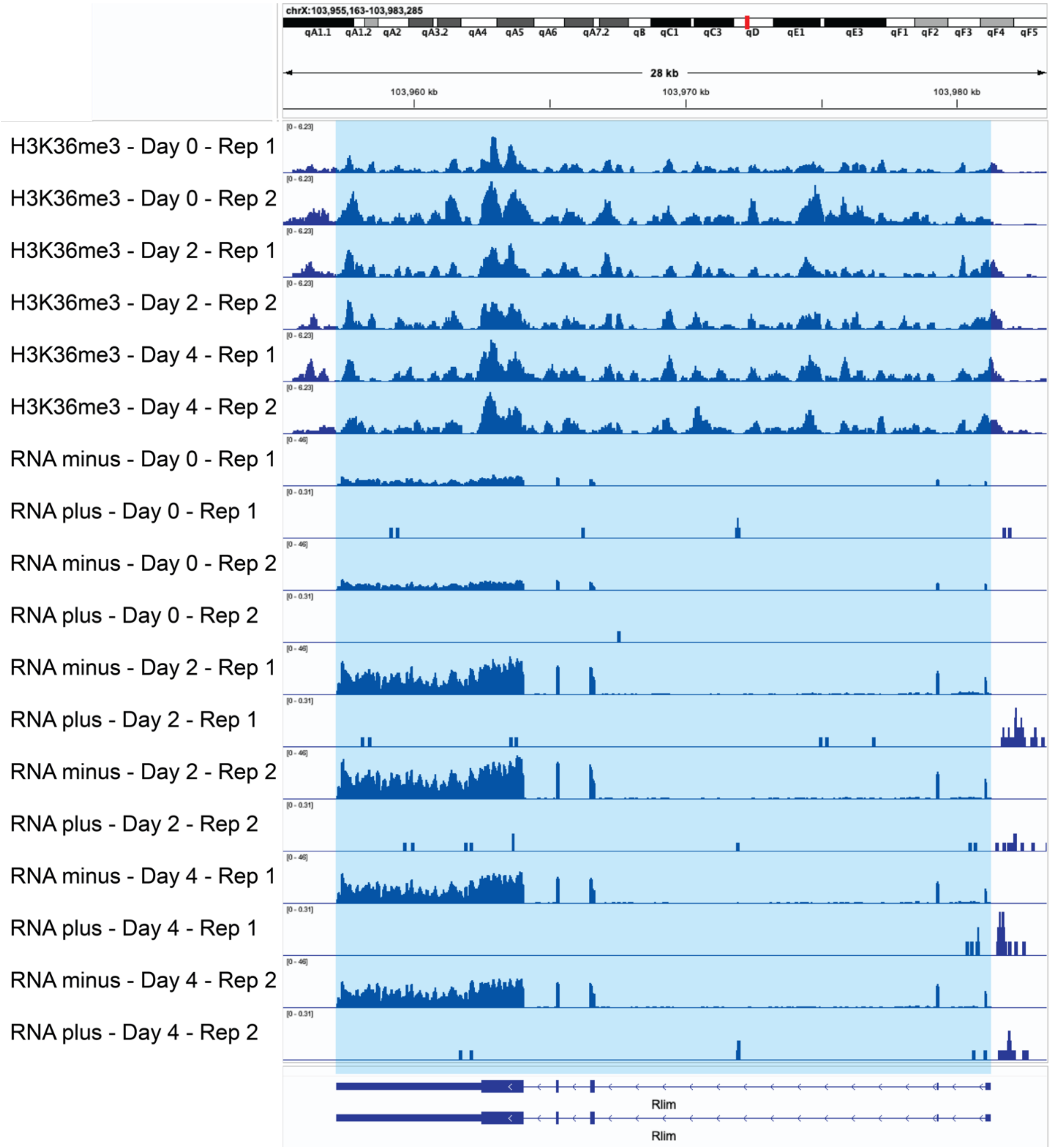


**Supplementary Fig. 43. IGV density plots of CUT & TAG H3K36me3 and strand specific RNA-seq for Rlim in the XXΔXic genotype published in [65].** Blue shaded area is used for quantification in Supplementary Fig. 41. H3K36me3, RNA plus and RNA minus strand tracks are each grouped to the same height.

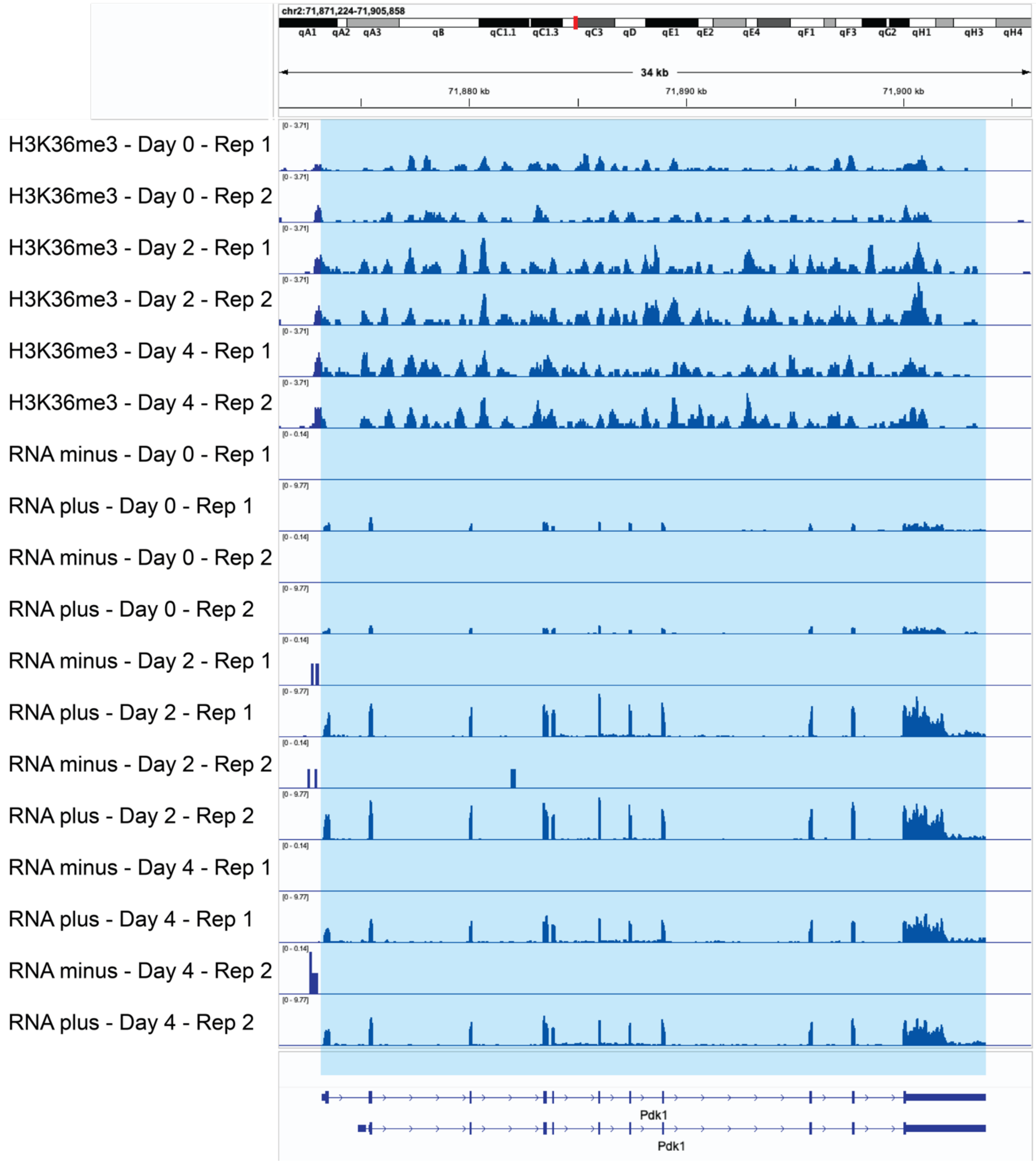


**Supplementary Fig. 44. IGV density plots of CUT & TAG H3K36me3 and strand specific RNA-seq for Pdk1 in the XXΔXic genotype published in [65].** Blue shaded area is used for quantification in Supplementary Fig. 41. H3K36me3, RNA plus and RNA minus strand tracks are each grouped to the same height.

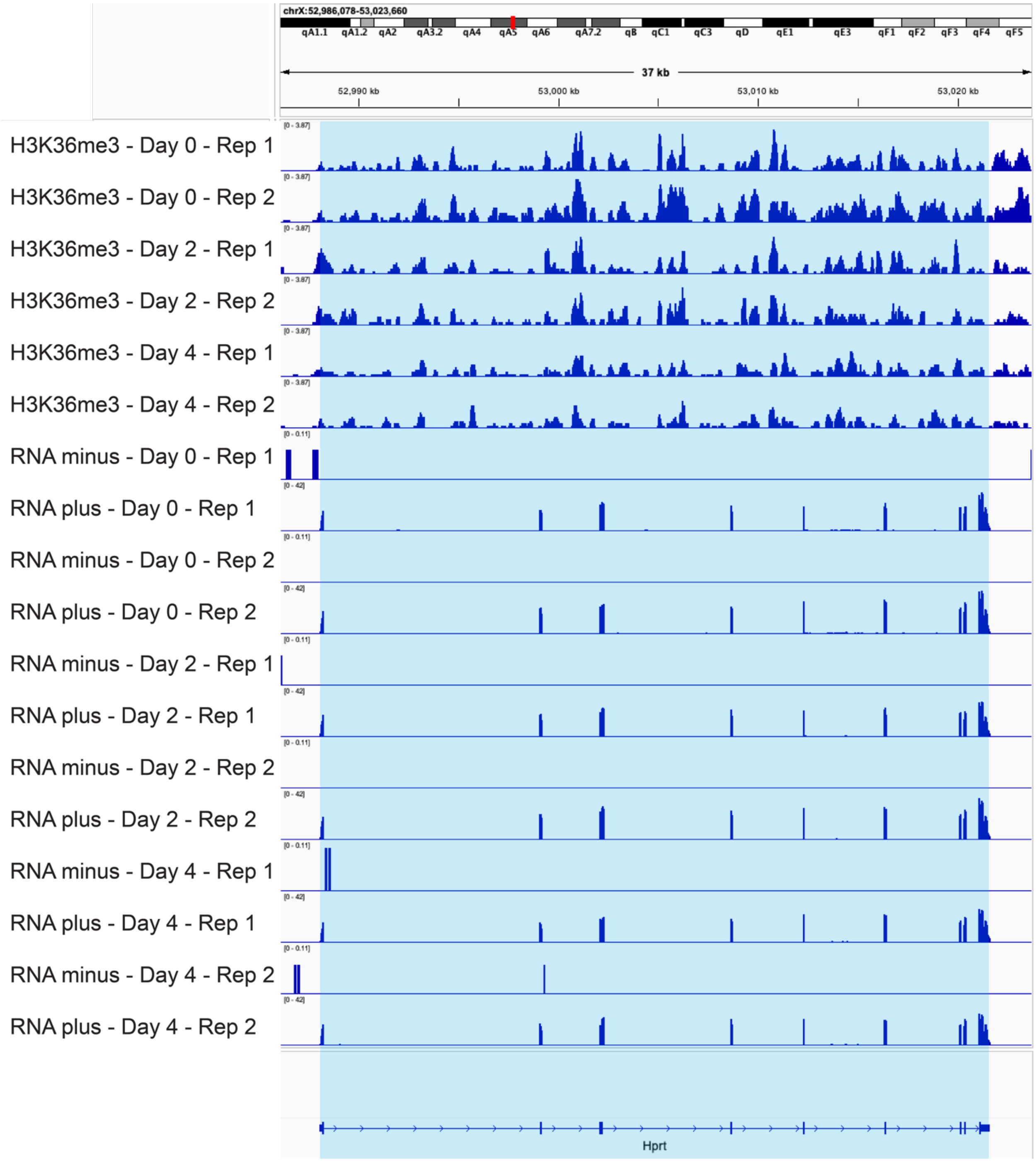


**Supplementary Fig. 45. IGV density plots of CUT & TAG H3K36me3 and strand specific RNA-seq for Hprt in the XXΔXic genotype published in [65].** Blue shaded area is used for quantification in Supplementary Fig. 41. H3K36me3, RNA plus and RNA minus strand tracks are each grouped to the same height.

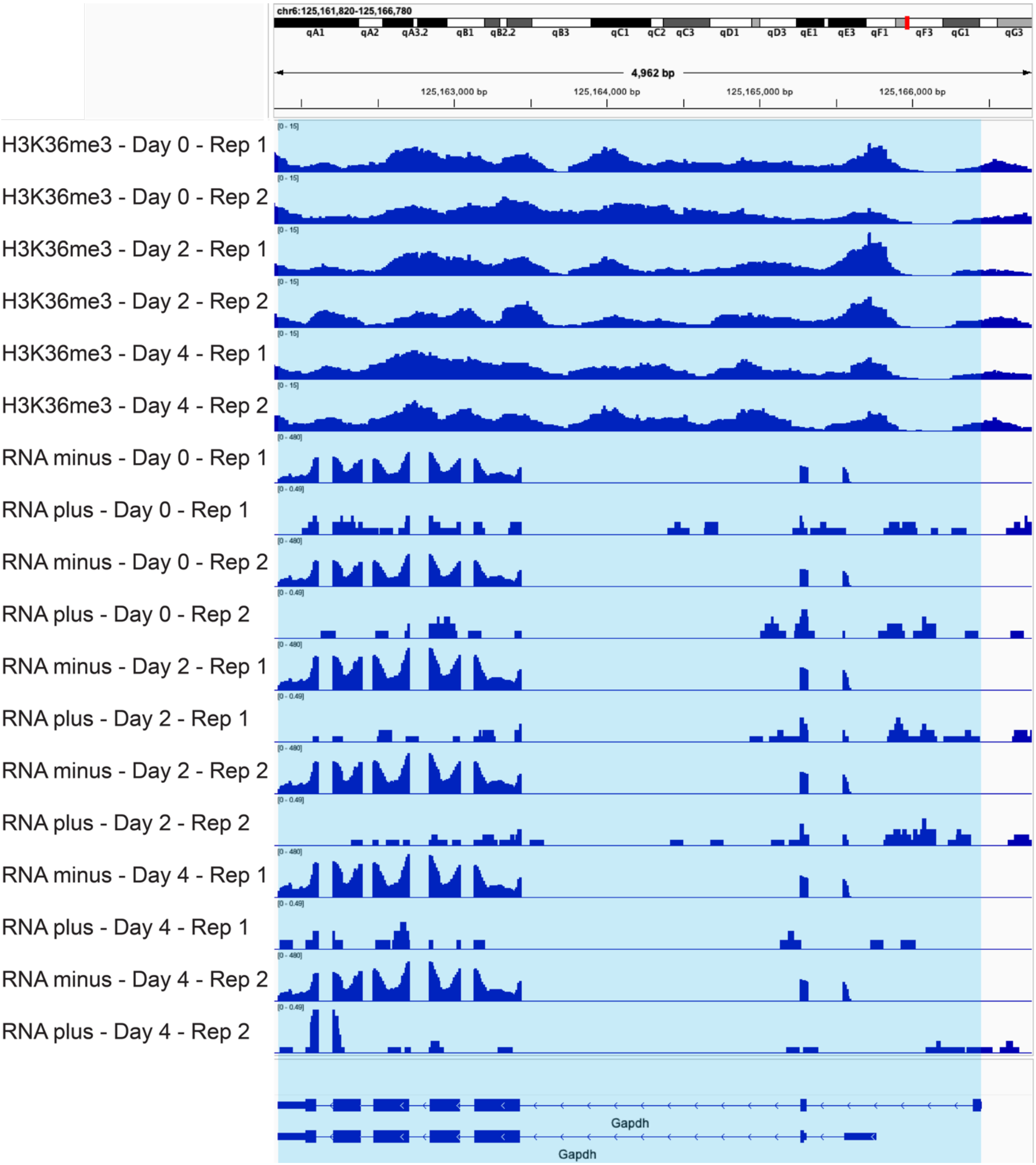


**Supplementary Fig. 46. IGV density plots of CUT & TAG H3K36me3 and strand specific RNA-seq for Gapdh in the XXΔXic genotype published in [65].** Blue shaded area is used for quantification in Supplementary Fig. 41. H3K36me3, RNA plus and RNA minus strand tracks are each grouped to the same height.

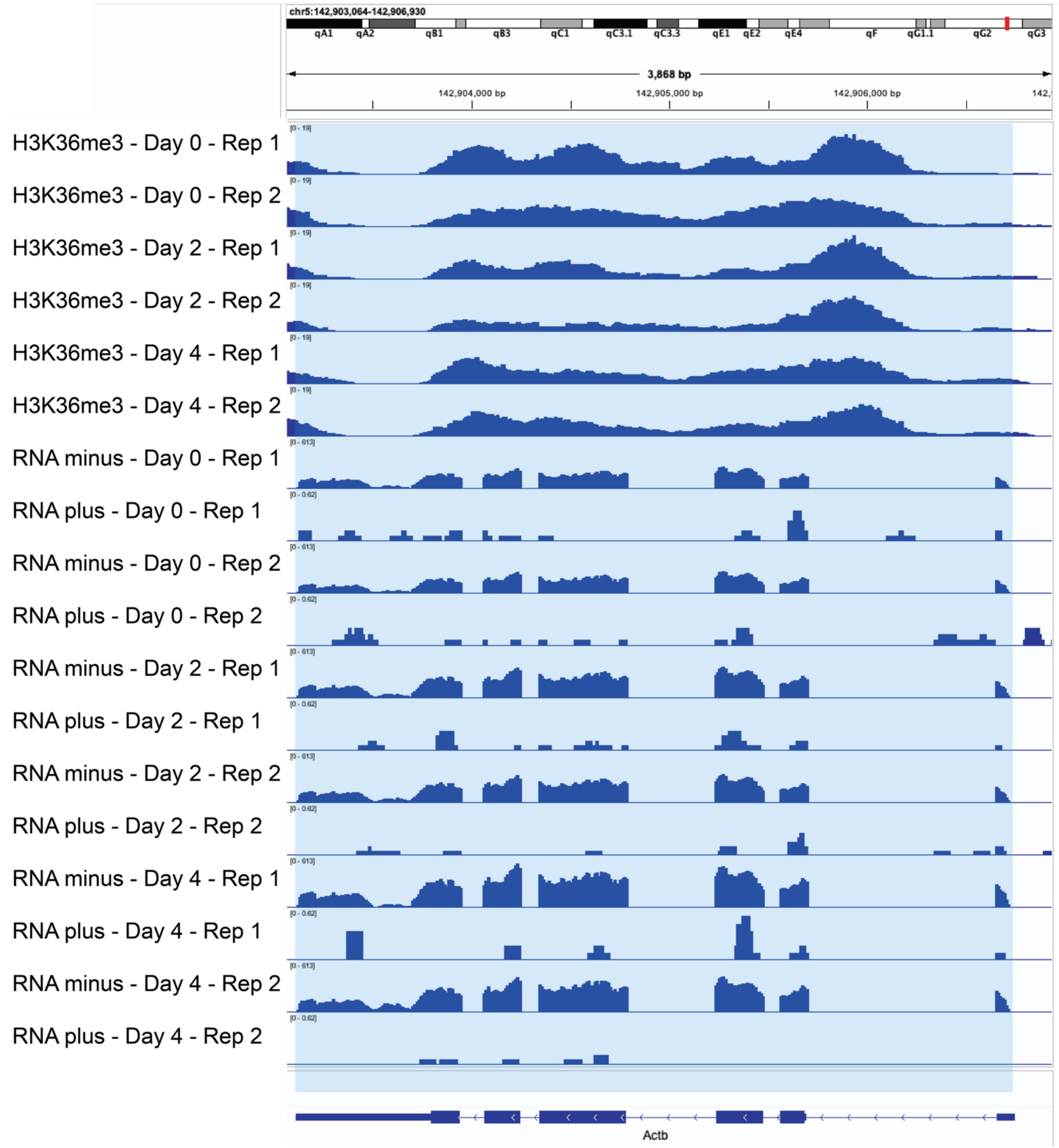


**Supplementary Fig. 47. IGV density plots of CUT & TAG H3K36me3 and strand specific RNA-seq for Actb in the XXΔXic genotype published in [65].** Blue shaded area is used for quantification in Supplementary Fig. 41. H3K36me3, RNA plus and RNA minus strand tracks are each grouped to the same height.

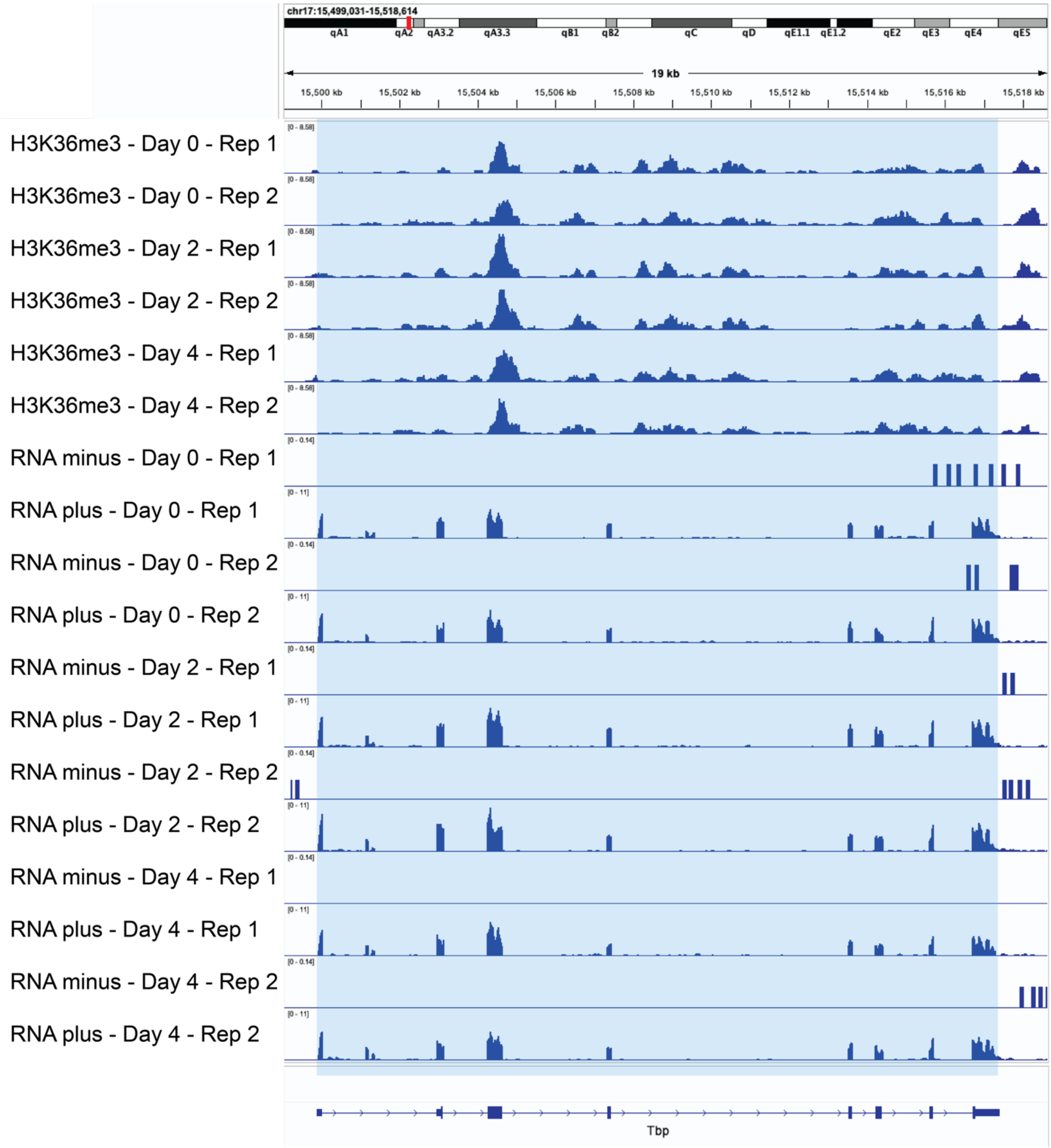


**Supplementary Fig. 48. IGV density plots of CUT & TAG H3K36me3 and strand specific RNA-seq for Tbp in the XXΔXic genotype published in [65].** Blue shaded area is used for quantification in Supplementary Fig. 41. H3K36me3, RNA plus and RNA minus strand tracks are each grouped to the same height.

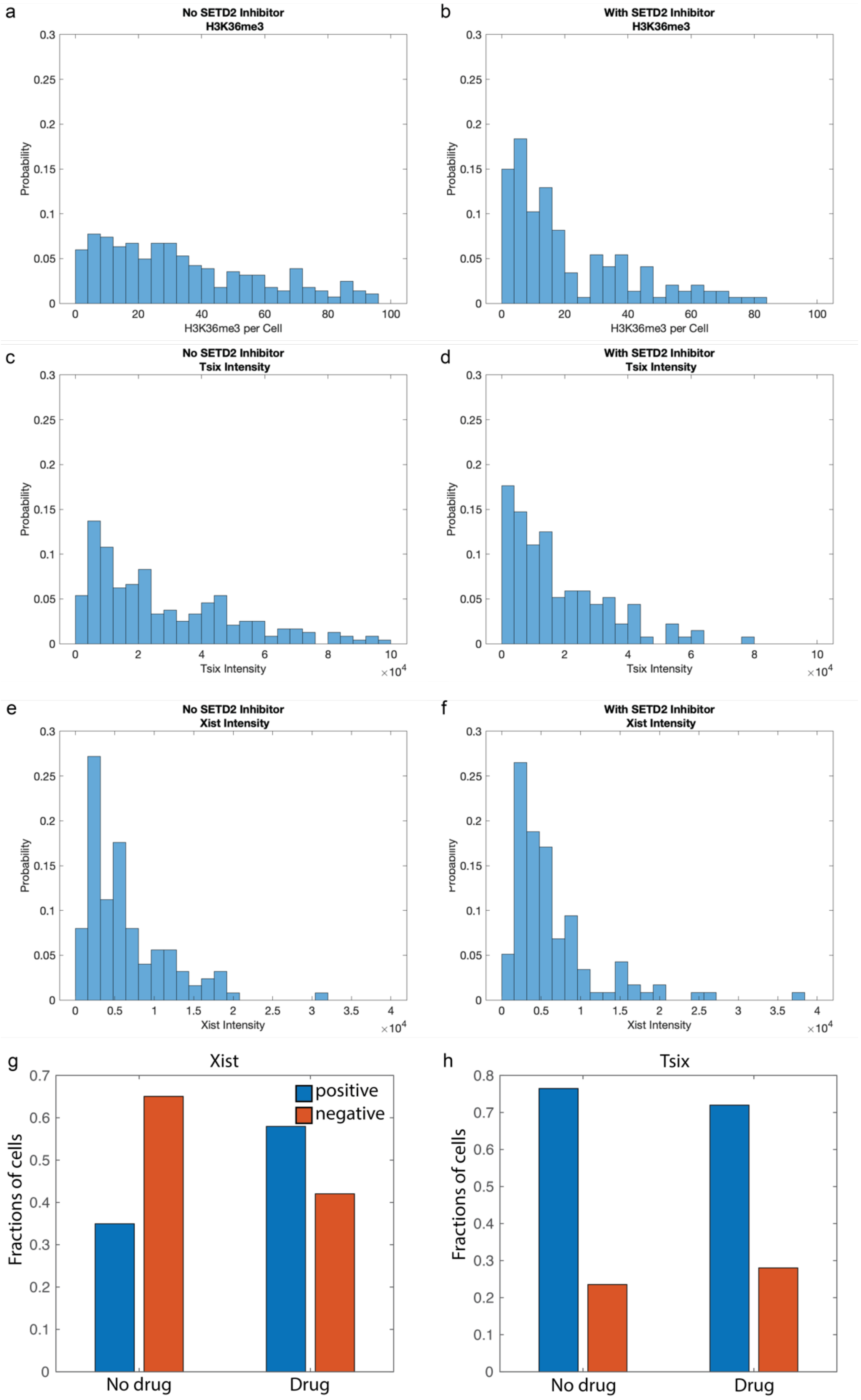
a
No SETD2 Inhibitor
H3K36me3
Probability
H3K36me3 per Cell
b
With SETD2 Inhibitor
H3K36me3
Probability
H3K36me3 per Cell
c
No SETD2 Inhibitor
Tsix Intensity
Probability
Tsix Intensity
d
With SETD2 Inhibitor
Tsix Intensity
Probability
Tsix Intensity
e
No SETD2 Inhibitor
Xist Intensity
Probability
Xist Intensity
f
With SETD2 Inhibitor
Xist Intensity
Xist Intensity
g
Xist
positive
negative
Fractions of cells
No drug
Drug
h
Tsix
Fractions of cells
No drug
Drug

**Supplementary Fig. 49.** Setd2 inhibition changes number of H3K36me3 marks, Tsix transcription site intensities and, and fraction of Xist expressing cells after two days of differentiation. **a-b** Number of H3K36me3 spots per cell in normal (a) and SETD2 inhibited (b) cells changes (KS-test). **c-d** Tsix nascent transcription site intensities in normal (c) and SETD2 inhibited (d) cells changes. **e-f** Xist nascent transcription site intensities in normal (e) and SETD2 inhibited (f) cells does not change. **g-h** Fraction of cells with Xist (g) and Tsix (h) nascent transcription sites in normal (no drug) and SETD2 inhibited cells. The fraction of cell with Xist nascent transcription change whereas the fraction of cell with Tsix nascent transcription does not changes.

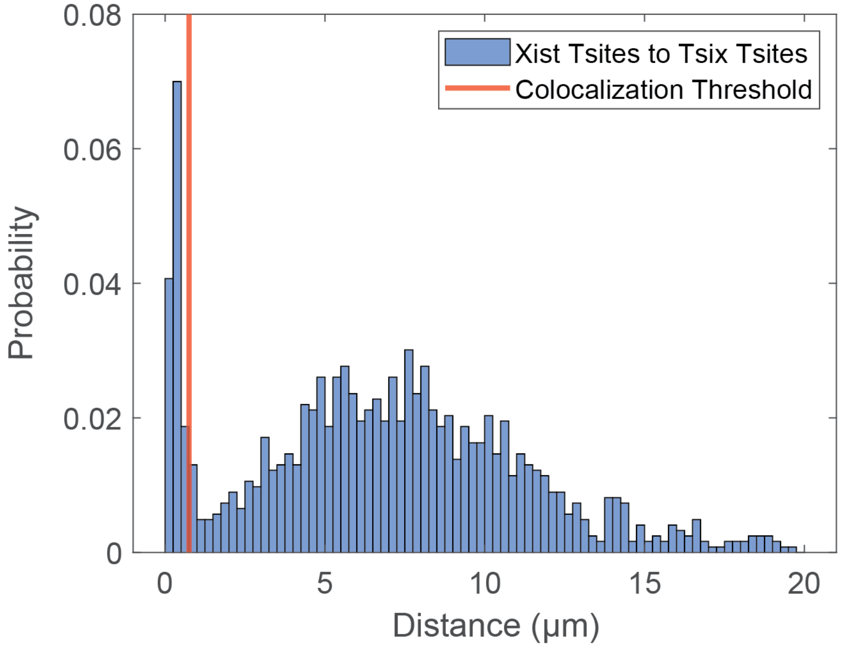


**Supplementary Fig. 50. Distances between Xist and Tsix Transcription Sites**. Histograms of the distances between Xist (labelled with Xist Intron probes) and Tsix (labelled with Tsix 5' Intron probes) transcription sites (tsites) at day 2 of differentiation show a bimodal distribution. Colocalization events at low distances correspond to co-transcription on the same allele, while the higher distances correspond to transcription on different alleles. N = 4 biological replicas, n = 818 total cells, m = 406 cells with both Xist and Tsix tsites, d = 1229 total distances calculated.

| **Antigen** | **Host / Isotype** | **Coupled Fluorophore** | **Manufacturer** | **Catalog Number** | **Dilution** |
|---|---|---|---|---|---|
| H3K36me3 | Rabbit | none | CS | 4909S | 1:270 |
| Anti-Rabbit | Goat | Alexa Fluor® 488 | CS | 4412S | 1:2000 |

**Supplementary Table 1. Antibodies and dilutions**. The antigen, host, fluorophore, manufacturer, Catalog number, and dilution are listed.
